# The network structure of hematopoietic cancers

**DOI:** 10.1101/2022.11.25.517762

**Authors:** Arturo Kenzuke Nakamura-García, Jesús Espinal-Enríquez

## Abstract

Hematopoietic cancers (HCs) are a heterogeneous group of malignancies that affect blood, bone marrow and lymphatic system. Here, by analyzing 1,960 RNA-Seq samples from three independent datasets, we explored the co-expression landscape in HCs, by inferring gene co-expression networks (GCNs) with four cancer phenotypes (B and T-cell acute leukemia -BALL, TALL-, acute myeloid leukemia -AML-, and multiple myeloma -MM-) as well as non-cancer bone marrow. We characterized their structure (topological features) and function (enrichment analyses). We found that, as in other types of cancer, the highest co-expression interactions are intra-chromosomal, which is not the case for control GCNs. We also detected a highly co-expressed group of overexpressed pseudogenes in HC networks. The four GCNs present only a small fraction of common interactions, related to canonical functions, like immune response or erythrocyte differentiation. With this approach, we were able to reveal cancer-specific features useful for detection of disease manifestations.

**Significance:** We demonstrate that gene co-expression is deregulated in four HC, observed by an elevated proportion of intrachromosome interactions in their GCNs with respect to their normal counterparts, and increased interactions between pseudogenes (more evident in AML). This deregulation might be associated with the age of the patients.

## Introduction

Hematological cancers (HC) are a heterogeneous group of malignancies from immune cells or their precursors. They can be categorized into three major groups: leukemias, lymphomas and multiple myeloma (1). Leukemias are malignancies from precursors of leukocytes that accumulate in blood or in the bone marrow. They can arise from myeloid or lymphoid linage. According to the velocity with which leukemias progress they can be acute, with a fast development, or chronic, if the disease progresses slowly. Lymphomas are tumors from lymphocytes that affect the lymph nodes, thymus or spleen, and can be broadly categorized into Hodgkin and Non-Hodgkin lymphomas. Multiple myeloma is a tumor of differentiated plasma cells, which unlike normal plasma cells, can proliferate and accumulate in the bone marrow. It is also characterized by the secretion of large amounts of a monoclonal immunoglobulin protein.

The malignant transformation of normal cells entails alterations in their regulatory circuits and molecular processes that have consequences at multiple scales of the organism. The interactions between the different components in each of these scales makes difficult to fully understand the global deregulation that produce and sustains the cancerous phenotype. Contemporary biology has developed computational approaches, able to analyze large data from next generation sequencing (NGS) technologies. These implementations have proven to be useful to unveil cancer-associated features, as well as structural or functional differences with normal tissues (2).

Next generation sequencing, in particular gene expression profiles, have been broadly used to discover key features that may alter the transcriptional program, triggering different outcomes (3, 4). Although the importance of genetic expression in cancer is out of discussion, it is also clear that the gene regulation during the carcinogenic process is strongly altered by several factors. Additionally, the gene expression landscape often does not provide information on how those genes are regulated. (5).

To overcome the latter challenge, a common approach used for high-throughout-derived datasets, is the gene co-expression network (GCN). These networks are commonly inferred by correlating the expression profile of gene couples within multiple samples. GCNs offer a framework that allows the analysis of global changes in a given phenotype, such as cancer. With this approach, it can be quantified the statistical dependence of any pair of genes’ expression (6–9).

In a nutshell, GCNs are theoretical objects composed of genes, which correspond to nodes in the network. Those genes are connected to other genes by means of a given kind of correlation between them. To analyze cancer NGS-derived data, GCNs have become one of the most broadly used tools, given that several implementations are freely available, the obtained results are reproducible and they can be easily contrasted with the phenotypic counterpart (10).

Previous works from our group have used GCNs to analyze the changes on the transcriptional regulation profiles from several carcinomas and their effects on the regulation of biological processes (BPs) in the disease (11–15). From these analysis we have identified two main common phenomena between the different cancer GCNs: 1) an elevated proportion of strong interactions between genes from the same chromosomes (intra-chromosomal interactions) in comparison to the network of their respective normal tissue, and 2) the size of cancer-derived networks connected components is much smaller than those observed in a normal-derived GCN. We refer to this phenomenon as loss of inter-chromosomal co-expression and it has been observed in each cancer GCN that we have analyzed (16).

To the extent of our knowledge, no analysis using this approach has been reported regarding hematological malignancies. Hence, we sought to construct GCNs of these diseases employing RNA-Seq data of 1,495 bone marrow samples from The Cancer Genome Atlas consortium (TCGA). We analyzed samples from patients diagnosed with acute lymphocytic leukemia from B and T cells (BALL (*n* = 210) and TALL (*n* = 245)), acute myeloid leukemia (AML (*n* = 159)), multiple myeloma (MM (*n* = 844)) and samples of normal bone marrow (*n* = 37).

We inferred GCNs by using Mutual Information as a measure of statistical dependency between gene couples. In order to evaluate the robustness of the resulting networks’ features, we also calculated the Spearman correlation between all gene pairs for each phenotype. We performed a topological analysis on these individual networks, assessing the proportion of intra-chromosomal interactions in each of them and the proportions of the interactions between the different RNA biotypes available in the data sets.

Given that the four hematological cancers affect the same organ (bone marrow), we wondered whether the different cancer phenotypes had a common topological structure. To this purpose, we inferred the network of the shared interactions between the four HCs. Furthermore, we implemented a community detection algorithm on this intersection network and a functional enrichment analysis (FEA) on the resulting communities in order to identify the biological processes (BPs) associated to each community. A differential expression analysis of the genes present in those communities was also included in order to observe whether the BPs had a trend of differential gene expression, thus indicating an increase or depletion for such processes in a specific phenotype. Finally, we performed the same workflow of community detection and functional enrichment analysis on the individual HCs networks in order to determine additional functional features. This allowed us to identify the common BPs among the four HCs and compare them by the differential expression trend of the genes responsible of the enrichment of the communities.

For validation purposes, we also constructed HC networks from two different datasets: on from the Royal Children Hospital (RCH (*n* = 127 (17))), containing BALL-only samples; the other dataset was obtained from Saint Jude cloud (*n* = 465, for BALL (146), TALL (145), and AML (174) (18)).

## Results

### Cancer networks are topologically different to their normal counterparts

By looking at the networks’ architecture, we observe that the GCNs of the normal bone marrow are mainly composed of a single giant component, while the cancerous networks have several smaller components (Fig. 1A and Supplementary Material 1).

**Fig. 1.**
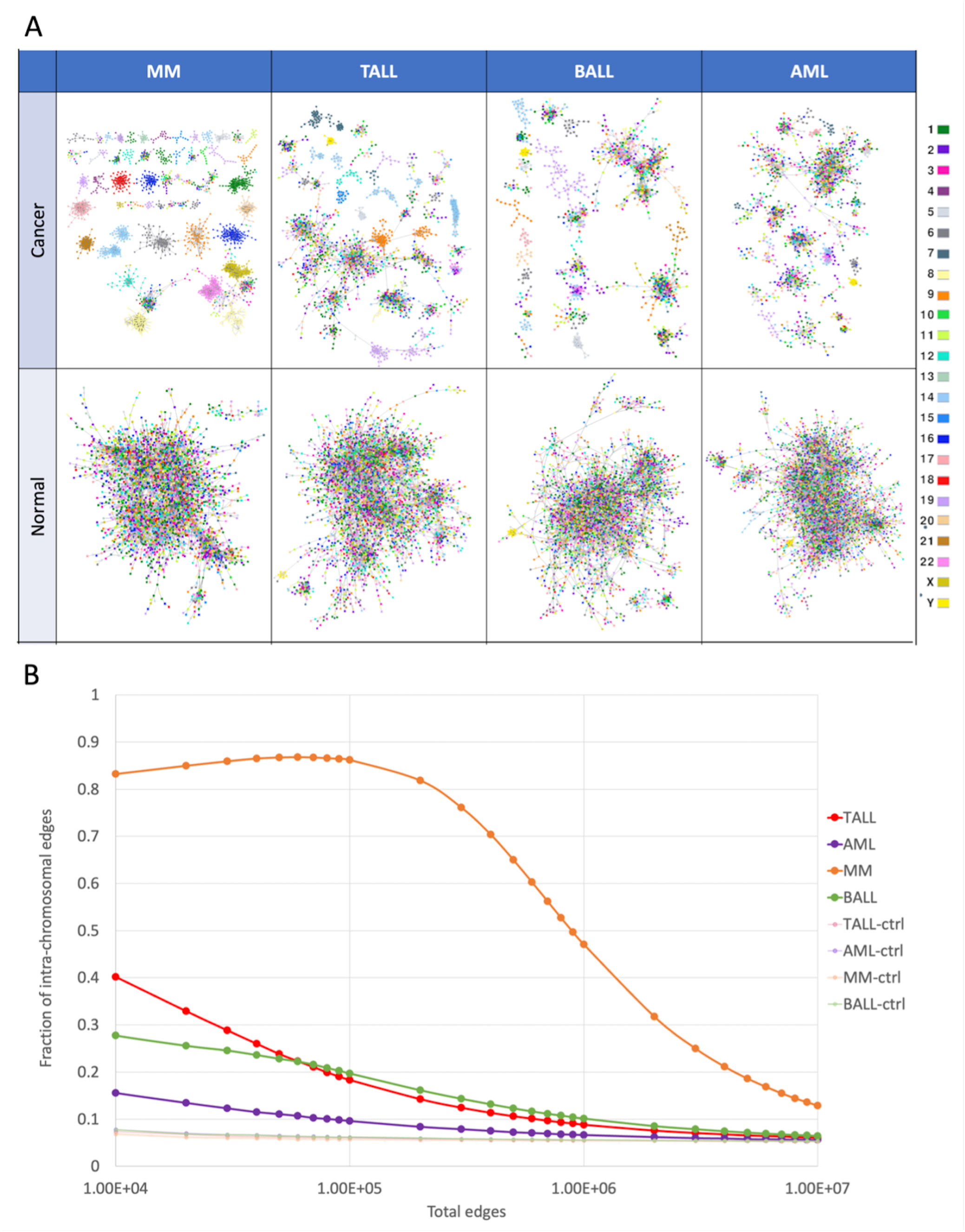
A) MI networks for the for phenotypes and their controls, painted by chromosome. B) Fraction of intrachromosomal (cis) interactions across different cut off values of the networks.

An aspect that we consider to be crucial in the topological analysis of the GCNs of cancer phenotypes is the regulation of inter-chromosomal interactions. As expected, several of the smaller components in our HCs networks are made up of genes from the same chromosome. This phenomenon is more clearly appreciated in MM, in which approximately 80% of the strongest interactions (at the cut-off point of 100,000) are intra-chromosome (Fig. 1B). TALL and BALL have a similar fraction of intra-chromosomal interactions between them, and AML has only a slight increase in these interactions in comparison with the normal bone marrow.

To assess whether the loss of inter-chromosomal interactions in HC networks was statistically significant when compared with the control GCNs, we performed a Kolmogorov-Smirnov (KS) test. Distribution of intra-chromosomal fractions in HCs and control phenotypes was compared by taking the values from top 10,000 to 1*e*^7^ interactions (Supplementary Material 2). KS tests indicate that the distributions of intra-chromosomal proportion in cancer and normal tissues are significantly different, with p-values ranging from 1*e*^*−*16^ to 1*e*^*−*5^, when the top 1 million interactions are considered. Therefore, in HCs, the strongest co-expressed gene pairs belong to the same chromosome, whereas for control GCNs, no biases towards high co-expression intra-chromosomal interactions were found.

### Cancer networks show elevated interactions between pseudogenes

To explore the co-expression between different types of genes other than protein coding, we did not apply any filter regarding RNA biotype. We constructed networks with all the available biotypes in the samples. In the AML network, more than 30% of the interactions occur between pseudogenes (green bar in Fig. 2A). This network has the least amount of intra-chromosome interactions of the four HCs. The opposite seems to happen in the MM network, with the least amount of interactions between pseudogenes and the biggest fractions of intra-chromosomal edges. Noteworthy, the MM patients are older than the other three HCs (Fig. 2B). The four HCs networks present an elevated fraction of interactions between pseudogenes in comparison with the normal BM networks.

**Fig. 2.**
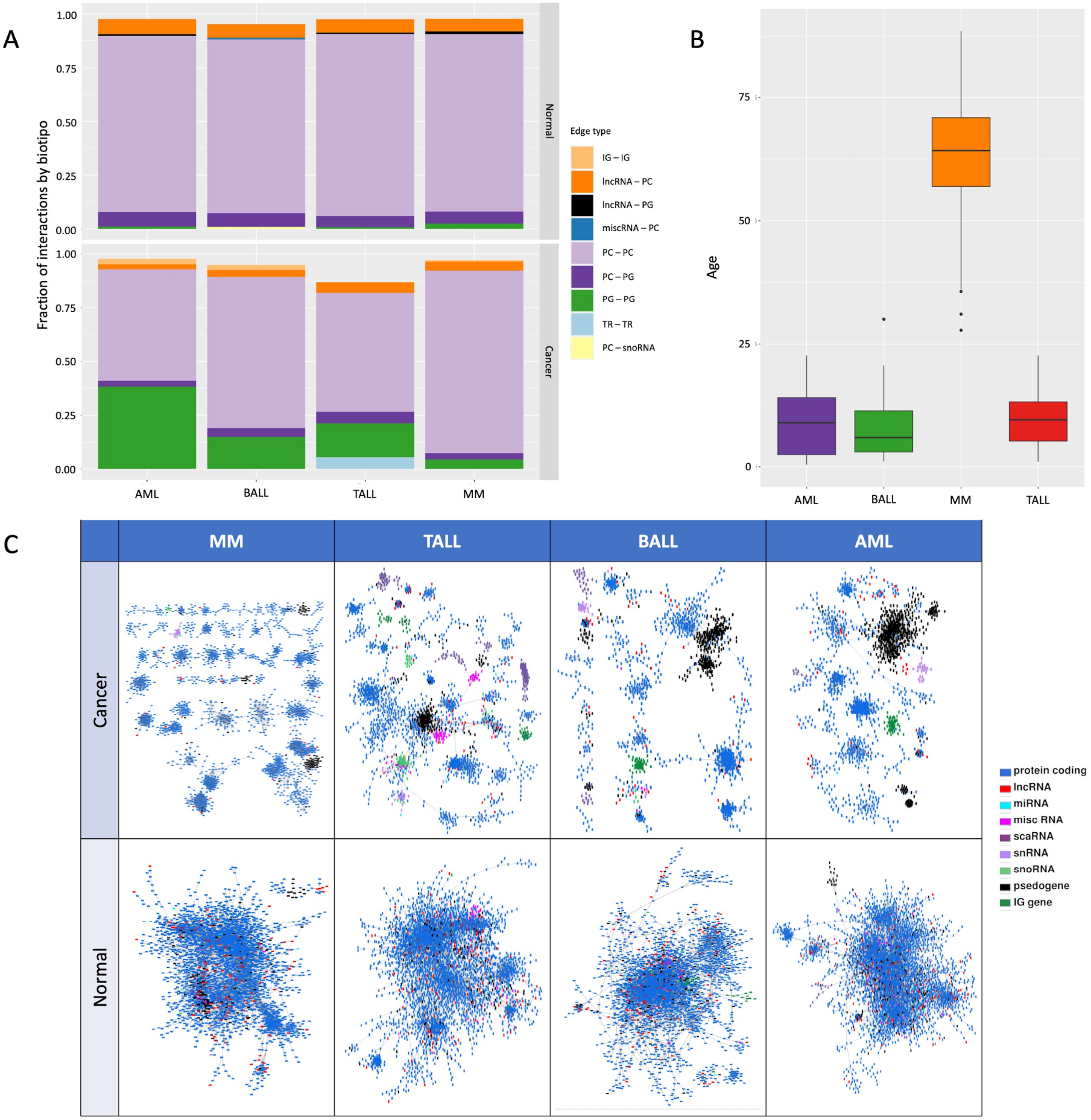
A) Fraction of Top 5 interacting biotypes on HC co-expression networks. Each color represents the most common type of interactions in the HC networks. IG = immunoglobulin; PC = protein coding; TR = T-cell receptor; PG = pseudogene. B) Distribution of ages per cancerous phenotype. C) MI networks for the four phenotypes and their controls, painted by the biotype.

Additionally, the co-expression pattern between pseudogenes in the cancer networks forms a cluster not observed in the normal BM networks (large black components in Fig. 2C). In the case of AML, the pseudogene component is the largest one of the entire network. These are pseudogenes of house-keeping genes, mostly of riboproteins and elongation factors.

So far, with the GCNs shown here we observe that 1) the gene co-expression landscape in HCs (as in carcinomas) is topologically different from normal GCNs, which is mainly manifested in the the loss of inter-chromosomal co-expression and the smaller connected components in HCs; and 2) that the co-expression interactions between pseudogenes is increased in the HCs networks, which is a feature that has not been explored by our group. As far as the authors are aware of, this has not been reported in the current literature.

### Differential expression analysis

A differential expression analysis was carried out to detect DEGs in each cancer phenotype in comparison the normal bone marrow group. Fig. 3 shows largest components of the four GCNs with the genes colored by its differential expression value. It is worth noticing that in the four cases, network components are predominantly formed by genes with the same differential expression trend. Table 1 shows the top five differentially expressed genes in BALL, TALL, MM, and AML, respectively. The complete differential expression analysis can be found in Supplementary Material 3.

**Fig. 3.**
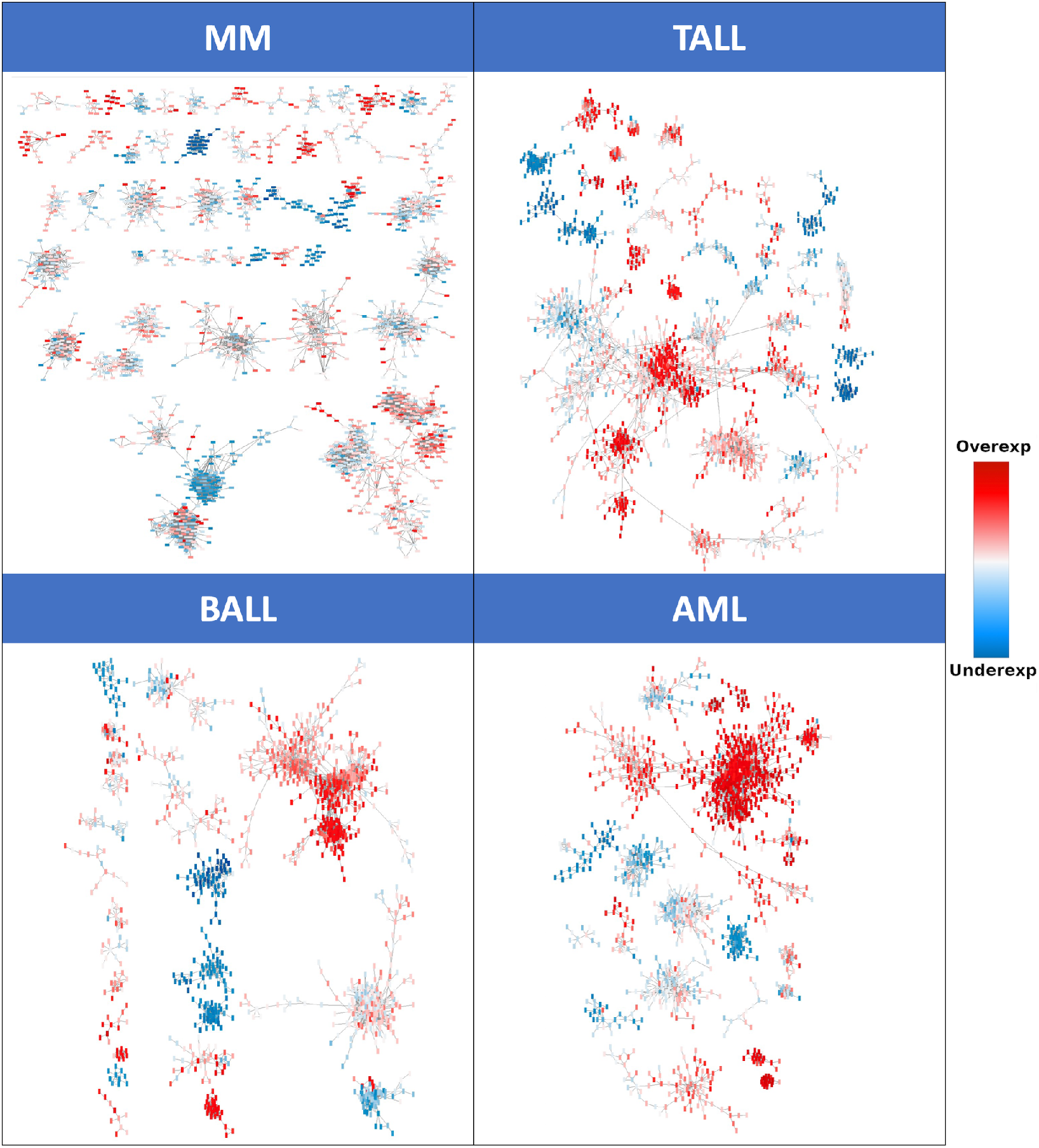
MI networks for the four cancer phenotypes painted by the differential expression values. Red nodes represent overexpression meanwhile underexpressed genes are in blue.

**Table 1.** Top and bottom five deferentially expressed genes each HC.

| Phenotype | Overexpressed genes |  |  | Underexpressed genes |  |  |
| --- | --- | --- | --- | --- | --- | --- |
|  | Genes | Log<sub>2</sub>FC | padj | Genes | Log<sub>2</sub>FC | padj |
| BALL | IRX2 | 12.72 | 2.19e<sup>-64</sup> | UNC5A | -6.49 | 7.19e<sup>-50</sup> |
|  | EPHA7 | 11.84 | 2.47e<sup>-113</sup> | CCR2 | -6.23 | 1.1e<sup>-73</sup> |
|  | KLKP1 | 11.34 | 4.2e<sup>-117</sup> | CHI3L1 | -6.02 | 1.78e<sup>-46</sup> |
|  | OR5P2 | 11.13 | 6.35e<sup>-41</sup> | HK3 | -6.01 | 1.54e<sup>-66</sup> |
|  | DSC3 | 11.03 | 1.12e<sup>-29</sup> | CEACAM3 | -5.98 | 1.13e<sup>-58</sup> |
| TALL | PEX5L-AS2 | 15.21 | 5.49e<sup>-119</sup> | IGHA1 | -6.92 | 1.02e<sup>-116</sup> |
|  | OR10T2 | 13.98 | 1.64e<sup>-69</sup> | IGHA2 | -6.75 | 1.46e<sup>-103</sup> |
|  | RNU7-7P | 13.09 | 3.39e<sup>-89</sup> | NRGN | -6.34 | 7.79e<sup>-98</sup> |
|  | OR10R3P | 12.51 | 1.19e<sup>-63</sup> | TARM1 | -6.29 | 1.85e<sup>-98</sup> |
|  | PCDH10 | 12.39 | 2.09e<sup>-198</sup> | RPH3A | -6.27 | 6.56e<sup>-153</sup> |
| MM | MAGEC2 | 16.65 | 2.77e<sup>-998</sup> | PZP | -8.76 | 8.63e<sup>-189</sup> |
|  | MAGEC1 | 15.09 | 2.54e<sup>-173</sup> | MIR3609 | -8.55 | 8.97e<sup>-242</sup> |
|  | LINC01518 | 14.32 | 2.54e<sup>-173</sup> | RN7SL5P | -8.52 | 5.19e<sup>-178</sup> |
|  | MAGEA3 | 12.87 | 1.95e<sup>-65</sup> | CD40LG | -8.47 | 4.31e<sup>-105</sup> |
|  | MAGEA6 | 12.72 | 1.1e<sup>-64</sup> | RMRP | -8.43 | 1.25e<sup>-148</sup> |
| AML | HMX3 | 9.74 | 1.45e<sup>-22</sup> | SCARNA10 | -9.56 | 3.66e<sup>-145</sup> |
|  | DSC3 | 9.72 | 4.73e<sup>-14</sup> | SCARNA5 | -7.47 | 5.48e<sup>-122</sup> |
|  | PRAME | 9.18 | 8.37e<sup>-78</sup> | SNORA74B | -7.06 | 1.26e<sup>-95</sup> |
|  | PCDH7 | 8.97 | 1.09e<sup>-06</sup> | KRT17P8 | -7.05 | 9.38e<sup>-87</sup> |
|  | IGF2BP1 | 8.9 | 8.66e<sup>-38</sup> | SNORA53 | -7.03 | 1.91e<sup>-111</sup> |

The top overexpressed and the top underexpressed genes are unique in each phenotype, except for DSC3, a member of the desmocollin subfamily that is overexpressed in both BALL and AML. BALL and TALL have members of the olfactory receptor family overexpressed, namely OR5P2 and OR10T2, respectively. OR10R3P, a pseudogene of the aforementioned family is also in the top overexpressed genes in TALL. The olfactory receptor family genes encode for G-protein coupled receptors related with the sense of smell, and have been reported to be overexpressed in several cancer cell lines (19), although their exact role in cancer is still unknown.

On the top underexpressed genes of BALL, TALL and MM are many genes related with the immune system, and in AML this top is conformed by four small nucleolar RNAs and a pseudogene of a keratin protein. Notwithstanding, the biological insights that we could obtain from these mere list of DEGs is limited; however, integrated with the topological analyses of GCNs and their functional enrichment analysis, we identified specific molecular processes that were exacerbated or diminished in HCs.

### GCNs of hematological malignancies share several interactions

In order to identify the main similarities in the architecture of the GCNs of the HCs, we inferred a network of the common interactions between the top-100,000 interactions of our four cancer networks. This network has 1,995 nodes and 4,140 edges.

We performed the community detection algorithm HiDeF (Hierarchical community Decoding Framework (20)) to identify robust structures across different scales, followed by a functional enrichment analysis using the clusterProfiler (21) R package (see Methods). This yielded a total of 189 hierarchical communities, from which 37 were enriched for at least one biological process (BP) from Gene Ontology (Supplementary Material 4).

The functional enrichment of these communities is represented in Fig. 4 in the form of a bipartite network of communities (circles) linked with their respective BPs (square diamonds). In that figure, communities are associated with those processes that resulted significantly enriched at the cut-off value (*p*_*val*_ *<* 1*e*^*−*10^).

**Fig. 4.**
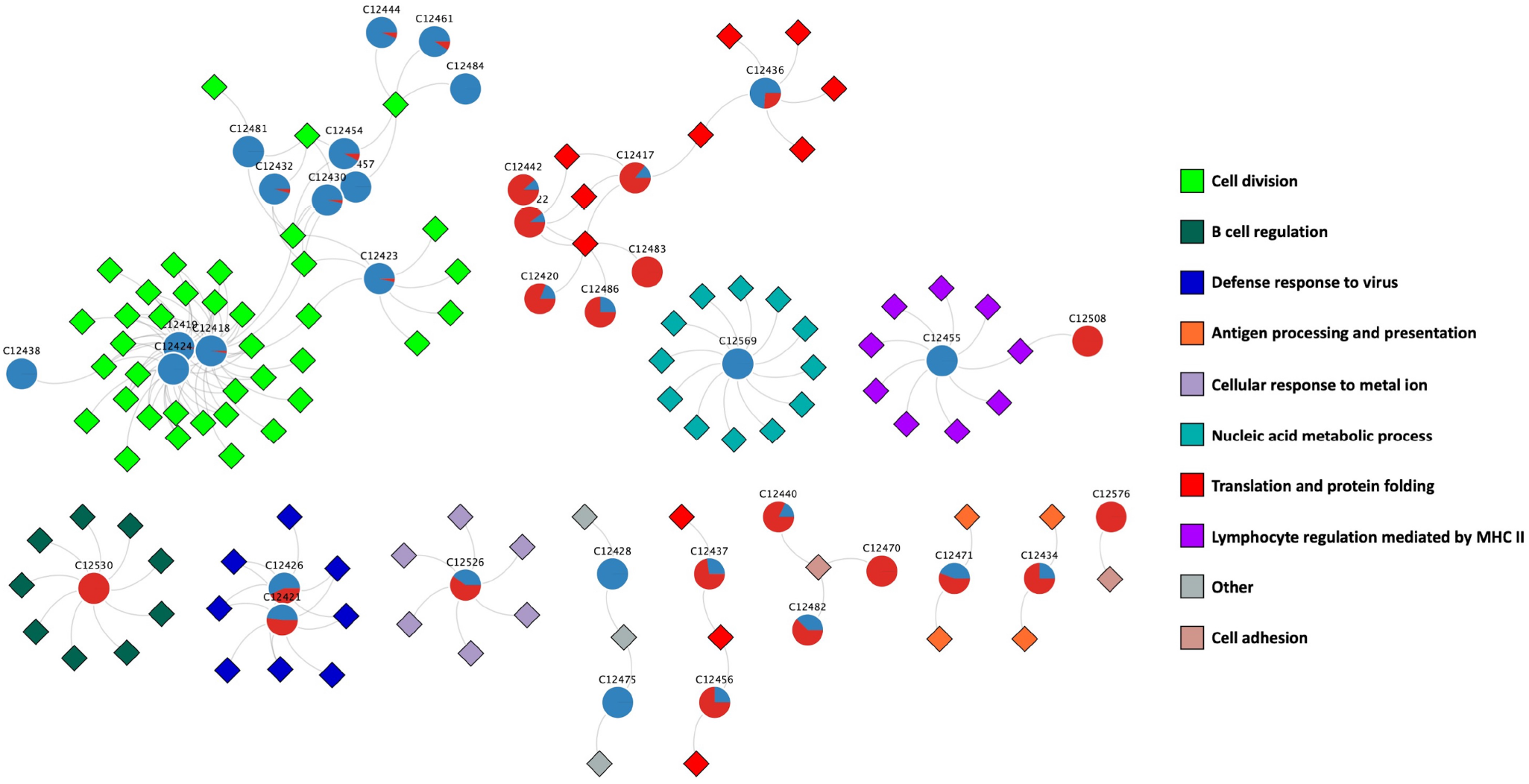
Bipartite network of communities (circles) detected in the intersection network and their associated biological processes (diamonds). The pie plots in the communities represent the proportions of the expression trends of the genes in those communities. A completely blue circle means that every gene in the community has an underexpression trend, and the opposite for a completely red circle. In this figure, we depict the differential expression trend of the analysis on multiple myeloma.

The latter results were combined with a differential expression analysis to determine the differential expression trend of those communities for each type of HC. This is shown in the pie plots on each community. Fig. 4 shows the differential expression trend for biological processes in MM. This allowed us to visualize the communities of the intersection network, the BPs that they are associated with, and to determine which BPs are enhanced or decreased in each HC in comparison to the normal bone marrow. The differential expression trend for the bipartite networks of the other HCs is shown in Supplementary Material 5.

As mentioned before, the largest component of the AML network is composed mostly of pseudogenes of riboproteins (RPs) and translation elongation factors. Several of these pseudogenes are also present in the intersection network, and are mainly overexpressed in all HCs. A large set of coexpressed riboprotein inter-chromosome genes has been also reported in networks from other 15 cancer tissues (16). We also found clusters of protein coding genes of ribonucleoproteins in our HCs. Interestingly, the interactions between this cluster of protein coding genes with the cluster of pseudogenes were scarce in the intersection network. The communities detected in this cluster of RPs were enriched for cytosplasmatic translation and other processes related with RNA processing.

Two communities in the intersection network (C12455 and C12508) are enriched for `immunoglobulin production`. C12455 is conformed by genes from the human leukocyte antigen (HLA) complex overexpressed in BALL and AML, and under-expressed in TALL and MM. C12508 is a community conformed by immunoglobulin light chain genes that are underexpressed on the three leukemias and overexpressed in the case of multiple myeloma, which could be associated with the elevated secretion of monoclonal immunoglobulin proteins by myeloma cells.

Another community in our intersection network (C12428) is enriched for `erythrocyte homeostasis` and `erythrocyte differentiation`, although with a little over *padj* = 1*e*^*−*10^, which is below our significance threshold. Notwithstanding, we consider this particular enrichment to be relevant, since anemia is one of the most common symptoms of hematological malignancies. Indeed, most genes in this community are underexpressed in every HC network, reaffirming that the production of erythrocytes is strongly impaired.

### The functional enrichment analysis on each HC network shows similar biological processes present in the cancer networks

The intersection network of the four HCs allowed us to identify which processes were represented by the same interactions in the four GCNs. Nevertheless, this analysis does not account for the shared BPs that are represented by different interactions in each phenotype, which could bias our comparison between the phenotypes. To have a better sense of the expression trends of the BPs in each phenotype, we performed the same community detection and functional enrichment analysis workflow on the individual cancer networks and identified the processes that were commonly present in the four phenotypes. We integrated this information in a heatmap (Fig. 5) that shows the expression trends of these common processes in each network.

**Fig. 5.**
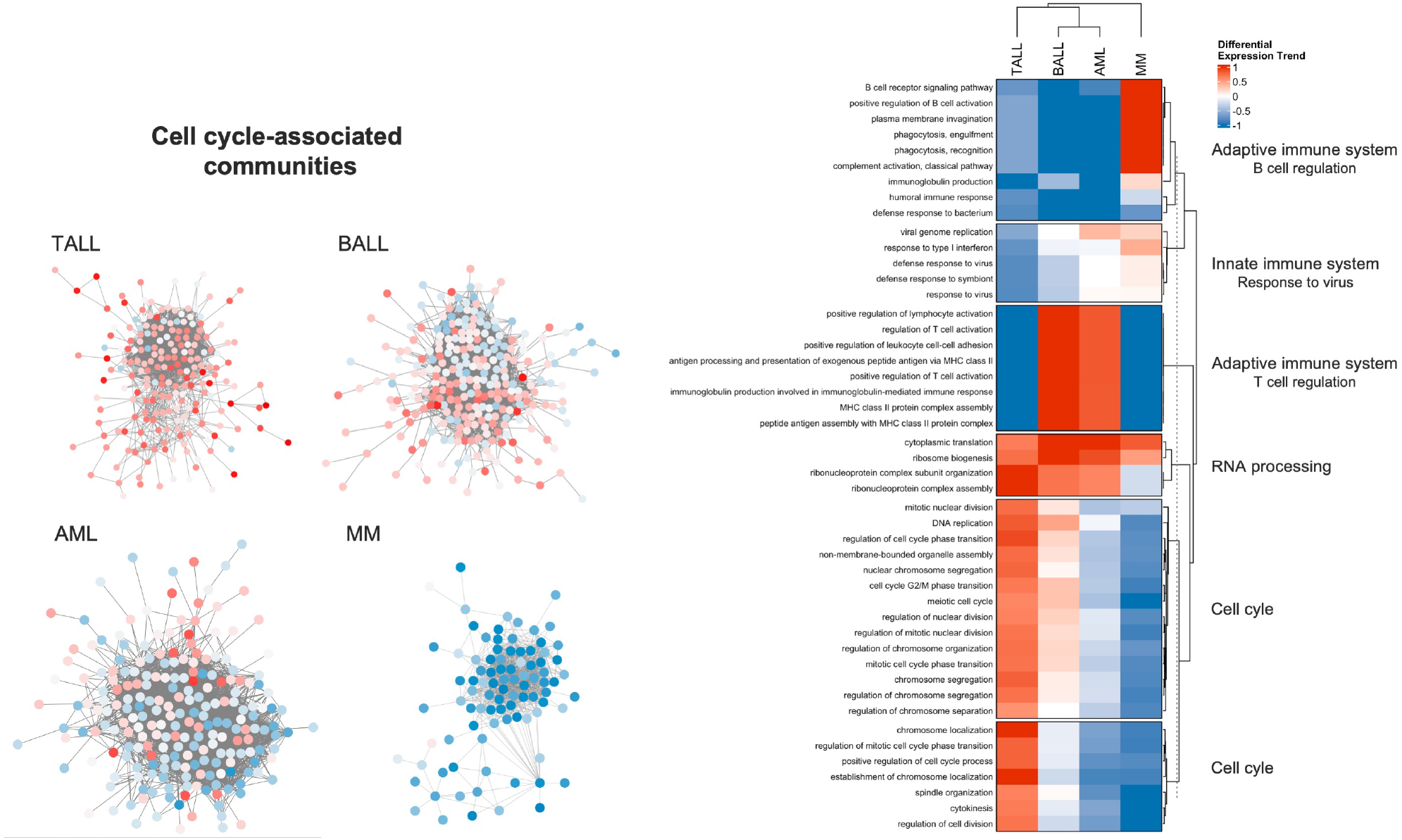
Differential expression trend of the common biological processes in the four HC networks. A value of 0 means that half of the genes enriched for a BP are overexpressed and the other half underexpressed.

This analysis shows a contrast in the differential expression trends of the BPs among HCs. For example, several communities are enriched for processes related to cell cycle and regulation of the immune system, but they have a clear different expression trend among the four phenotypes.

The main similarity in the figure are those BPs related with the assembly of ribonucleoprotein complexes, whose genes are mainly overexpressed. Meanwhile, the processes related with cell division present a mixture of differential expression trend, which is also depicted in the four networks at the left part of the figure.

In the case of multiple myeloma, all protein-coding genes except for two (ZWILCH and STAG1) are underexpressed. In AML we observed a similar underexpression trend in cell cycle processes. However, in this case there are some overexpressed genes that have an active role in the regulation of cell cycle, like CKS1B, that is crucial for the function of cyclin dependent kinases and for progression through cell cycle in cancer cells (22, 23). For BALL, the majority of the most significant differentially expressed genes are overexpressed and also related with cell proliferation. For example, MYBL2 is known to be ubiquitously expressed on proliferating cells (24). In TALL, almost every gene related with cell cycle is overexpressed. The most relevant underexpressed genes are E2F8 (log2FoldChange = -0.9143, padj = 1.24*e*^*−*06^) and PIF1 (log2FoldChange = -0.6803, padj = 1.5*e*^*−*04^).

An important amount of common biological processes of the four cancer GCNs are related to the adaptive immune system. This was expected given the origin of these malignancies. The processes of B cell activation are under-regulated in every phenotype except for MM. On the other hand, the processes related with T cell regulation are exacerbated in TALL and MM, but diminished in BALL and AML.

These shared BPs are very similar to those enriched on the intersection network. This indicates that hematological malignancies have a shared set of affected BPs, and that these processes are probably deregulated through different mechanisms in each HCs, given their unique expression pattern in each phenotype.

### Validation with other datasets

It is worth noticing that the results presented here have been obtained from calculations using the TCGA hematological cancer dataset. To validate these results we decided to perform all the calculations using two different datasets: one from the Royal Children Hospital (n = 127 (17)) containing only samples from BALL patients, and another from Saint Jude’s Hospital (n = 465 in total, 174 for AML, 145 for TALL, 146 for BALL (18)).

We pre-processed, normalized, and constructed networks for each of these datasets. Given that no normal bone marrow samples were identified in them, we normalized the data with respect to the TCGA normal bone marrow samples. Multiple myeloma RNASeq dataset was not found elsewhere. We calculated the fraction of intra-chromosome interactions, the interactions between different RNA biotypes in each networks, and the community structure of the networks.

As expected, the RNA molecules are grouped by biotype and a cluster of pseudogenes and lncRNAs takes relevance in the top 10,000 interactions in these networks, such as in the TCGA dataset (Fig. 6A). On the other hand, the fraction of intra-chromosomal interactions resulted higher in HCs than the control networks in all cases (Fig. 6B). In this respect, the BALL and TALL networks of SJ datasets behave very similarly between them, and the AML network remains as the networks with the lesser amount of cis-edges (in agreement with the case of TCGA AML network). Lastly, several of the BPs enriched in the validation GCNs are also found in the TCGA networks. This can be appreciated by the number of shared processes between TCGA and the validation datasets (Fig. 6C). In all cases, the shared processes between TCGA and validation GCNs are more than those exclusive for any phenotype, except for TALL in the TCGA dataset.

**Fig. 6.**
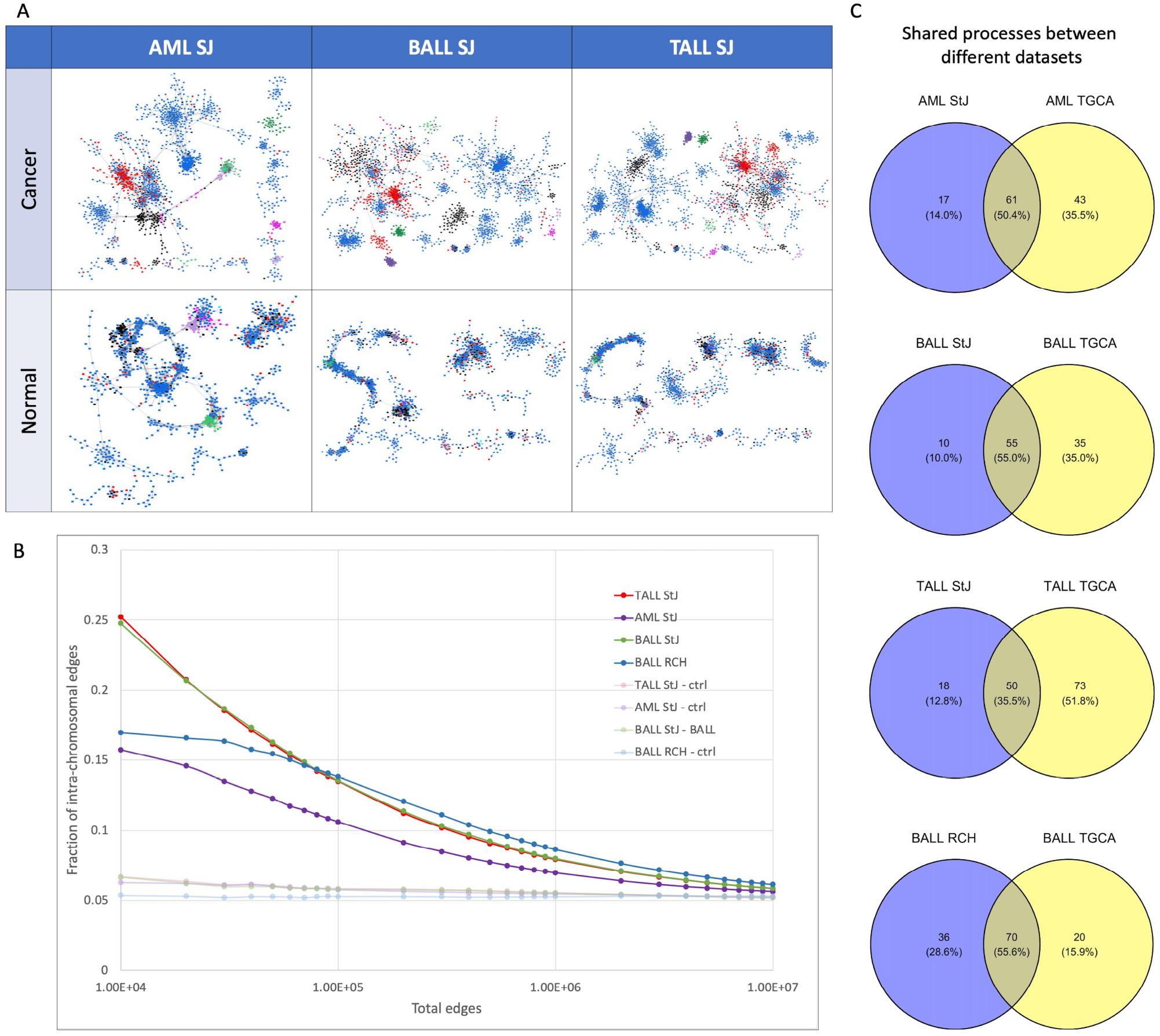
Validation of findings with other datasets. A) GCNs from AML, BALL and TALL from St. Jude dataset with their respective control networks. B) Intra-chromosomal proportion of HCs from RCH and SJ datasets. C) Shared biological processes between communities from networks from TCGA and the validation datasets.

## Discussion

This study sought to characterize the gene co-expression landscape of hematopoietic cancers. Here, using a total of 1,495 samples (AML = 159, TALL = 245, BALL = 210, MM = 844, normal bone marrow = 37), we constructed GCNs for each cancer phenotype and analyzed their architecture and functional enrichments, and compared them against the normal bone marrow.

As expected, the architecture of the cancer networks are different from the normal bone marrow in two main aspects: the number and size of network components, and 2) the chromosomal location of genes that composed them. As mentioned before, the cancerous networks have several small components, many of them made up of genes from the same chromosome. This result coincide with the previous GCNs analyzed from breast cancer (13, 25), lung (26), and clear cell renal carcinoma (27, 28). In these studies, the cancer networks have a diminished amount of inter-chromosomal interactions. With these results, we show that this phenomenon is not restricted to carcinomas. Hence, we could argue that the loss of inter-chromosomal interactions is a common deregulated feature of the co-expression landscape in cancer.

The topological differences between the four HC networks are noteworthy, especially regarding the augmented intrachromosomal interactions in comparison with the normal bone marrow. In Fig. 1 we can observe that the four HCs networks have different proportions of cis-interactions. This suggests that, although the loss of inter-chromosomal regulation is present in the networks, there must be a difference in the deregulation of co-expression between the four HCs.

The case that most resembles to the phenomenon observed in carcinomas is MM, where the loss of long-range interactions is much more evident. One difference between the MM samples and the samples from leukemias is the ages of the patients in each group. The MM group is conformed by samples from adults, while our leukemias samples correspond mainly to pediatric patients (Fig. 2B). This could suggest that the loss of inter-chromosomal interactions could be associated with the age of the patients. Further studies on samples from pediatric cancer patients are needed to further explore this possibility.

The exacerbated co-expression between pseudogenes in HCs is a feature that, as far as the authors are aware of, has not been reported elsewhere in the current literature. Previous reports regarding the expression of pseudogenes have revealed that their transcription is tissue specific, suggesting an active role in the tissues (29, 30).

A compelling explanation regarding the pseudogenes function is centered on their role as competitive endogenous RNA (ceRNA). According to this, the transcripts of pseudo-genes regulate their parental gene expression through competition for its regulatory elements, such as miRNAs (31, 32). However, our results show that most pseudogenes have very strong co-expression relationships between them, which are even stronger than the interactions with their respective parental genes, and that occur between pseudogenes with different parental genes. This suggests that these co-expression interactions are not a manifestation of the ceRNA capability of pseudogenes.

It could be possible that a deregulation on the transcription of RPs is allowing their co-expression, and that this serves as a mechanism used by cancer cells to sustain high rates of protein translation that contribute to their proliferation. A similar mechanism could be affecting the expression pattern of their pseudogenes, since they retain some characteristics of their parental genes, which could prompted their co-expression. Importantly, pseudogene complexes observed in HC networks are mostly overexpressed (Figs. 2C and 3). However, the question would still remain as of what type of signals or mechanisms are indicating the transcriptional machinery to transcribe these genes at a similar rate.

Interestingly, this pattern in the pseudogenes co-expression could be associated with the loss of inter-chromosomal regulation, since the network with the most amount of relationships between pseudogenes (AML) has the least amount of cis-chromosomal interactions. Furthermore, the opposite occurs in MM, which has the least amount of relationships between pseudogenes and the highest proportion of cis-chromosomal edges. Further studies on the co-expression landscape of non-protein coding genes should be carried out to address this possibility.

The mechanism regulating these protein coding genes and their pseudogenes should be similar, but not exactly the same, since otherwise one would detect interactions between pseudogenes and protein coding genes in the top interactions. Further research on the subject of the transcription regulation of RPs and their pseudogenes in cancer is necessary to propose a more robust hypothesis regarding their co-expression patterns.

Given the common site of origin of the samples (the bone marrow) we investigated the interactions shared between the four HCs networks. Most of these shared interactions are inter-chromosomal, and are enriched for specific biological processes. Nevertheless, the genes in this intersection network have a phenotype-specific differential expression trend. This suggests differences on their regulation in each disease.

To better assess these differences, we analyzed the functional enrichment on the individual networks and identified the common processes between them. These shared BPs are similar to those enriched on the intersection network. In this regard, the finding that every gene in the cell division-associated communities for MM (with the exception of ZWILCH and STAG1) is underexpressed, could mislead to the conclusion that the myeloma cells are not proliferating. We believe that this underexpression pattern must be retained on myeloma cells from their normal cell of origin, the long-lived plasma cell, which no longer divides after its differentiation from naive B cells (33–35).

It is worth noting that MM is a progressive disease, which typically arises after monoclonal gammopathy of undetermined significance (MGUS) and smoldering multiple myeloma (SMM). The progression from these stages to MM is approximately 1% per year for MGUS and 10% per year for SMM (36). Given these relatively slow rates of progression, we could assume that the proliferation of malignant plasma cells occurs slowly and in a lower rate than the proliferation of normal cells inside the normal bone marrow. The low proliferation rates would also explain the underexpression of genes related with cell cycle, and the differences of this in comparison with the acute leukemias, which are malignancies of rapid evolution. Since the RNA-Seq samples for these analyses were performed in bulk, we are not able to dissect the specific expression profile of the cells into the sample. Given that the BM has a relatively small amount of plasma cells (around 1% of total cells (37)), we believe that further studies involving myeloma and plasma cells could better asses the cell cycle regulation in the disease.

The analysis of cell cycle related communities in AML and BALL clearly reflects an ongoing proliferation, with the overexpression of genes like CKS1B, PCLAF or PCNA (Fig. 5). CKS1B is crucial for the function of cyclin dependent kinases and for progression through cell cycle in cancer cells (22, 23). PCLAF and PCNA genes are involved in DNA synthesis and the G1/S transition (38). However, we also detected several underexpressed genes that regulate the mitotic spindle and chromosome segregation (like KIFC1, ANLN, etc.). This suggests that, although there is an active proliferation in these phenotypes, the processes responsible for an adequate division are downregulated, which could contribute to (or generate) the genomic instability in theses diseases.

An important amount of common biological processes of the four cancer GCNs are related to the adaptive immune system. This was expected given the origin of these malignancies. The processes of B cell activation are downregulated in every phenotype except for MM. On the other hand, the processes related with T cell regulation are diminished in TALL and MM, but exacerbated in BALL and AML. We believe that this expression trend is linked to the cells of origin of the malignancies. B cells and members of the myeloid lineage are professional antigen presenting cells (APCs) which activate T cells through MHC molecules. Hence, the overexpression of genes related with this processes (mainly HLA genes) could be a legacy from the normal cell of origin that are specialized in the presentation of antigens to T cells.

The genes enriched for B cell regulation are mainly immunoglobulins, and their overexpression in MM is expected as its normal cell of origin, the plasma cell, produces antibodies and do not have the ability to present antigens to T cells (which explains the underexpression of HLA genes in this phenotype).

Together, our results on the validation datasets show that different datasets of the same HCs reflects the underlying deregulation of the co-expression landscape manifested through the loss of inter-chromosomal interactions in these phenotypes. Furthermore, and despite the validation networks have not the exact same architecture of the TCGA networks, most of the biological processes represented by the interactions in the validation networks are also found in the enrichment of the TCGA GCNs.

In order to unveil the mechanisms behind the loss of interchromosome gene co-expression in cancer, other omic technologies have been evaluated. For instance, copy number variation data, transcription factor binding sites, or micro-RNA co-expression (28, 39–41). However, none of those omics have been determinant to explain the significant decrease of long-range gene co-expression. Further studies complementary to the gene co-expression analysis, such as the methylome profile, or Hi-C data in different cancer tissues are also ongoing.

Complementary to the last, a recent study on the protein co-expression landscape of breast cancer, showed that the loss of long distance co-expression is not observed at the proteome level. The most relevant protein co-expression interactions occur between members of the same protein family or between proteins with similar functions (42). So far, the loss of long-range co-expression has only been seen at the transcriptomic level of regulation.

Despite the lack of a mechanistic explanation for the bias of cancer GCNs to aggregate gene clusters from the same chromosome, we argue that the different phenomena described here must be part of the broad deregulation that occurs in HCs at the transcriptional level. The significant coincidence between the TCGA-derived GCNs with the Saint Jude and Royal Children Hospital datasets, together with the statistical assessment of network topologies and enrichment analyses, allows to suggest that this phenomenon is not an artefact of the datasets nor of the methods that determine the gene correlation. We observed a high similarity between networks constructed with MI and Spearman correlation (See Methods and Supplementary Material 5). Furthermore, in breast cancer molecular subtypes, the loss of long-distance co-expression have been also reported in GCNs inferred with Pearson correlation (43). Therefore, the co-expression method does not influence the phenomenon.

### Final considerations and future perspectives

In this work, we performed a global analysis of the transcriptional landscape from four hematopoietic cancers. Through the use of GCNs we were able to identify several gene interactions that are shared by the four HC networks, which translates in biological processes common to these malignancies. Nevertheless, the genes in these common interactions have a unique differential expression profile that reflects differences in their regulation in each phenotype. It is necessary to further analyze the topology of the individual GCNs of HCs to understand how the circuits of normal hematopoietic cells are deregulated to promote these diseases.

An aspect that also needs to be further explored is the topology of normal bone marrow network. Here, the data set used was made up of 37 RNA-Seq samples, which was the total number of normal bone marrow samples present in the TCGA dataset at the moment of the analysis. Given that we performed a normalization workflow for each HC data set with the same normal group, we obtained a total of four networks for the normal bone marrow. However, our approach can be highly sensitive to the number of samples analyzed, which could represent a limitation to detect co-expression interactions with a high confidence. Further studies using a larger group size could provide more robust networks to further compare the topology of the normal and cancerous phenotypes and identify interactions deregulated in the disease. Nevertheless, our results show that our network theory approach to investigate cancer phenotypes is able to elucidate relevant clinical features. For example, the communities associated with erythrocyte development that are downregulated, evidence the anemia suffered by patients of these diseases. To clarify whether these biological features of HCs are a consequence of the deregulation of the normal co-expression landscape, additional research is necessary.

An important aspect of the GCNs of hematologic cancers should be taken into account: the effect of the ages of the patients whose samples are included in these type of studies. Multiple myeloma, the network with the more pronounced loss of inter-chromosomal co-expression, was inferred with all patients in advanced edges, meanwhile the rest of HCs were obtained from pediatric patients.

Future investigation should be done involving lymphomas samples, which were neglected here given the lack of an appropriate control group to compare these samples. Single-cell RNA-Seq data will certainly improve the accuracy of the gene co-expression landscape. To analyze other -omics data could add molecular interactions that may help to disentangle the molecular mechanisms that make different these hematopoietic malignancies. However, with this approach we emphasize the importance of this kind of analyses to better understand the co-expression landscape in cancer and its associated functional features.

## Methods and Materials

The computational code to develop the present pipeline is available on GitHub. It can be found at https://github.com/AKNG97/Network_structure_of_hematopoietic_cancers. All data shown here can be reproduced using this code.

### Data acquisition

The samples used in this worked were retrieved from https://portal.gdc.cancer.gov, The Cancer Genome Atlas consortium. The cancer samples included those categorized as “Recurrent Blood Derived Cancer - Bone Marrow” and “Primary Blood Derived Cancer - Bone Marrow” of patients with a primary diagnosis of “acute myeloid leukemia, nos”, “precursor b-cell lymphoblastic leukemia”, “t lymphoblastic leukemia/lymphoma” and “multiple myeloma”. Also, normal bone marrow samples used were those categorized as “bone marrow normal”.

The validation datasets were retrieved from (17) (RCH dataset) and from https://www.stjude.cloud/research-domains/pediatric-cancer (St. Jude dataset). The St. Jude data downloaded corresponded to the samples of diagnoses of “B cell Acute Lymphoblastic Leukemis, NOS”, “Acute Myeloid Leukemia” and “T cell Acute Lymphoblastic Leukemis, NOS”.

A complementary annotation file was obtained from BioMart in order to obtain the chromosome name, gene type, GC content, gene start and gene end. The gene length was inferred by the rest of the gene end minus the gene start.

### Data pre-processing

Each cancer dataset was preprocessed along with the normal bone marrow dataset. Hence, we obtained four cancer expression matrices and four normal bone marrow matrices, one for each cancer phenotype. The first step in our workflow was to apply according filters to remove low count genes by means of the TCGAbiolinks package in R (44), as follows:

1. Genes on the 0.25 quantile
2. Genes with an expression value of 0 across the majority of the samples in the datasets
3. Genes with a mean expression less than 10 were removed.

After filtering the genes, we performed normalization procedures for several biases of the sequencing technology using EDASeq (45) and NOISeq (46) in R:

1. Normalization for the length transcript and the GC content of the genes.
2. Between-lane normalization followed by Trimmed Mean of M values (TMM).
3. Lastly, we apply the multidimensional noise reduction algorithm ARSyN to assure that the cancer and normal samples were separate and clustered correctly, PCA visual exploration was used to confirm this (Supplementary Material 6).

### Data processing

#### Differential expression analysis

We tested for differentially expressed genes in each cancer data set in comparison to the normal bone marrow data set using the DESeq2 (47) package in R, which is based on a model using the negative binomial distribution.

#### Networks construction

Despite mutual information (MI) is a good estimator of dependency between variables, it can be strongly affected by the number of samples in the analysis. Considering the small amount of samples in our control group, we constructed networks with both MI and Spearman correlation, which is less dependent on the number of samples, although it specifically detects monotonic relationships. To decide with which type of network (MI or Spearman correlation) we should continue our analysis, we identified the gene interactions that are shared between each network in different cut-off points (Supplementary Material 5).

Mutual information values were calculated by implementing the ARACNe algorithm (48). Spearman correlation values were calculated using the function rcorr from the package Hmisc in R. These co-expression values were sorted using their absolute values for further analyses.

Using the function ks.test of the package dgof in R, we performed the Kolmogorov-Smirnov test on the fraction of interchromosome interactions at different cut-off points on the networks (Supplementary material 2).

#### Intersection network construction

The intersection network was constructed from the common interactions between each of the four cancer GCNs using their respective 100,000 strongest interactions.

#### Community detection and functional enrichment analysis

In order to correlate topological structures inside our networks with biological processes, we first performed the community detection algorithm HiDeF through the CyCommunityDetection app in Cytoscape (49). This integrates the concept of persistent homology with existing algorithms for community detection in order to identify robust structures intrinsic to a dataset. We applied HiDeF with its default setting in Cytoscape using the Louvain algorithm. The communities detected were then enriched using the R package clusterProfiler to correlate the genes inside the communities to biological processes of Gene Ontology. We used the EnsemblID of each gene in this analysis and considered that a community was enriched if the correlation to a biological process had an adjusted p-value less than 1*e*^*−*10^).

#### Analysis of the common biological processes on the four HCs

We calculated the percentage of up and down expressed genes on the communities enriched for each BP on the individual cancer networks. We rested the proportions and called the result `differential expression trend` for a BP on a specific phenotype. A value of 1 depicts that every gene enriched for a BP has an overexpression tendency, a value of -1 depicts that every gene has an underexpression tendency, and a value of 0 depicts that there are the same proportion of up and down expressed genes enriched for the process.

## Supporting information

Kolmogorov-Smirnov p-values for the comparison between fraction of intra-chromosomal interactions in normal and cancer tissues at different thresholds

Network files for the top-100,000 interactions in the five phenotypes

Differential gene expression analysis for all hematopoietic cancers.

Community structure of the four HC networks. These files contain the name of all communities and the corresponding elements for each community.

Top: Bipartite networks of communities and enriched biological processes. Bottom: Shared interactions between MI and Spearman networks.

Quality control for data pre-processing.

## Acknowledgments

Arturo Kenzuke Nakamura-Garcia is a doctoral student from Programa de Doctorado en Ciencias Biomédicas, Universidad Nacional Autónoma de México (UNAM) and received fellowship 806341 from CONACYT. Both authors thank to National Institute of Genomic Medicine (INMEGEN) where this work took place.

## Supplementary material

1. Network files for the top-100,000 interactions in the five phenotypes.
2. Kolmogorov-Smirnov p-values for the comparison between fraction of intra-chromosomal interactions in normal and cancer tissues at different MI thresholds.
3. Differential gene expression analysis for all hematopoietic cancers.
4. Community structure of the four HC networks. These files contain the name of all communities and the corresponding elements for each community.
5. Top: Bipartite networks of communities and enriched biological processes. Bottom: Shared interactions between MI and Spearman networks.
6. Quality control for data pre-processing.

