## Supplementary figures and images for "The network structure of hematopoietic cancers"

### supplementary_figure.jpg

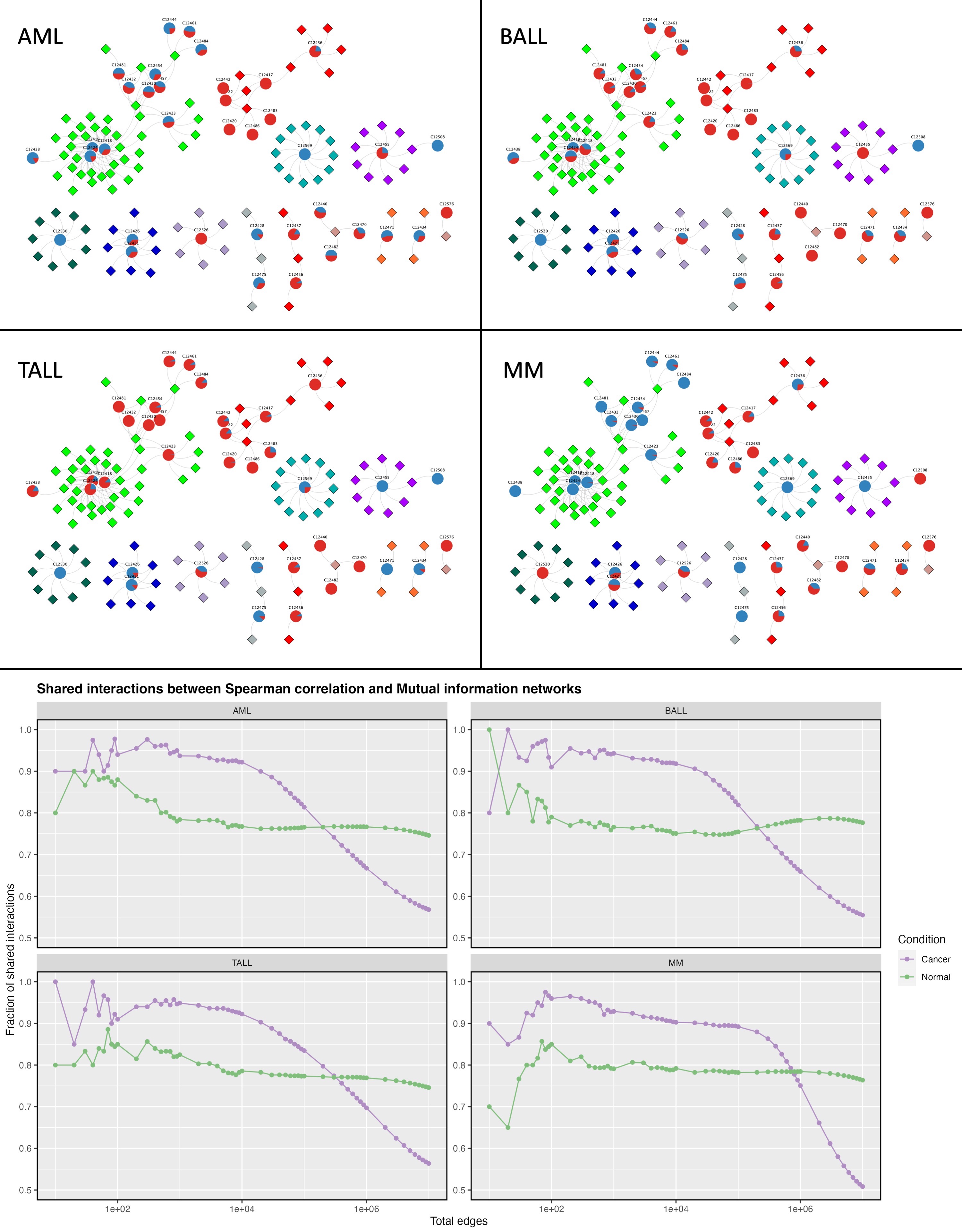
