## Supplementary material for "The network structure of hematopoietic cancers": Quality control for data pre-processing.: QCreport_AML_After.pdf

### Quality Control of Expression Data

*Generated by NOISeq on 24 Feb 2023, 13:38:23*

#### Content

| *Plot* | *Description* |
|---|---|
| **Biotype detection** | Biotype abundance in the genome with %genes detected (counts > 0) in the sample/condition.<br>Biotype abundance within the sample/condition. |
| **Biotype expression** | Distribution of gene counts per million per biotype in sample/condition (only genes with counts > 0). |
| **Saturation** | Number of detected genes (counts > 0) per sample across different sequencing depths |
| **Expression boxplot** | Distribution of gene counts per million (all biotypes) in each sample/condition |
| **Expression barplot** | Percentage of genes with >0, >1, >2, >5 or >10 counts per million in each sample/condition. |
| **Length bias** | Mean gene expression per each length bin. Fitted curve and diagnostic test. |
| **GC content bias** | Mean gene expression per each GC content bin. Fitted curve and diagnostic test. |
| **RNA composition bias** | Density plots of log fold changes (M) between pairs of samples.<br>Confidence intervals for the median of M values. |
| **Exploratory PCA** | Principal Component Analysis score plots for PC1 vs PC2, and PC1 vs PC3. |

### Biotype detection

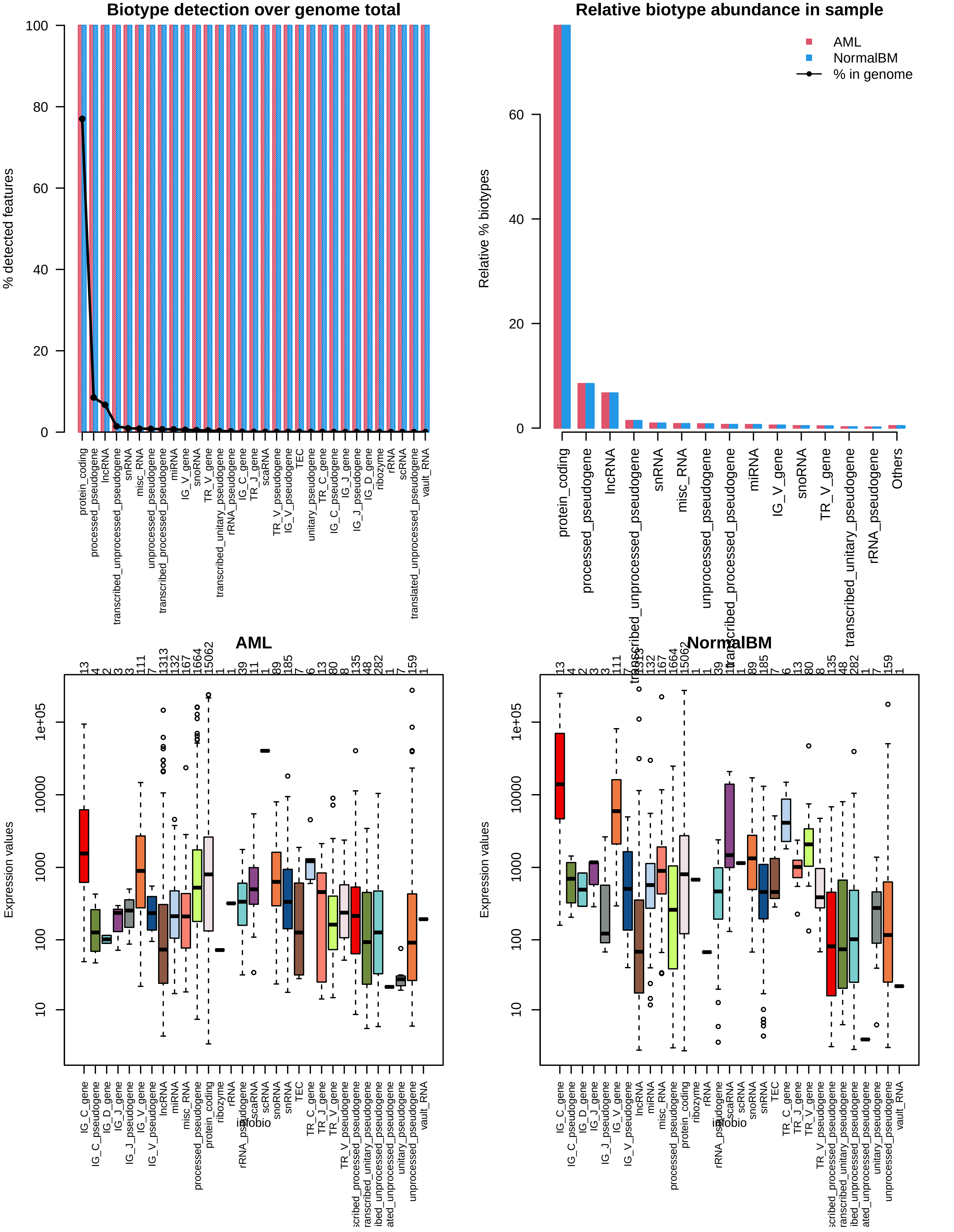

### Sequencing depth & Expression quantification

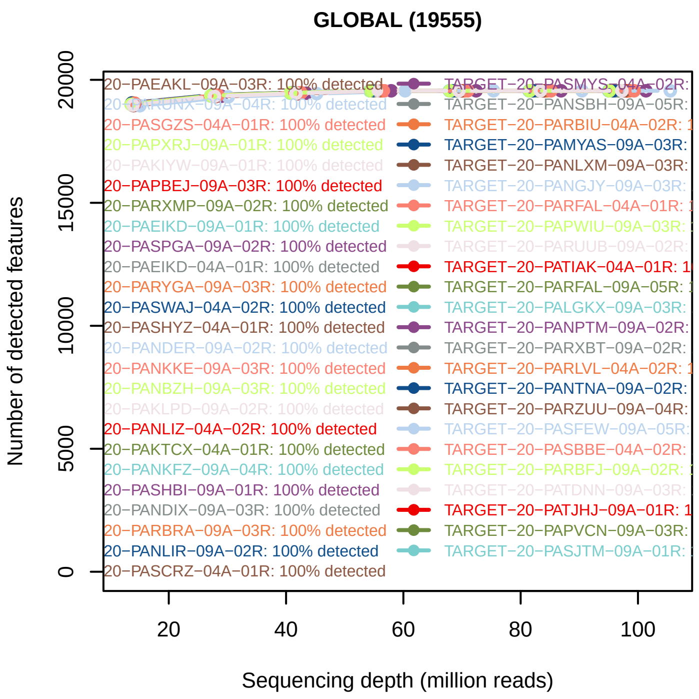


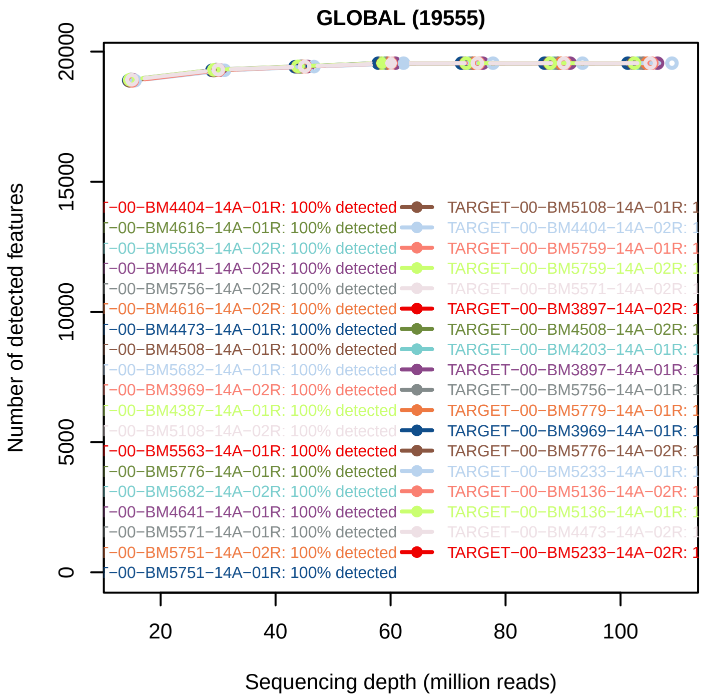


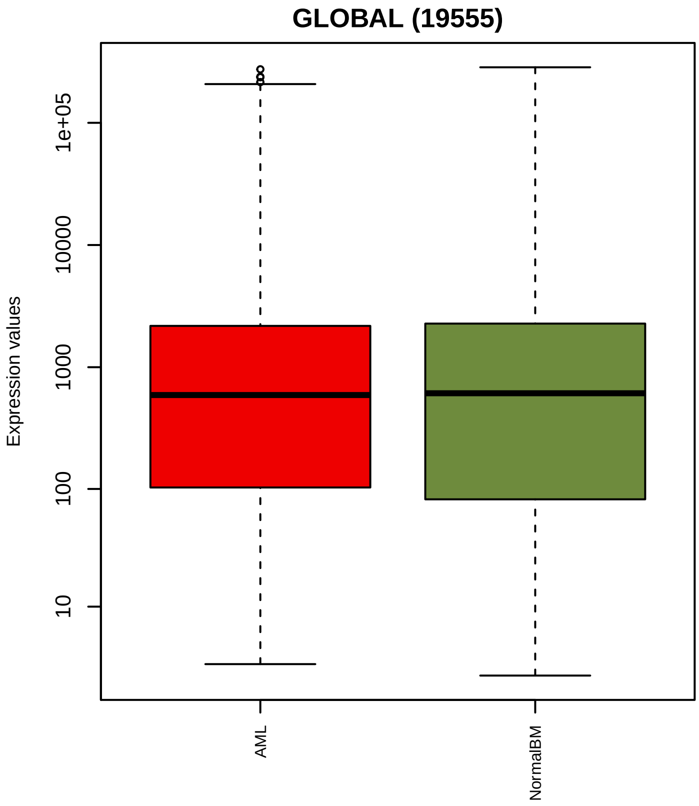


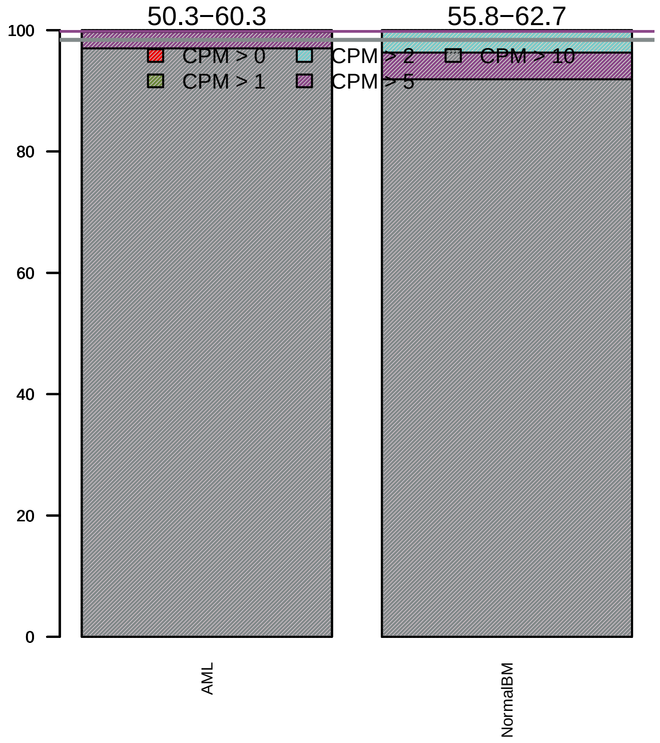

### Sequencing bias detection

#### *Diagnostic plot for feature length bias*

WARNING. At least one of the model p−values was lower than 0.05, but R2 < 70% for at least one condition.

Normalization for correcting length bias could be advisable.
Plese check in the plots below the strength of the relationship between length and expression.

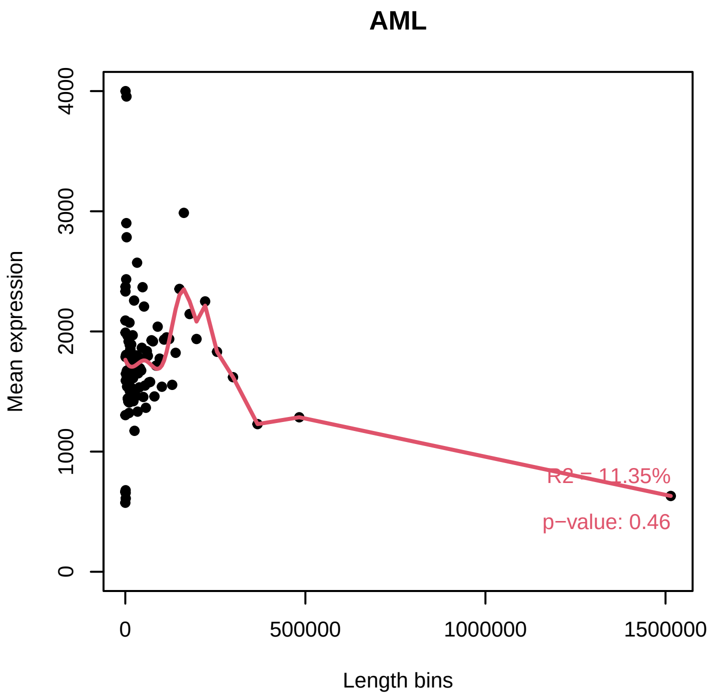


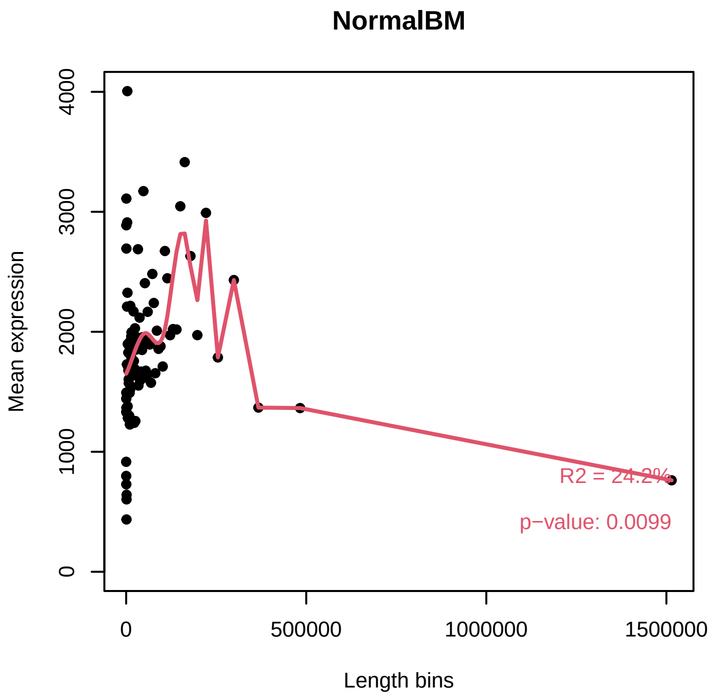


#### *Diagnostic plot for GC content bias*

PASSED. No normalization for correcting GC content bias is required.

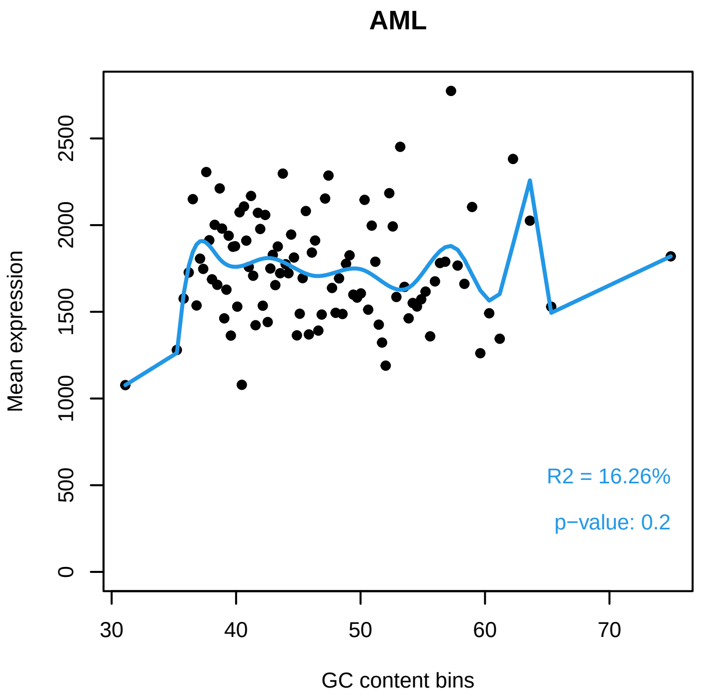


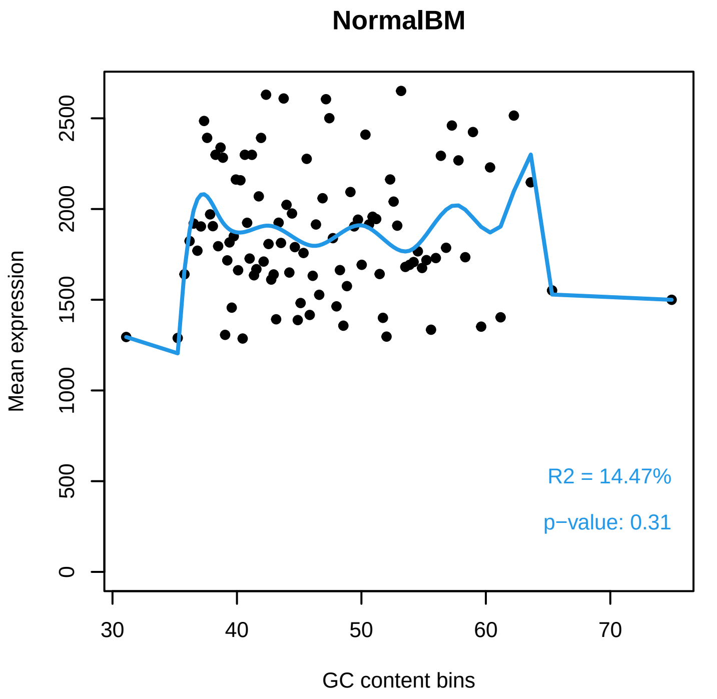

#### *Diagnostic plot for differences in RNA composition*

FAILED. There is a pair of samples with significantly different RNA composition

Normalization for correcting this bias is required.

##### Reference sample: TARGET−20−PASTUH−04A−01R

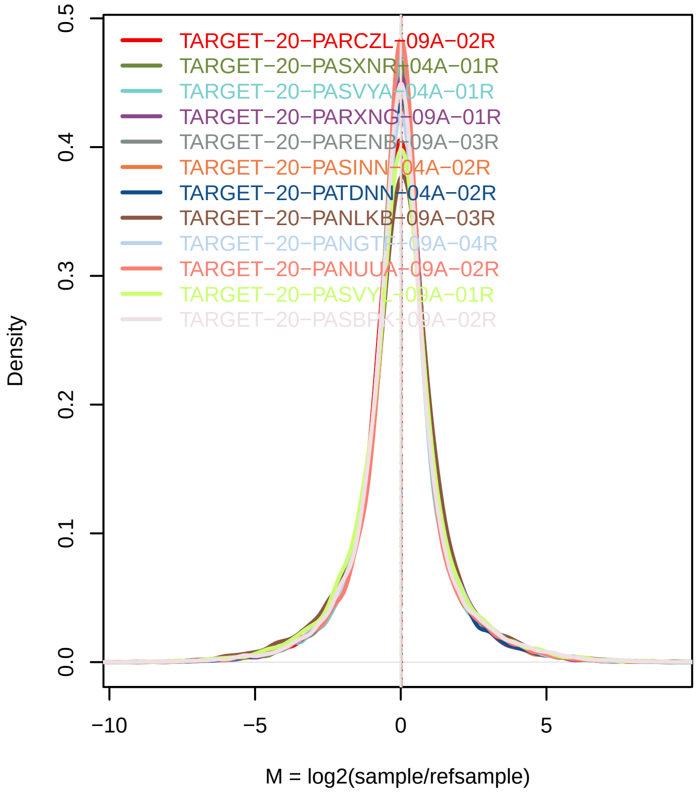


##### Confidence intervals for median of M values

| Sample | 0.01% | 99.99% | Diagnostic Test |
|---|---|---|---|
| TARGET−20−PARCZL−09A−02R | −0.0197 | 0.0398 | PASSED |
| TARGET−20−PASXNR−04A−01R | −0.0344 | 0.01 | PASSED |
| TARGET−20−PASVYA−04A−01R | −0.0304 | 0.0182 | PASSED |
| TARGET−20−PARXNG−09A−01R | −0.0113 | 0.0378 | PASSED |
| TARGET−20−PARENB−09A−03R | −0.0047 | 0.0462 | PASSED |
| TARGET−20−PASINN−04A−02R | −0.0184 | 0.0384 | PASSED |
| TARGET−20−PATDNN−04A−02R | −0.0047 | 0.0478 | PASSED |
| TARGET−20−PANLKB−09A−03R | 0.0126 | 0.0662 | FAILED |
| TARGET−20−PANGTT−09A−04R | −0.0093 | 0.0369 | PASSED |
| TARGET−20−PANUUA−09A−02R | −0.0157 | 0.0359 | PASSED |
| TARGET−20−PASVYL−09A−01R | −0.0259 | 0.0267 | PASSED |
| TARGET−20−PASBPK−09A−02R | −0.0304 | 0.0227 | PASSED |
| TARGET−20−PARAJX−[illegible]A−02R | 0.0312 | 0.0871 | FAILED |
| TARGET−20−PANVGE−[illegible]A−02R | −0.0261 | 0.0291 | PASSED |
| TARGET−20−PASTTW−[illegible]A−01R | −0.0218 | 0.0255 | PASSED |
| TARGET−20−PARCUK−[illegible]A−03R | −0.0182 | 0.0449 | PASSED |
| TARGET−20−PASIBG−[illegible]A−01R | −0.0266 | 0.0224 | PASSED |
| TARGET−20−PARUNX−[illegible]A−01R | −0.0321 | 0.0242 | PASSED |
| TARGET−20−PASVVS−[illegible]A−03R | 0.0059 | 0.0595 | FAILED |
| TARGET−20−PASXYG−[illegible]A−05R | −0.0118 | 0.0428 | PASSED |
| TARGET−20−PARBIU−[illegible]A−03R | −0.0113 | 0.0352 | PASSED |
| TARGET−20−PAPXWI−[illegible]A−03R | −0.0012 | 0.0478 | PASSED |
| TARGET−20−PARTAL−[illegible]A−01R | −0.0059 | 0.0345 | PASSED |
| TARGET−20−PARHVK−[illegible]A−04R | 0.0071 | 0.0534 | FAILED |
| TARGET−20−PASCFW−[illegible]A−01R | 0.0196 | 0.0647 | FAILED |
| TARGET−20−PAPWYK−[illegible]A−03R | −0.0012 | 0.0454 | PASSED |
| TARGET−20−PARCZL−[illegible]A−02R | −0.0143 | 0.0352 | PASSED |
| TARGET−20−PARUTH−[illegible]A−04R | −9e−04 | 0.0482 | PASSED |

### Exploratory PCA

Use this plot to see if samples are clustered according to the experimental design.

Use ARSyNseq function to correct potential batch effects.

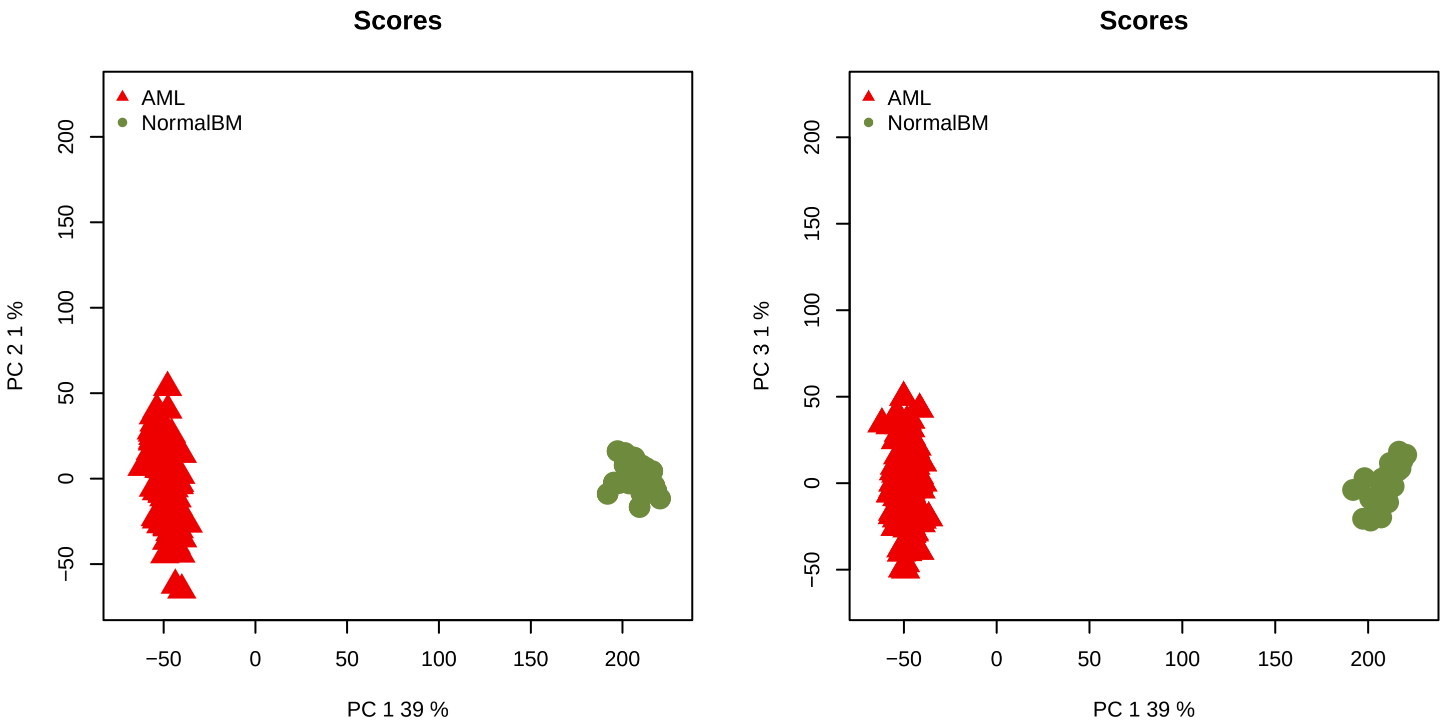
