## Supplementary material for "The network structure of hematopoietic cancers": Quality control for data pre-processing.: QCreport_AML_Before.pdf

### Quality Control of Expression Data

*Generated by NOISeq on 24 Feb 2023, 13:08:47*

#### Content

| *Plot* | *Description* |
|---|---|
| **Biotype detection** | Biotype abundance in the genome with %genes detected (counts > 0) in the sample/condition.<br>Biotype abundance within the sample/condition. |
| **Biotype expression** | Distribution of gene counts per million per biotype in sample/condition (only genes with counts > 0). |
| **Saturation** | Number of detected genes (counts > 0) per sample across different sequencing depths |
| **Expression boxplot** | Distribution of gene counts per million (all biotypes) in each sample/condition |
| **Expression barplot** | Percentage of genes with >0, >1, >2, >5 or >10 counts per million in each sample/condition. |
| **Length bias** | Mean gene expression per each length bin. Fitted curve and diagnostic test. |
| **GC content bias** | Mean gene expression per each GC content bin. Fitted curve and diagnostic test. |
| **RNA composition bias** | Density plots of log fold changes (M) between pairs of samples.<br>Confidence intervals for the median of M values. |
| **Exploratory PCA** | Principal Component Analysis score plots for PC1 vs PC2, and PC1 vs PC3. |

### Biotype detection

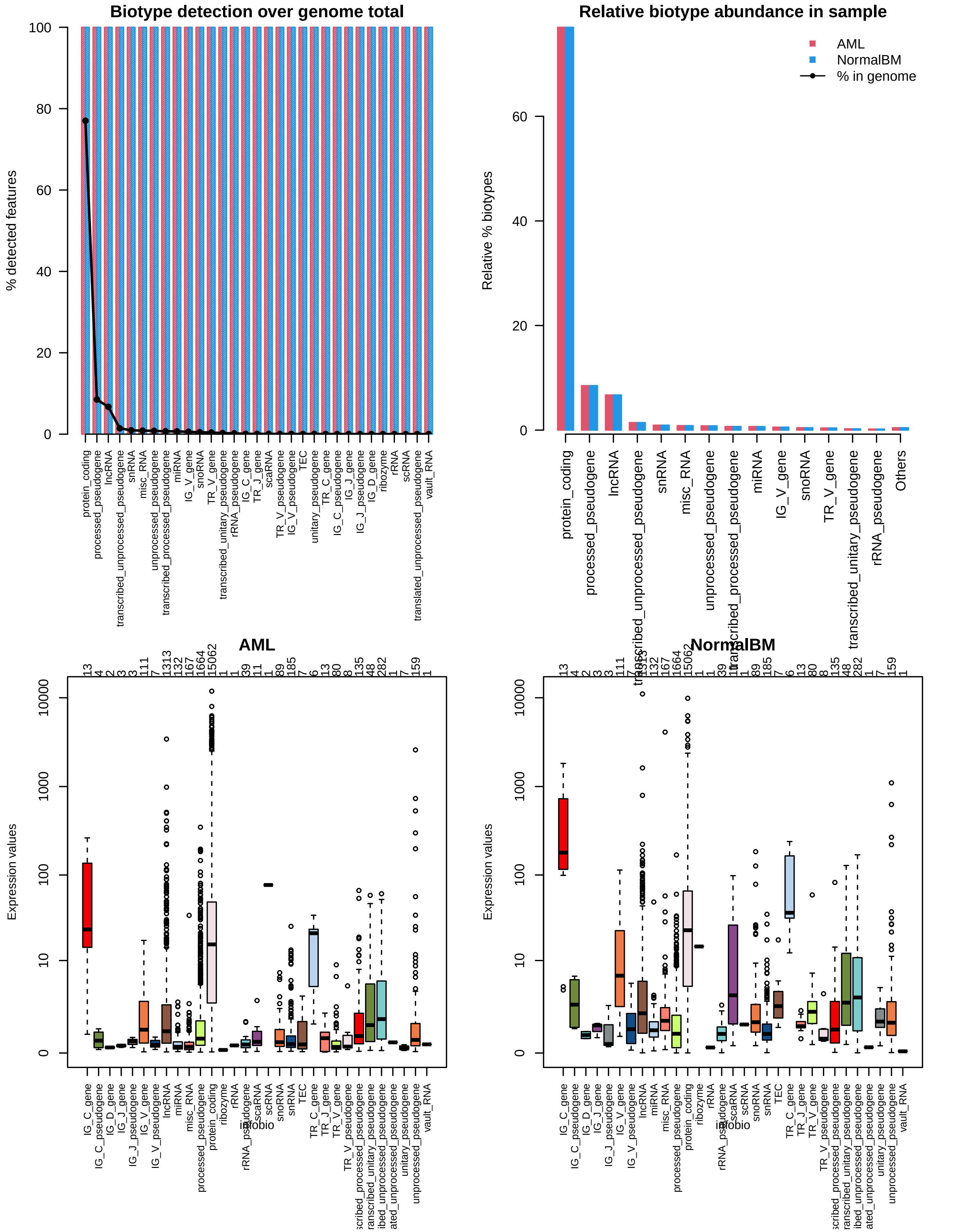

### Sequencing depth & Expression quantification

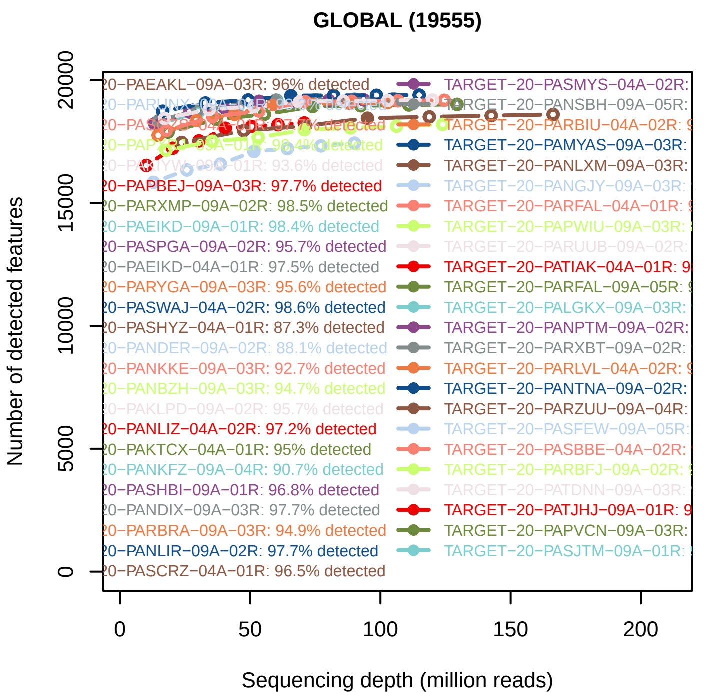


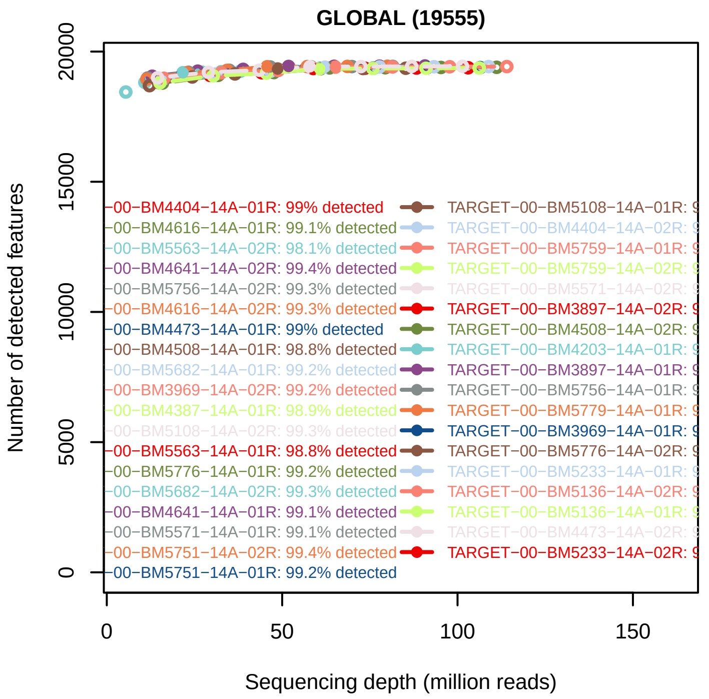


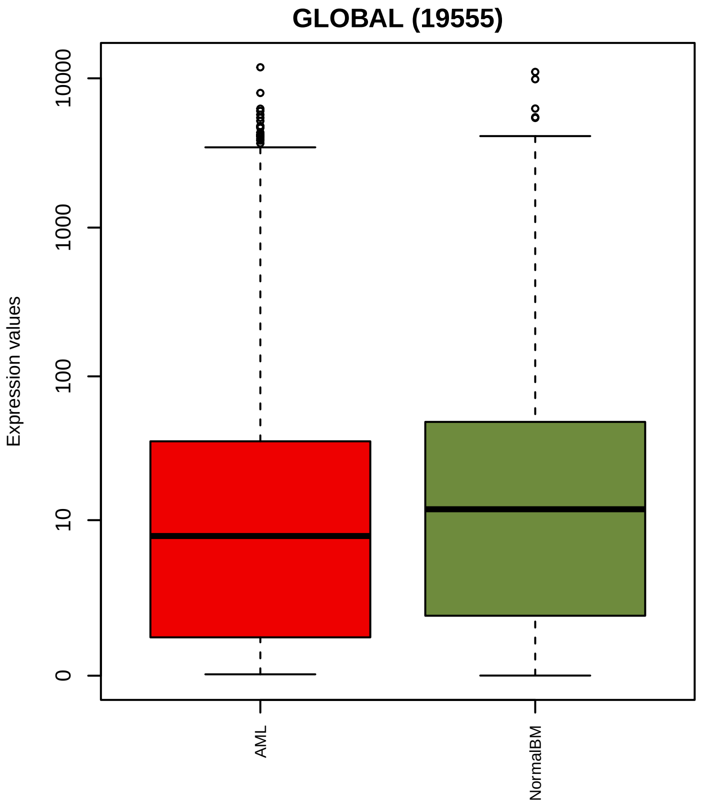


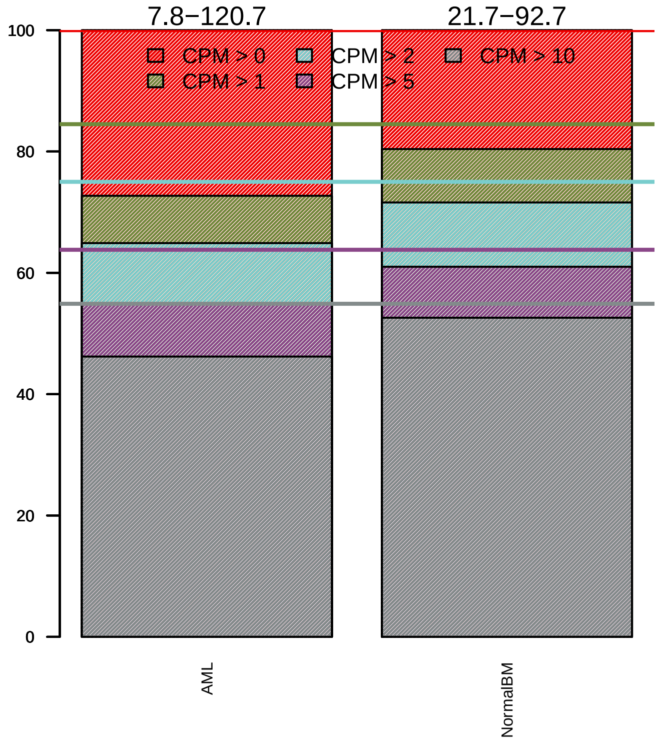

### Sequencing bias detection

#### *Diagnostic plot for feature length bias*

FAILED. At least one of the model p−values was lower than 0.05 and R2 > 70%.

Normalization for correcting length bias is recommended.

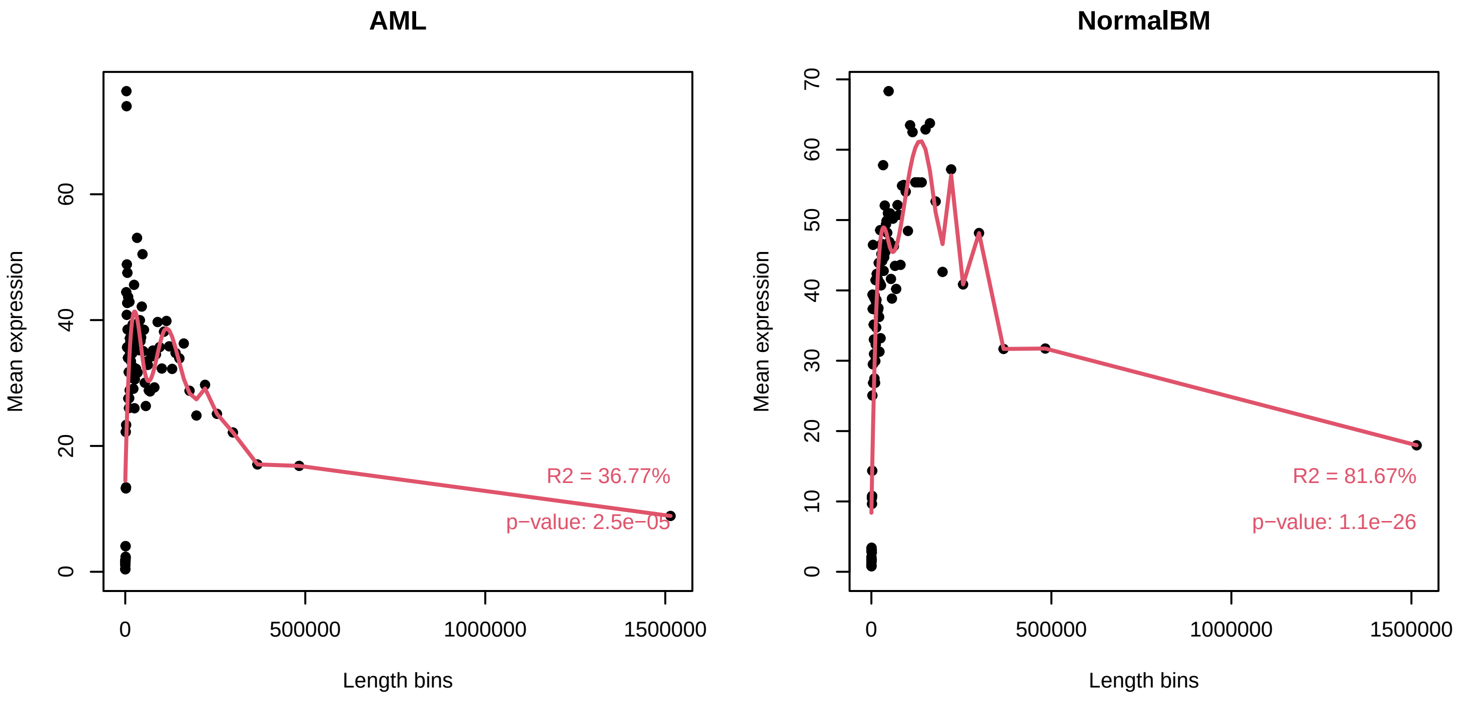


#### *Diagnostic plot for GC content bias*

WARNING. At least one of the model p−values was lower than 0.05, but R2 < 70% for at least one condition.

Normalization for correcting GC content bias could be advisable.
Plese check in the plots below the strength of the relationship between GC content and expression.

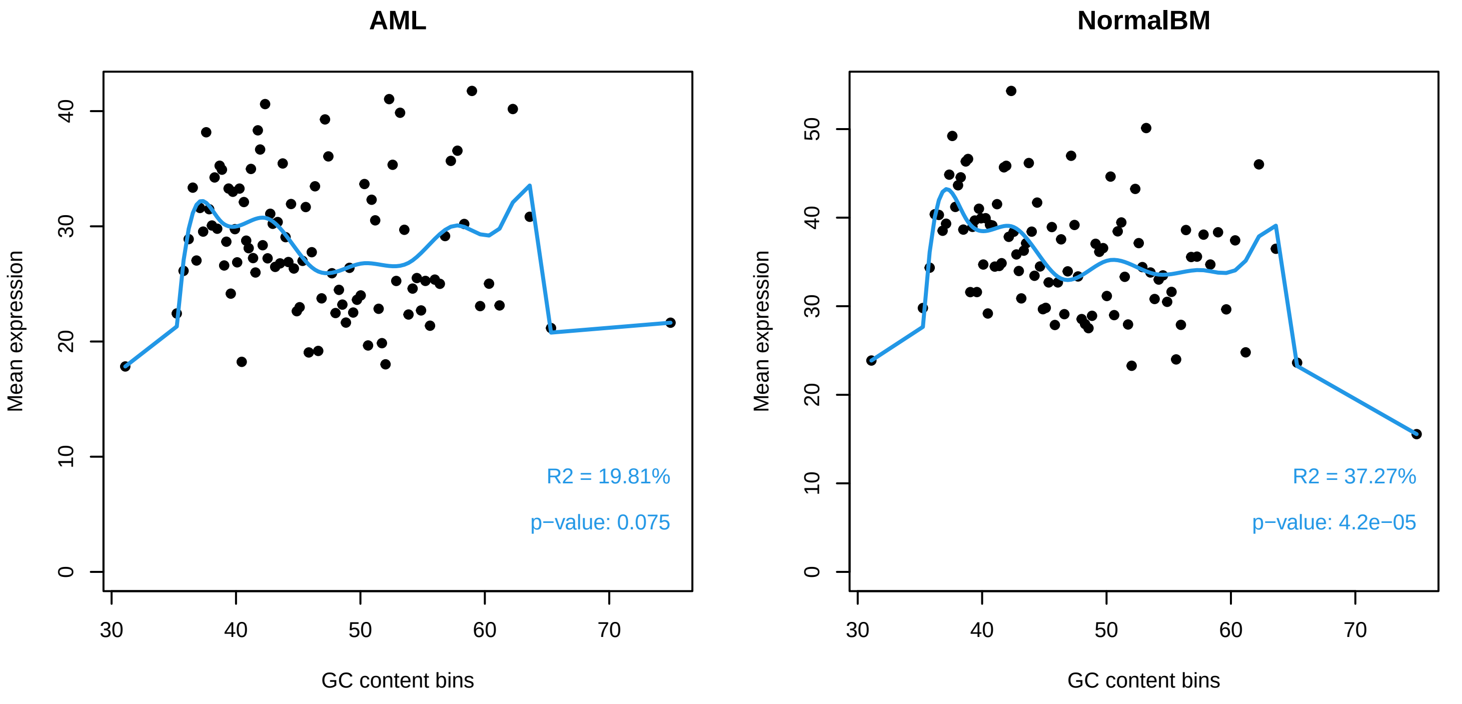

#### *Diagnostic plot for differences in RNA composition*

FAILED. There is a pair of samples with significantly different RNA composition

Normalization for correcting this bias is required.

##### Reference sample: TARGET−20−PASTUH−04A−01R

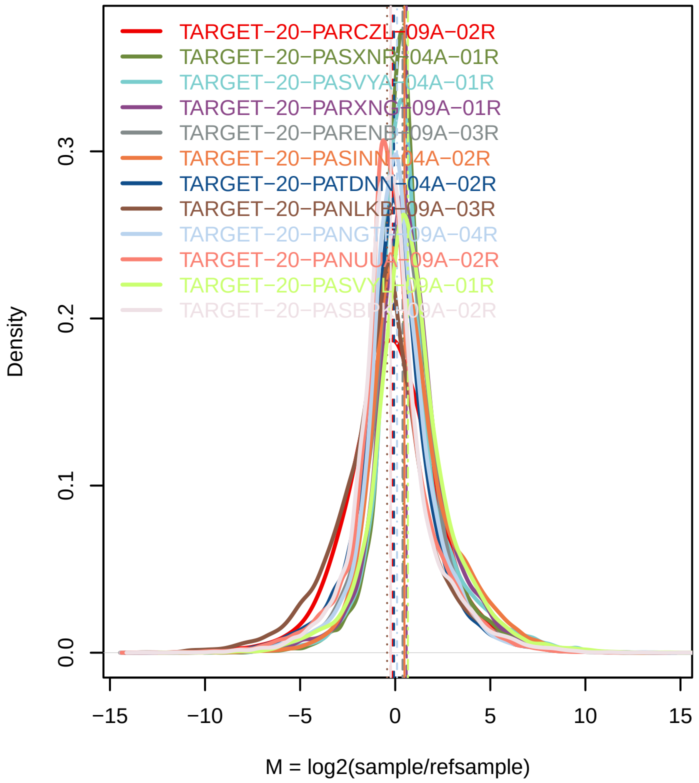


##### Confidence intervals for median of M values

| Sample | 0.01% | 99.99% | Diagnostic Test |
|---|---|---|---|
| TARGET−20−PARCZL−09A−02R | −0.2096 | −0.0782 | FAILED |
| TARGET−20−PASXNR−04A−01R | 0.4241 | 0.453 | FAILED |
| TARGET−20−PASVYA−04A−01R | 0.4396 | 0.51 | FAILED |
| TARGET−20−PARXNG−09A−01R | 0.5519 | 0.6544 | FAILED |
| TARGET−20−PARENB−09A−03R | 0.3363 | 0.4266 | FAILED |
| TARGET−20−PASINN−04A−02R | 0.4264 | 0.5241 | FAILED |
| TARGET−20−PATDNN−04A−02R | −0.1114 | −0.0189 | FAILED |
| TARGET−20−PANLKB−09A−03R | −0.4877 | −0.3609 | FAILED |
| TARGET−20−PANGTF−09A−04R | 0.0612 | 0.1346 | FAILED |
| TARGET−20−PANUUA−09A−02R | −0.2831 | −0.208 | FAILED |
| TARGET−20−PASVYL−09A−01R | 0.6215 | 0.7228 | FAILED |
| TARGET−20−PASBPK−09A−02R | −0.2718 | −0.1854 | FAILED |
| TARGET−20−PARAJX−09A−02R | −0.51 | −0.4155 | FAILED |
| TARGET−20−PANVGE−09A−02R | −0.8322 | −0.7998 | FAILED |
| TARGET−20−PASTTW−09A−01R | 0.5482 | 0.6668 | FAILED |
| TARGET−20−PARCUK−09A−03R | −0.2937 | −0.2617 | FAILED |
| TARGET−20−PASIBG−09A−01R | 0.4908 | 0.5252 | FAILED |
| TARGET−20−PARUNX−09A−01R | 0.4391 | 0.5241 | FAILED |
| TARGET−20−PASVVS−09A−03R | 0.0072 | 0.0764 | FAILED |
| TARGET−20−PASXYG−09A−05R | −0.0395 | 0.0603 | PASSED |
| TARGET−20−PARBIU−09A−03R | 0.5126 | 0.6024 | FAILED |
| TARGET−20−PAPXWI−09A−03R | −0.7951 | −0.7737 | FAILED |
| TARGET−20−PARTAL−09A−01R | 0.3943 | 0.4587 | FAILED |
| TARGET−20−PARHVK−09A−04R | 0.172 | 0.2465 | FAILED |
| TARGET−20−PASCFW−09A−01R | 0.0595 | 0.1515 | FAILED |
| TARGET−20−PAPWYK−09A−03R | −0.1309 | −0.0512 | FAILED |
| TARGET−20−PARCZL−09A−02R | 0.2247 | 0.3075 | FAILED |
| TARGET−20−PARUTH−09A−04R | −0.3891 | −0.3011 | FAILED |

### Exploratory PCA

Use this plot to see if samples are clustered according to the experimental design.

Use ARSyNseq function to correct potential batch effects.

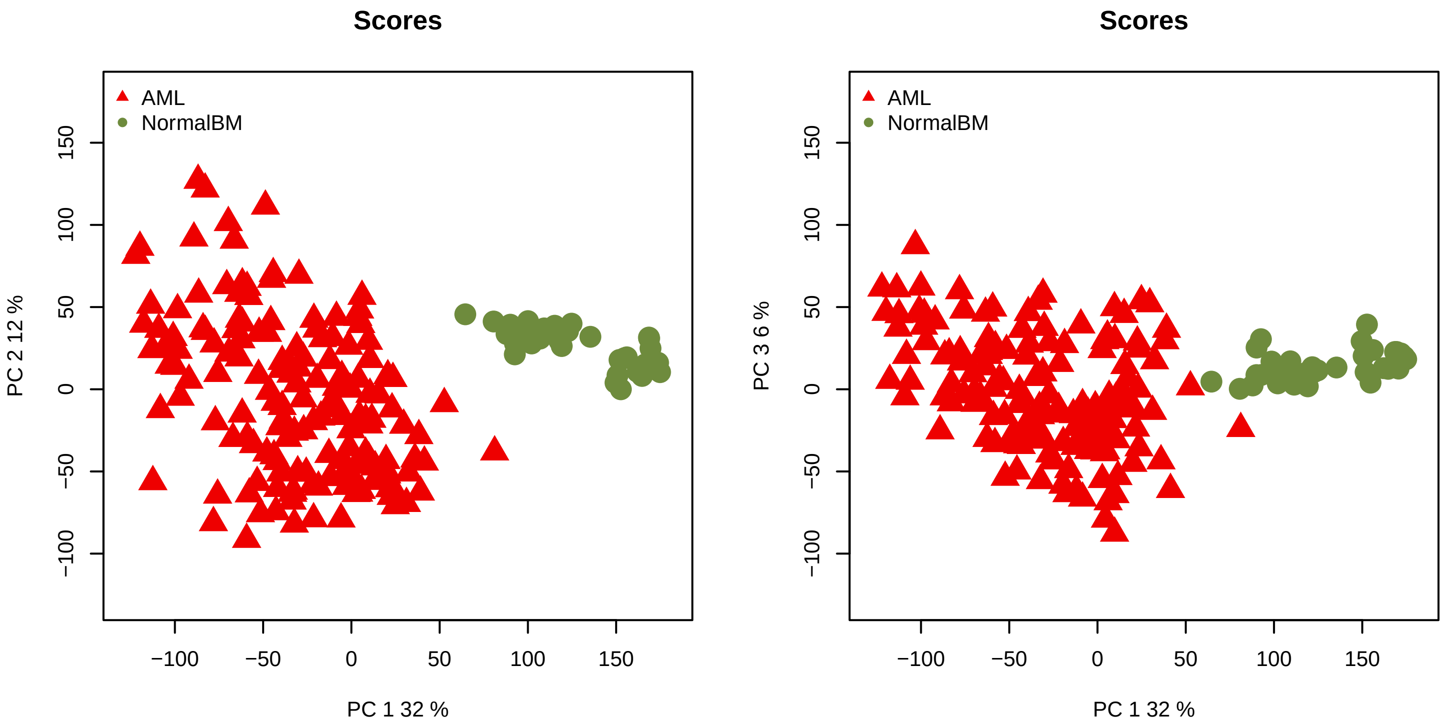
