## Supplementary material for "The network structure of hematopoietic cancers": Quality control for data pre-processing.: QCreport_BALL_After.pdf

### Quality Control of Expression Data

*Generated by NOISeq on 24 Feb 2023, 13:29:28*

#### Content

| *Plot* | *Description* |
|---|---|
| **Biotype detection** | Biotype abundance in the genome with %genes detected (counts > 0) in the sample/condition.<br>Biotype abundance within the sample/condition. |
| **Biotype expression** | Distribution of gene counts per million per biotype in sample/condition (only genes with counts > 0). |
| **Saturation** | Number of detected genes (counts > 0) per sample across different sequencing depths |
| **Expression boxplot** | Distribution of gene counts per million (all biotypes) in each sample/condition |
| **Expression barplot** | Percentage of genes with >0, >1, >2, >5 or >10 counts per million in each sample/condition. |
| **Length bias** | Mean gene expression per each length bin. Fitted curve and diagnostic test. |
| **GC content bias** | Mean gene expression per each GC content bin. Fitted curve and diagnostic test. |
| **RNA composition bias** | Density plots of log fold changes (M) between pairs of samples.<br>Confidence intervals for the median of M values. |
| **Exploratory PCA** | Principal Component Analysis score plots for PC1 vs PC2, and PC1 vs PC3. |

### Biotype detection

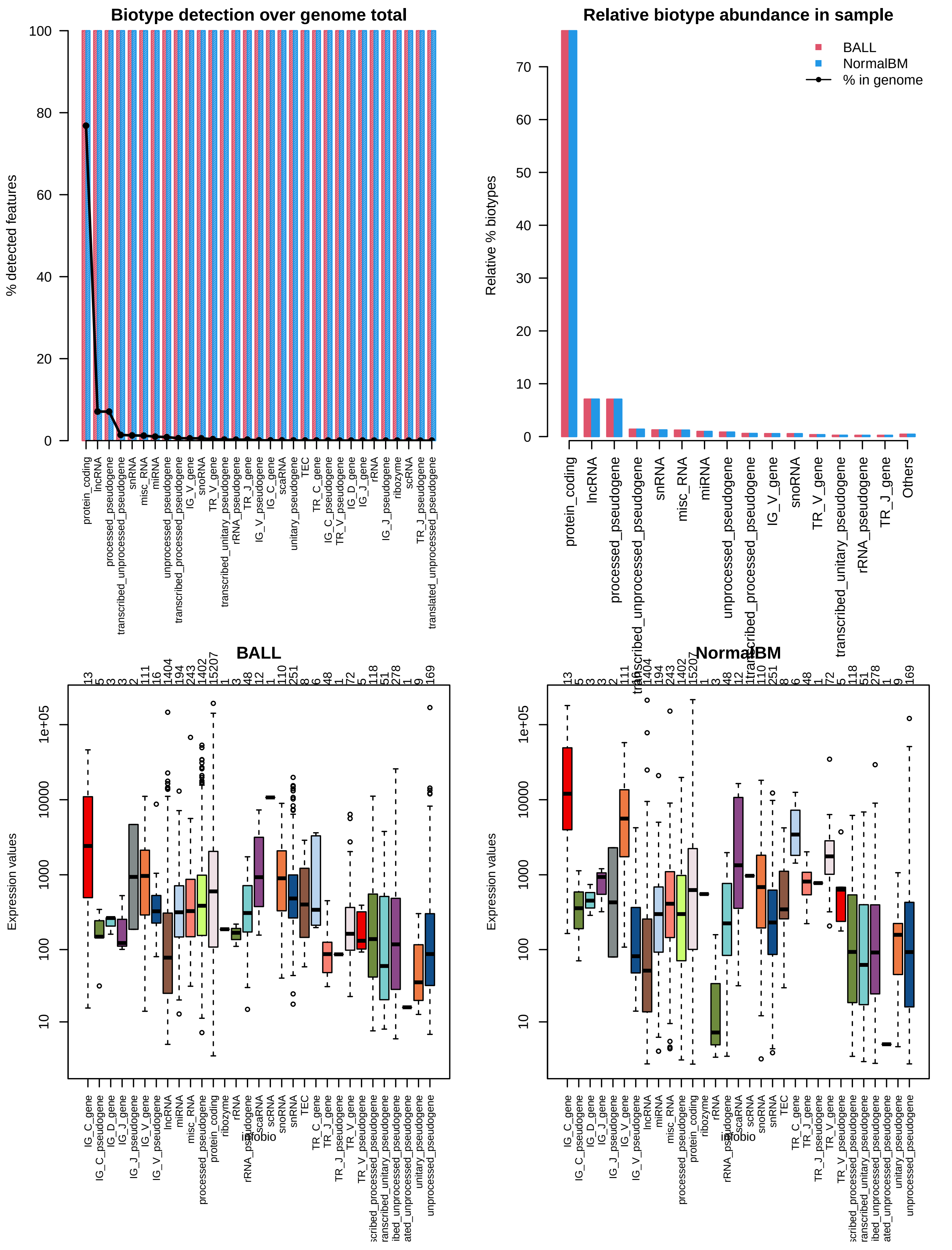

### Sequencing depth & Expression quantification

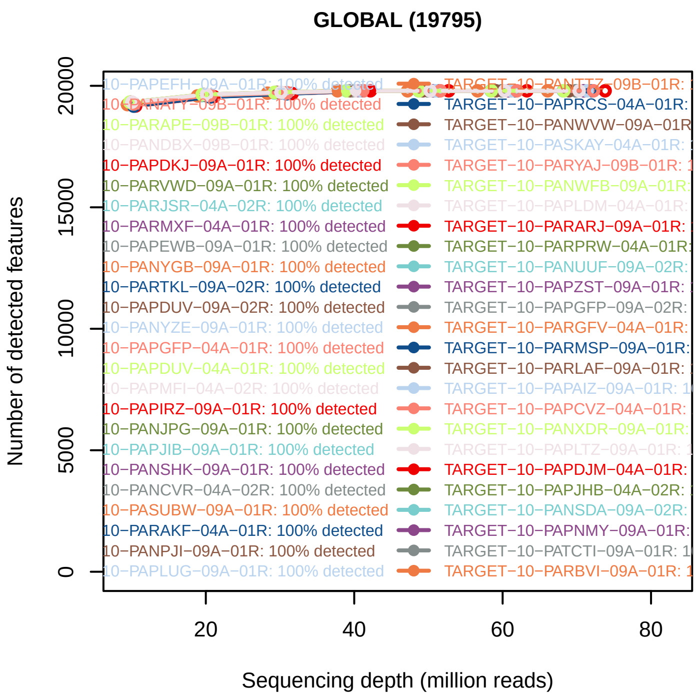


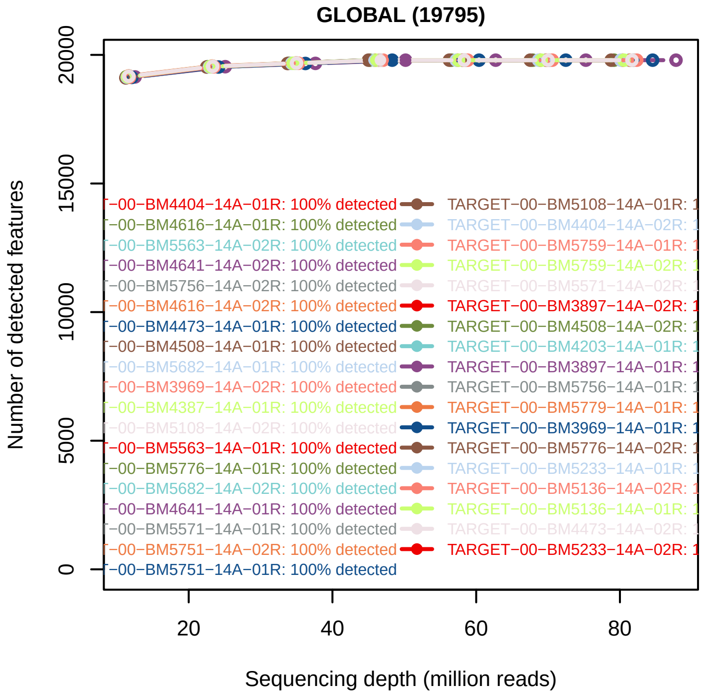


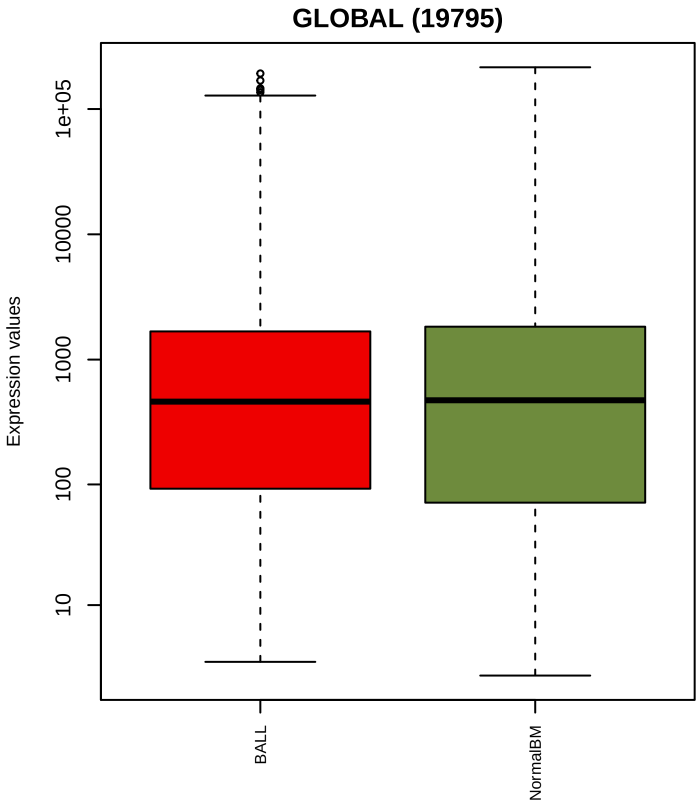


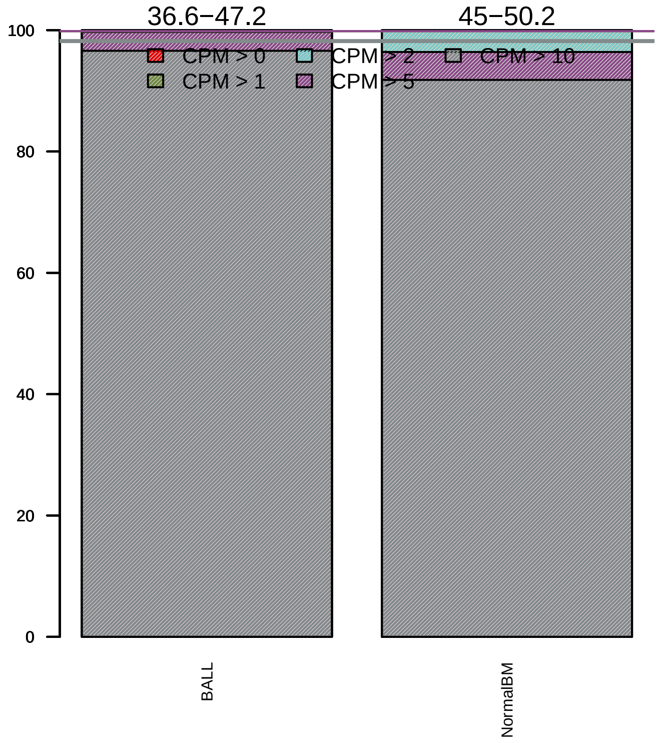

### Sequencing bias detection

#### *Diagnostic plot for feature length bias*

WARNING. At least one of the model p−values was lower than 0.05, but R2 < 70% for at least one condition.

Normalization for correcting length bias could be advisable.
Plese check in the plots below the strength of the relationship between length and expression.

**BALL**

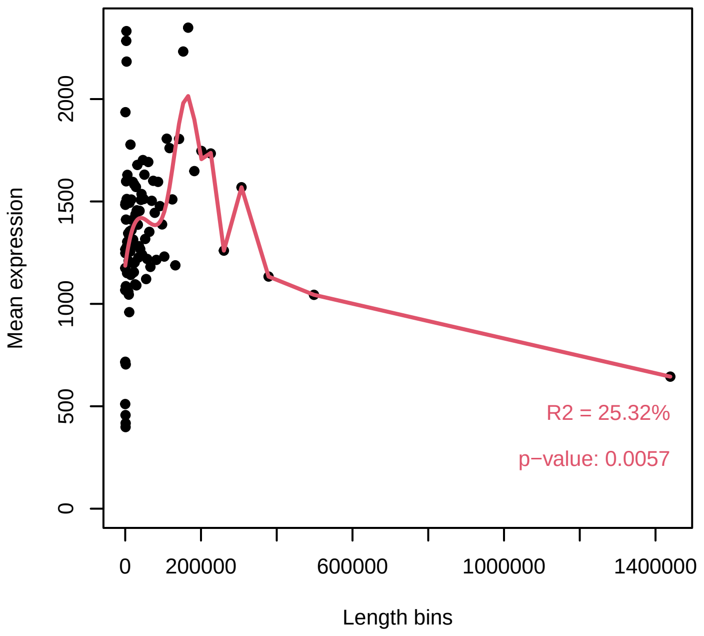


**NormalBM**

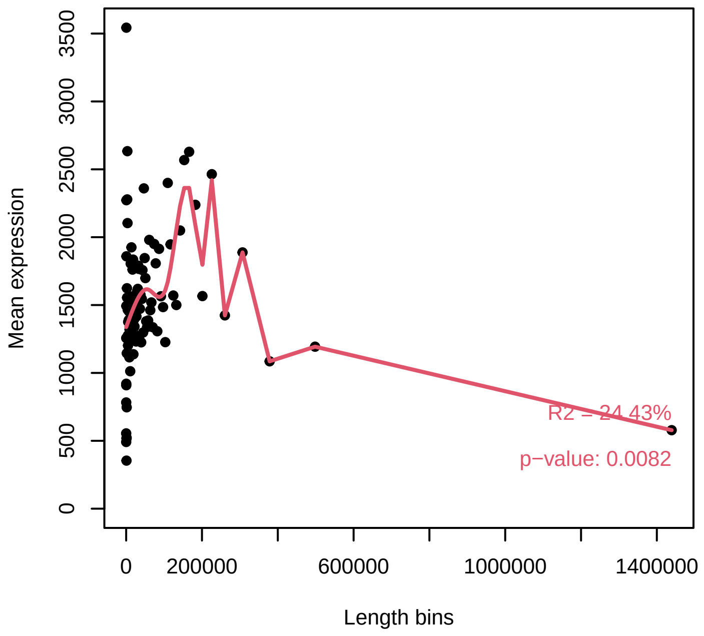


#### *Diagnostic plot for GC content bias*

PASSED. No normalization for correcting GC content bias is required.

**BALL**

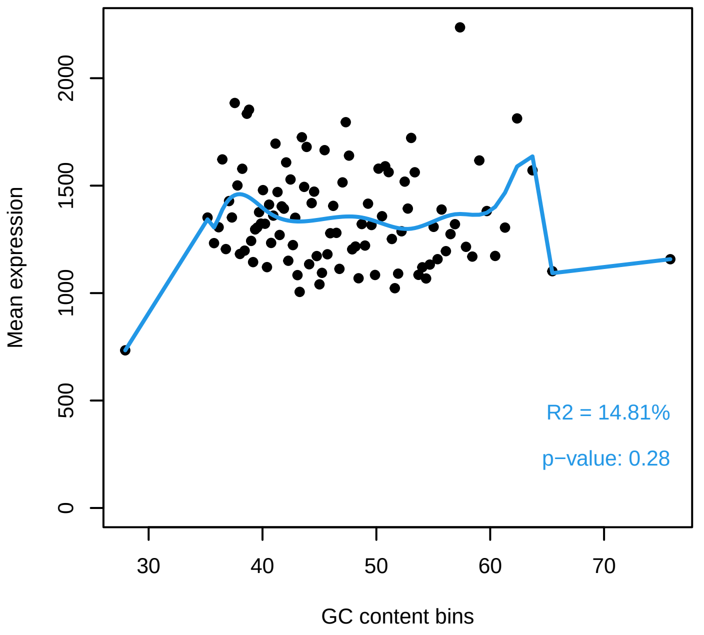


**NormalBM**

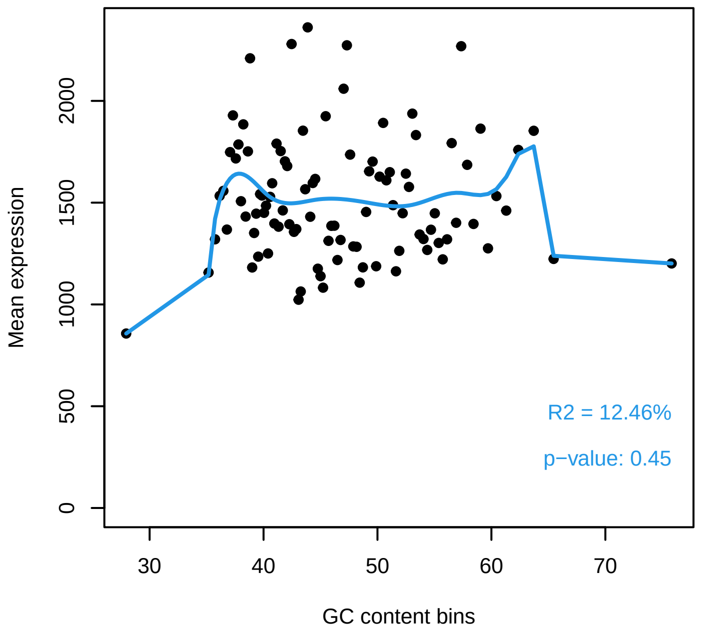

### *Diagnostic plot for differences in RNA composition*

FAILED. There is a pair of samples with significantly different RNA composition

Normalization for correcting this bias is required.

#### Reference sample: TARGET−10−PASFXA−04A−02R

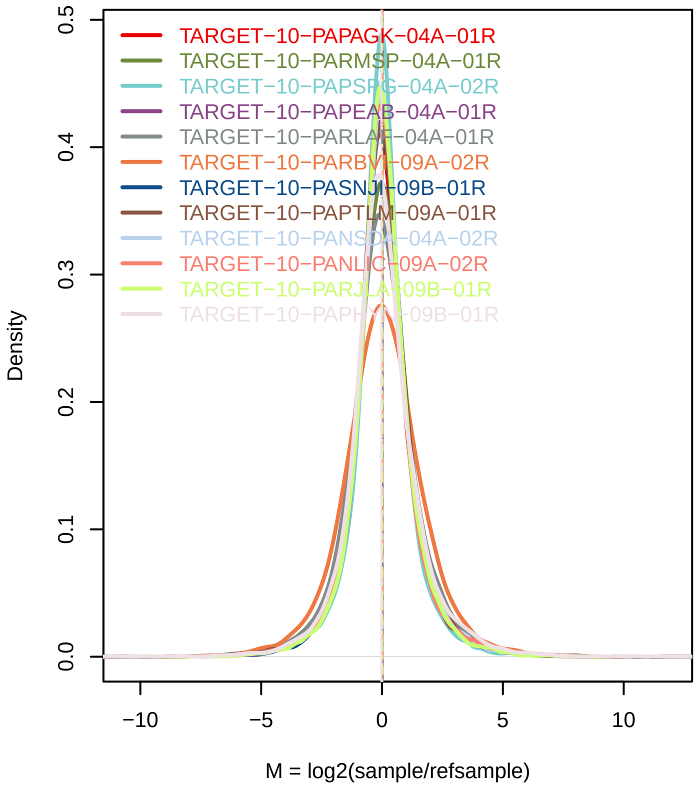


#### Confidence intervals for median of M values

| Sample | 0.01% | 99.99% | Diagnostic Test |
|---|---|---|---|
| TARGET−10−PAPAGK−04A−01R | −0.0487 | 0.0067 | PASSED |
| TARGET−10−PARMSP−04A−01R | −0.0175 | 0.0388 | PASSED |
| TARGET−10−PAPSPG−04A−02R | −0.0272 | 0.0176 | PASSED |
| TARGET−10−PAPEAB−04A−01R | −0.0241 | 0.0295 | PASSED |
| TARGET−10−PARLAF−04A−01R | −0.0549 | 0.0017 | PASSED |
| TARGET−10−PARBVI−09A−02R | −0.0171 | 0.0612 | PASSED |
| TARGET−10−PASNJH−09B−01R | −0.0124 | 0.0399 | PASSED |
| TARGET−10−PAPTLM−09A−01R | −0.0217 | 0.0243 | PASSED |
| TARGET−10−PANSDA−04A−02R | −0.0158 | 0.0383 | PASSED |
| TARGET−10−PANLIC−09A−02R | −0.0394 | 0.0103 | PASSED |
| TARGET−10−PARJLA−09B−01R | −0.0251 | 0.029 | PASSED |
| TARGET−10−PAPHYN−09B−01R | −0.0539 | 0.0088 | PASSED |
| TARGET−10−PARXCD−09B−01R | −0.097 | −0.0112 | FAILED |
| TARGET−10−PARUFL−09B−01R | −0.0431 | 0.0061 | PASSED |
| TARGET−10−PANZPJ−04A−01R | −0.0412 | 0.0172 | PASSED |
| TARGET−10−PAPIJM−09A−02R | −0.0975 | 0.0395 | PASSED |
| TARGET−10−PANKGK−09A−01R | −0.0417 | 0.0144 | PASSED |
| TARGET−10−PANEUH−09A−01R | −0.0048 | 0.0439 | PASSED |
| TARGET−10−PAPNNX−04A−01R | −0.0188 | 0.041 | PASSED |
| TARGET−10−PAPGYC−09A−01R | −0.0418 | 0.0144 | PASSED |
| TARGET−10−PASHUU−04A−02R | −0.0417 | 0.0351 | PASSED |
| TARGET−10−PAPHGD−09A−01R | −0.049 | −0.0097 | FAILED |
| TARGET−10−PANKAK−09A−01R | −0.0491 | 0.008 | PASSED |
| TARGET−10−PANWEZ−09B−01R | −0.0407 | 0.0139 | PASSED |
| TARGET−10−PAPJHR−09A−01R | −0.0393 | 0.0151 | PASSED |
| TARGET−10−PARBVI−04A−01R | −0.0317 | 0.0217 | PASSED |
| TARGET−10−PANWYH−09B−01R | −0.027 | 0.0144 | PASSED |
| TARGET−10−PAMXSP−09A−01R | −0.0097 | 0.0454 | PASSED |

### Exploratory PCA

Use this plot to see if samples are clustered according to the experimental design.

Use ARSyNseq function to correct potential batch effects.

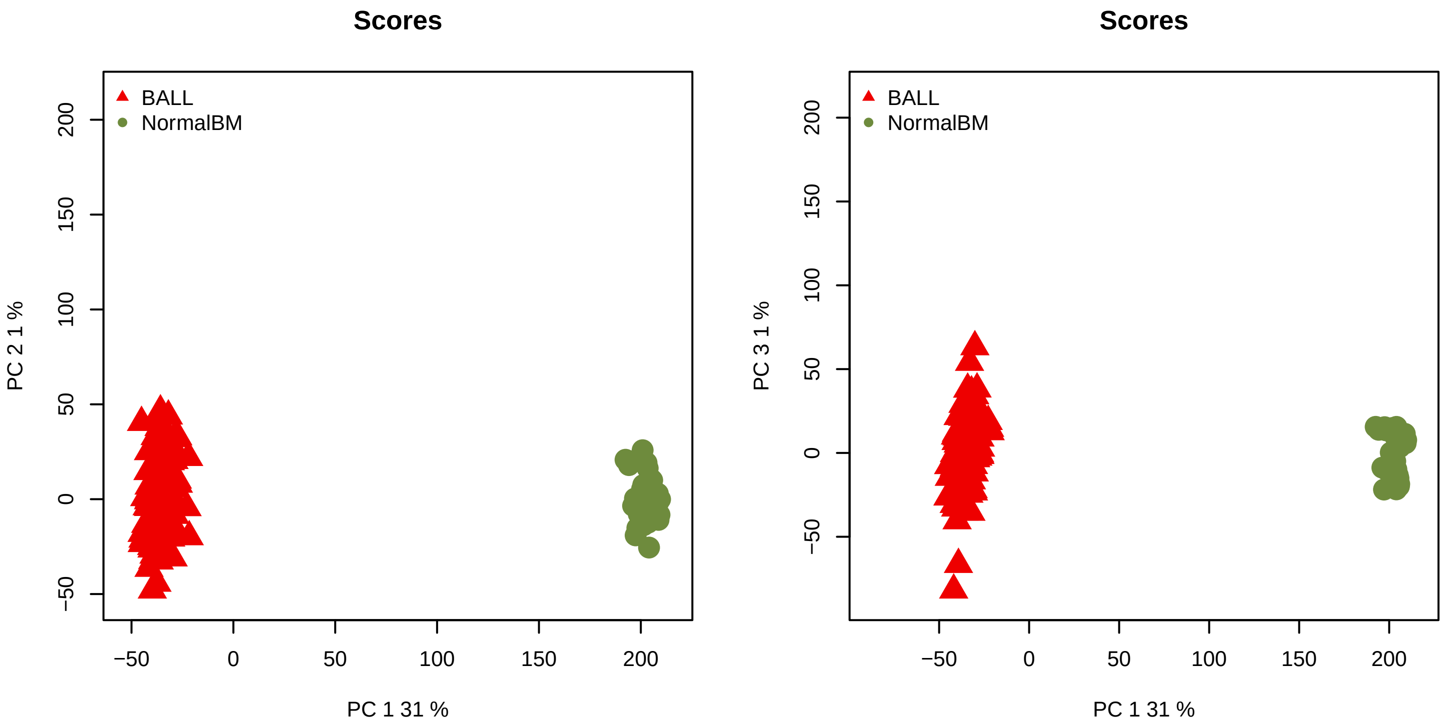
