## Supplementary material for "The network structure of hematopoietic cancers": Quality control for data pre-processing.: QCreport_BALL_Before.pdf

### Quality Control of Expression Data

*Generated by NOISeq on 24 Feb 2023, 12:58:13*

#### Content

| *Plot* | *Description* |
| --- | --- |
| **Biotype detection** | Biotype abundance in the genome with %genes detected (counts > 0) in the sample/condition.<br>Biotype abundance within the sample/condition. |
| **Biotype expression** | Distribution of gene counts per million per biotype in sample/condition (only genes with counts > 0). |
| **Saturation** | Number of detected genes (counts > 0) per sample across different sequencing depths |
| **Expression boxplot** | Distribution of gene counts per million (all biotypes) in each sample/condition |
| **Expression barplot** | Percentage of genes with >0, >1, >2, >5 or >10 counts per million in each sample/condition. |
| **Length bias** | Mean gene expression per each length bin. Fitted curve and diagnostic test. |
| **GC content bias** | Mean gene expression per each GC content bin. Fitted curve and diagnostic test. |
| **RNA composition bias** | Density plots of log fold changes (M) between pairs of samples.<br>Confidence intervals for the median of M values. |
| **Exploratory PCA** | Principal Component Analysis score plots for PC1 vs PC2, and PC1 vs PC3. |

### Biotype detection

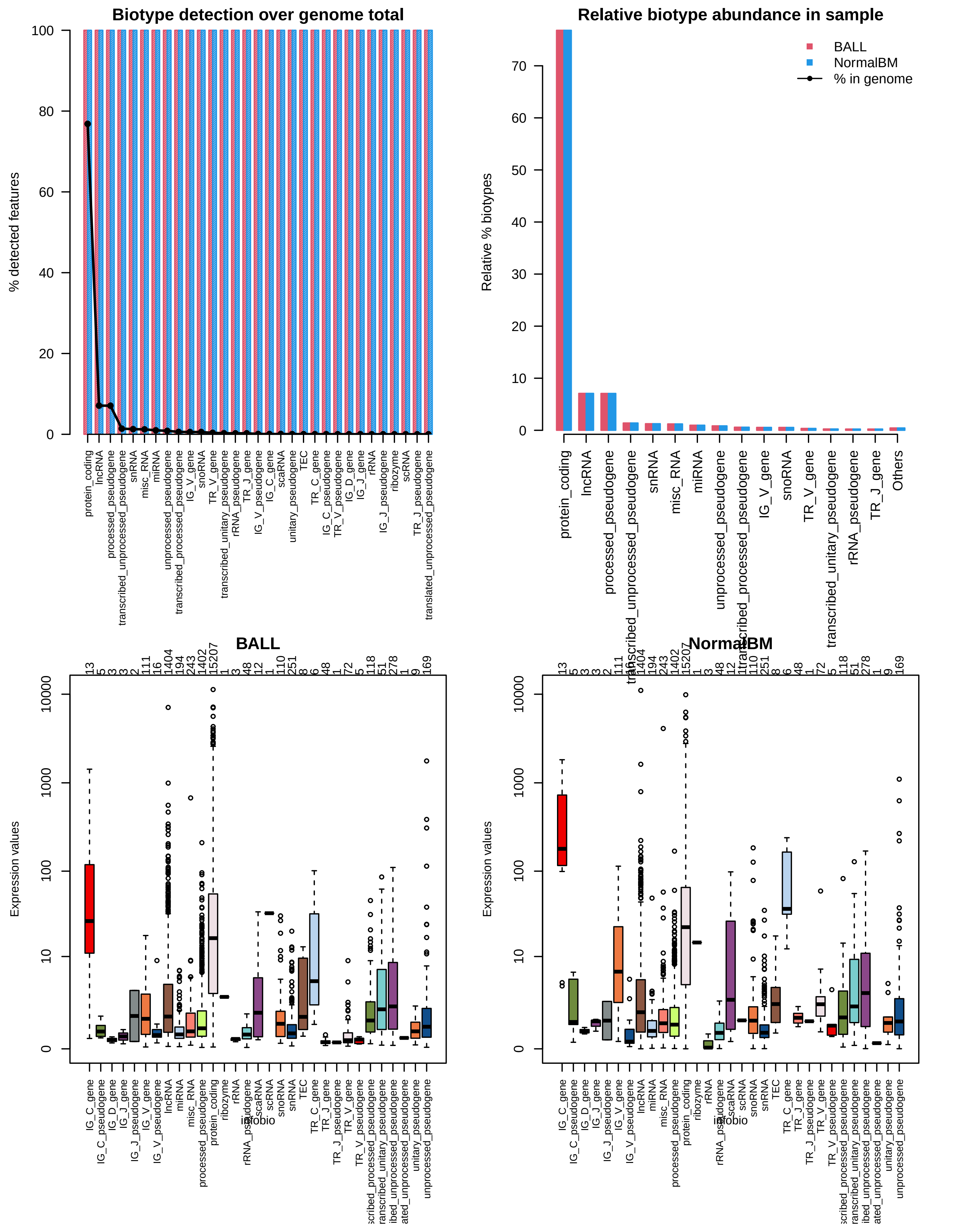

### Sequencing depth & Expression quantification

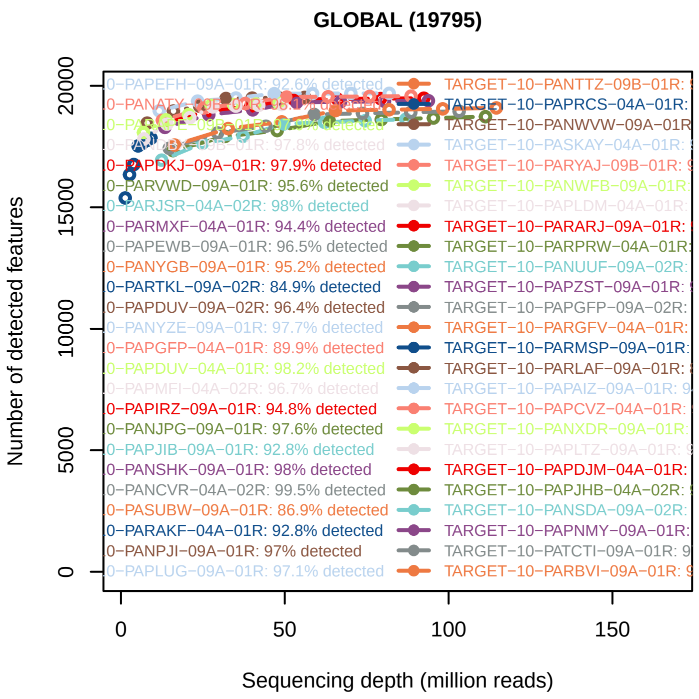


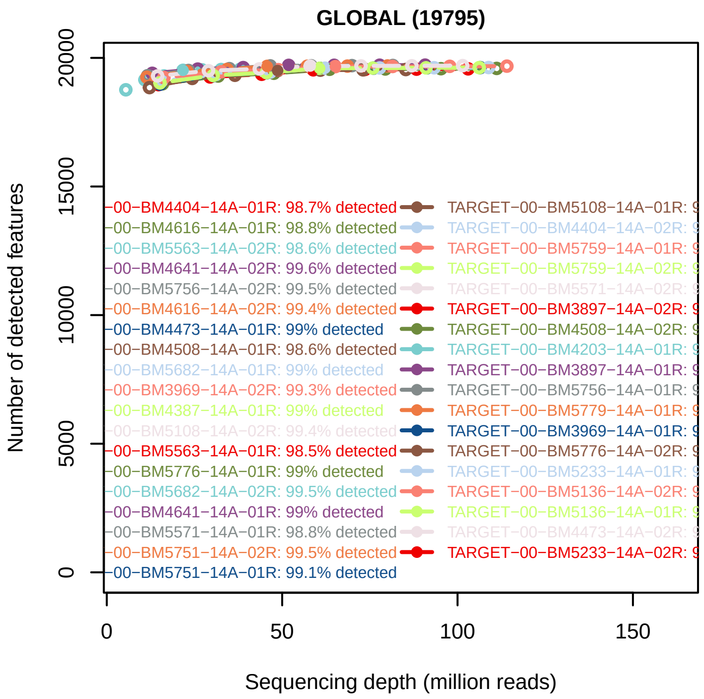


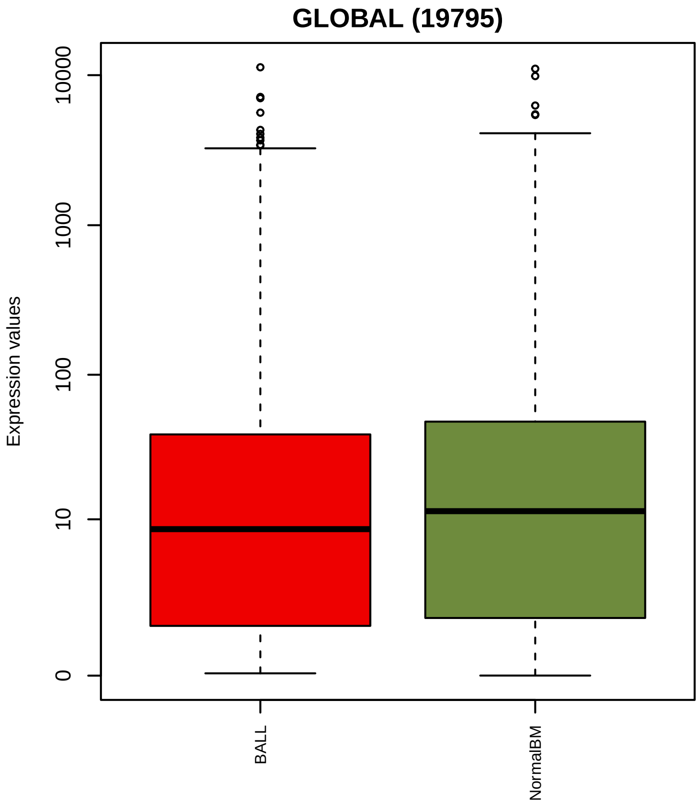


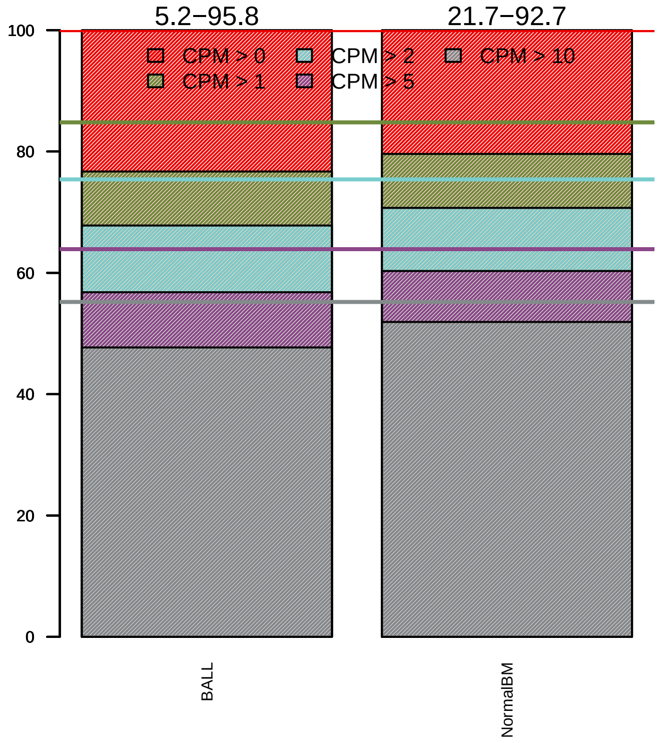

### Sequencing bias detection

#### *Diagnostic plot for feature length bias*

FAILED. At least one of the model p−values was lower than 0.05 and R2 > 70%.

Normalization for correcting length bias is recommended.

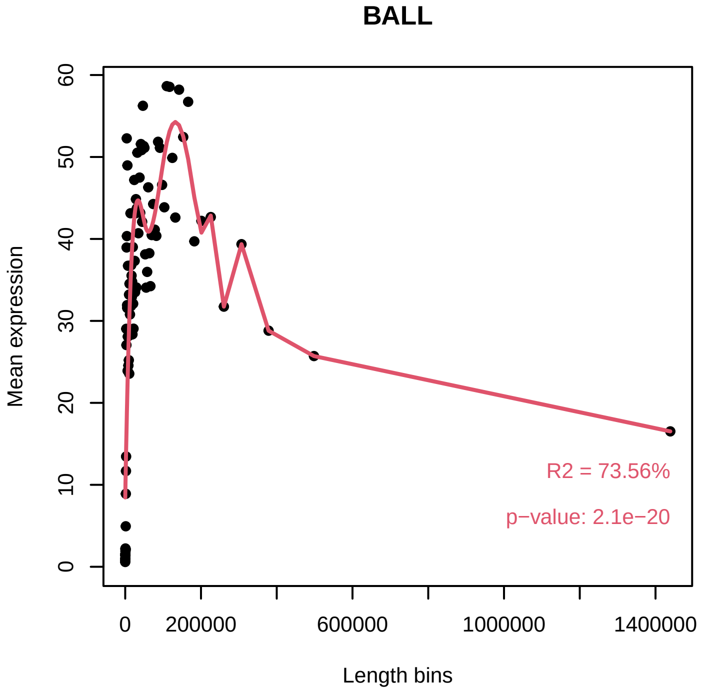


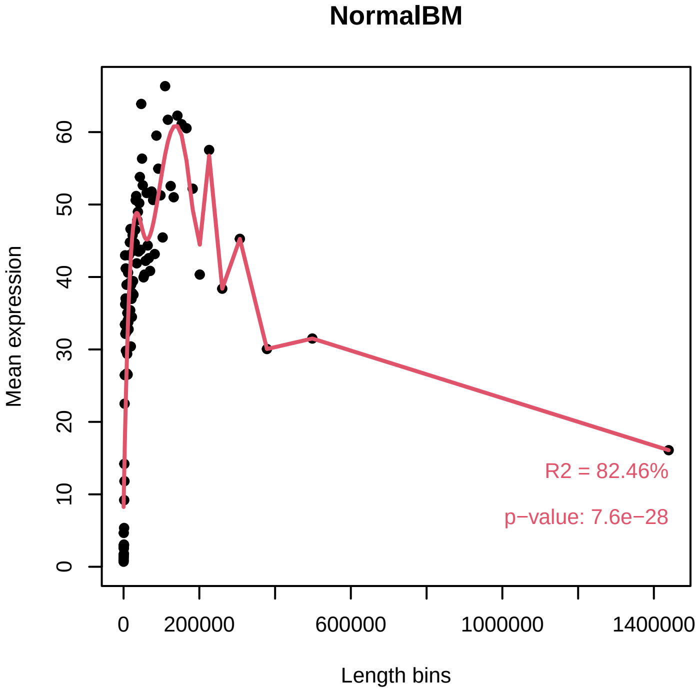


#### *Diagnostic plot for GC content bias*

WARNING. At least one of the model p−values was lower than 0.05, but R2 < 70% for at least one condition.

Normalization for correcting GC content bias could be advisable.
Plese check in the plots below the strength of the relationship between GC content and expression.

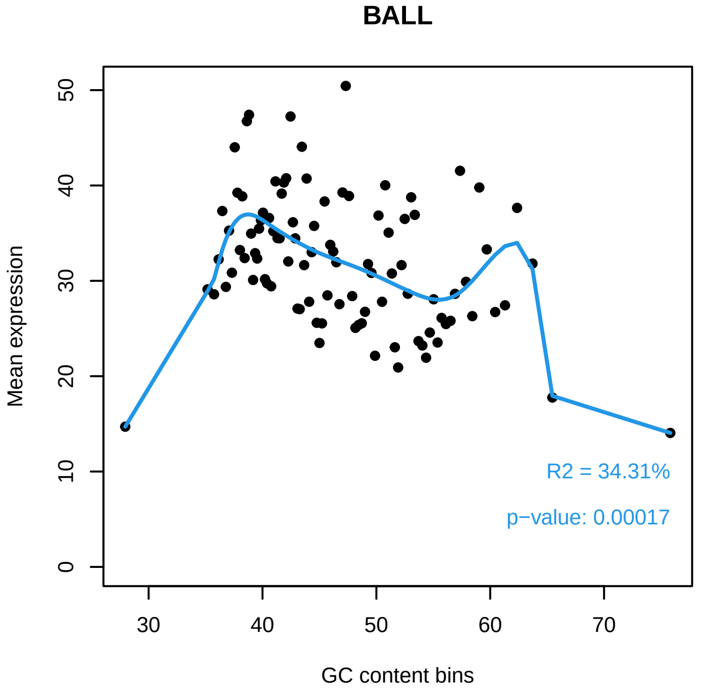


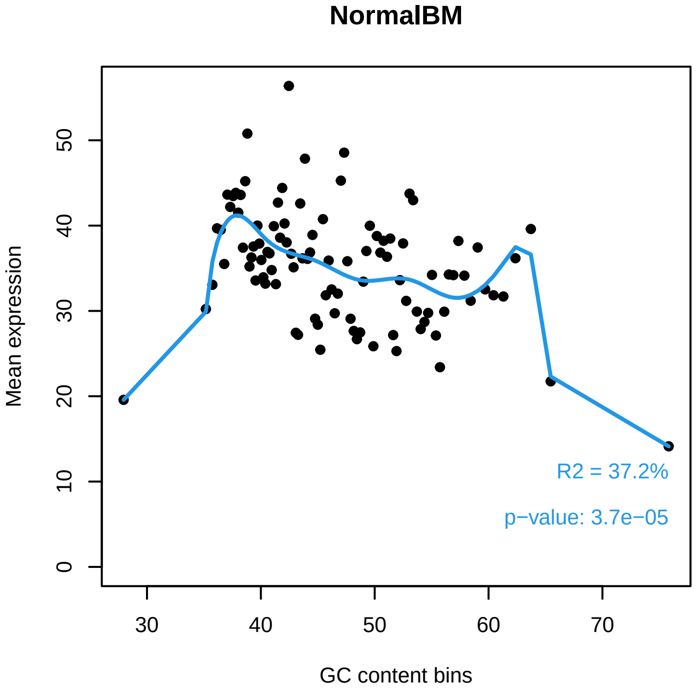

#### *Diagnostic plot for differences in RNA composition*

FAILED. There is a pair of samples with significantly different RNA composition

Normalization for correcting this bias is required.

**Reference sample: TARGET−10−PASFXA−04A−02R**

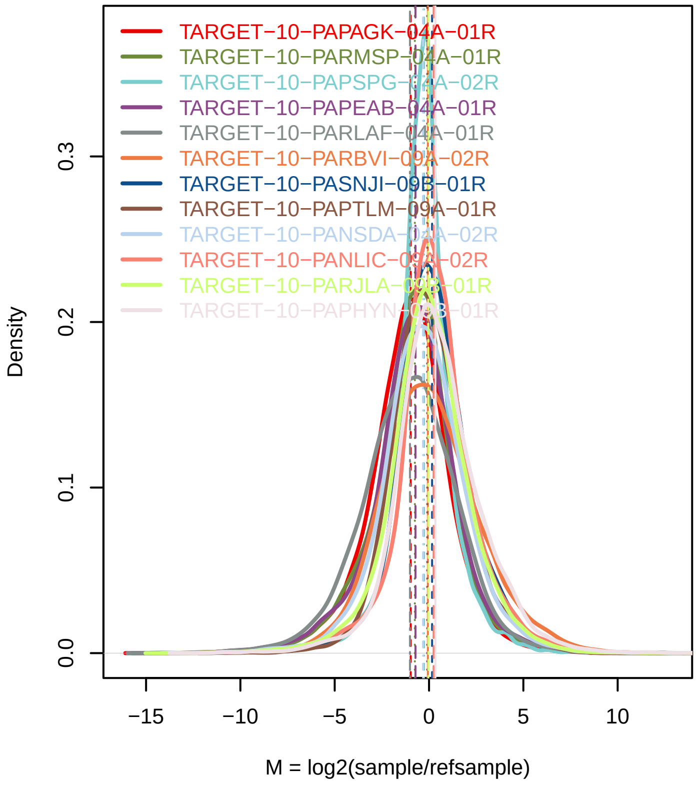


**Confidence intervals for median of M values**

| Sample | 0.01% | 99.99% | Diagnostic Test |
|---|---|---|---|
| TARGET−10−PAPAGK−04A−01R | −1.0202 | −0.9296 | FAILED |
| TARGET−10−PARMSP−04A−01R | −0.8002 | −0.6904 | FAILED |
| TARGET−10−PAPSPG−04A−02R | −0.2897 | −0.2364 | FAILED |
| TARGET−10−PAPEAB−04A−01R | −0.7662 | −0.665 | FAILED |
| TARGET−10−PARLAF−04A−01R | −1.0394 | −0.935 | FAILED |
| TARGET−10−PARBVI−09A−02R | −0.0135 | 0.0101 | PASSED |
| TARGET−10−PASNJI−09B−01R | 0.1968 | 0.2124 | FAILED |
| TARGET−10−PAPTLM−09A−01R | −0.1595 | −0.0506 | FAILED |
| TARGET−10−PANSDA−04A−02R | −0.3704 | −0.2641 | FAILED |
| TARGET−10−PANLIC−09A−02R | 0.1964 | 0.289 | FAILED |
| TARGET−10−PARJLA−09B−01R | −0.0864 | 0.0109 | PASSED |
| TARGET−10−PAPHYN−09B−01R | 0.2838 | 0.408 | FAILED |
| TARGET−10−PARXCD−09B−01R | −0.0493 | 0.0795 | PASSED |
| TARGET−10−PARUFL−09B−01R | 0.4921 | 0.5867 | FAILED |
| TARGET−10−PANZPJ−09A−01R | −0.8951 | −0.7702 | FAILED |
| TARGET−10−PAPIJM−09A−02R | 0.1907 | 0.2817 | FAILED |
| TARGET−10−PANKGK−09A−01R | −0.8638 | −0.7474 | FAILED |
| TARGET−10−PANEUH−09A−01R | −0.6491 | −0.5211 | FAILED |
| TARGET−10−PAPNNX−04A−01R | −1.246 | −1.1437 | FAILED |
| TARGET−10−PAPGYC−09A−01R | −0.1079 | −0.0851 | FAILED |
| TARGET−10−PASHUI−09A−02R | 0.0425 | 0.1208 | FAILED |
| TARGET−10−PAPHGD−09A−01R | −0.1853 | −0.0957 | FAILED |
| TARGET−10−PANKAK−09A−01R | −0.3041 | −0.2059 | FAILED |
| TARGET−10−PANWEZ−09B−01R | −0.1298 | −0.0034 | FAILED |
| TARGET−10−PAPJHR−09A−01R | −0.6802 | −0.5684 | FAILED |
| TARGET−10−PARBVI−04A−01R | −1.4418 | −1.3825 | FAILED |
| TARGET−10−PANWYH−09B−01R | 0.4925 | 0.6049 | FAILED |
| TARGET−10−PAMXSP−09A−01R | −0.4939 | −0.3836 | FAILED |

### Exploratory PCA

Use this plot to see if samples are clustered according to the experimental design.

Use ARSyNseq function to correct potential batch effects.

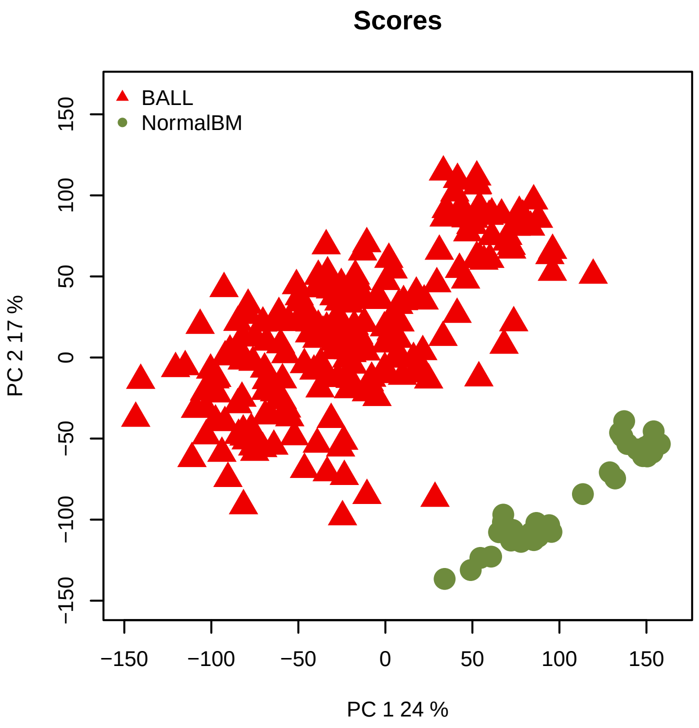


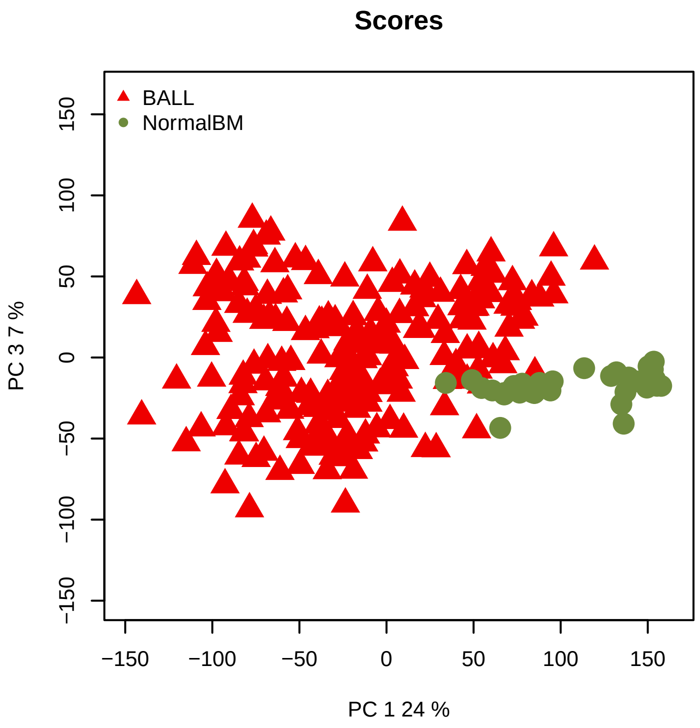
