## Supplementary material for "The network structure of hematopoietic cancers": Quality control for data pre-processing.: QCreport_MM_After.pdf

### Quality Control of Expression Data

*Generated by NOISeq on 24 Feb 2023, 13:24:42*

#### Content

| *Plot* | *Description* |
|---|---|
| **Biotype detection** | Biotype abundance in the genome with %genes detected (counts > 0) in the sample/condition.<br>Biotype abundance within the sample/condition. |
| **Biotype expression** | Distribution of gene counts per million per biotype in sample/condition (only genes with counts > 0). |
| **Saturation** | Number of detected genes (counts > 0) per sample across different sequencing depths |
| **Expression boxplot** | Distribution of gene counts per million (all biotypes) in each sample/condition |
| **Expression barplot** | Percentage of genes with >0, >1, >2, >5 or >10 counts per million in each sample/condition. |
| **Length bias** | Mean gene expression per each length bin. Fitted curve and diagnostic test. |
| **GC content bias** | Mean gene expression per each GC content bin. Fitted curve and diagnostic test. |
| **RNA composition bias** | Density plots of log fold changes (M) between pairs of samples.<br>Confidence intervals for the median of M values. |
| **Exploratory PCA** | Principal Component Analysis score plots for PC1 vs PC2, and PC1 vs PC3. |

### Biotype detection

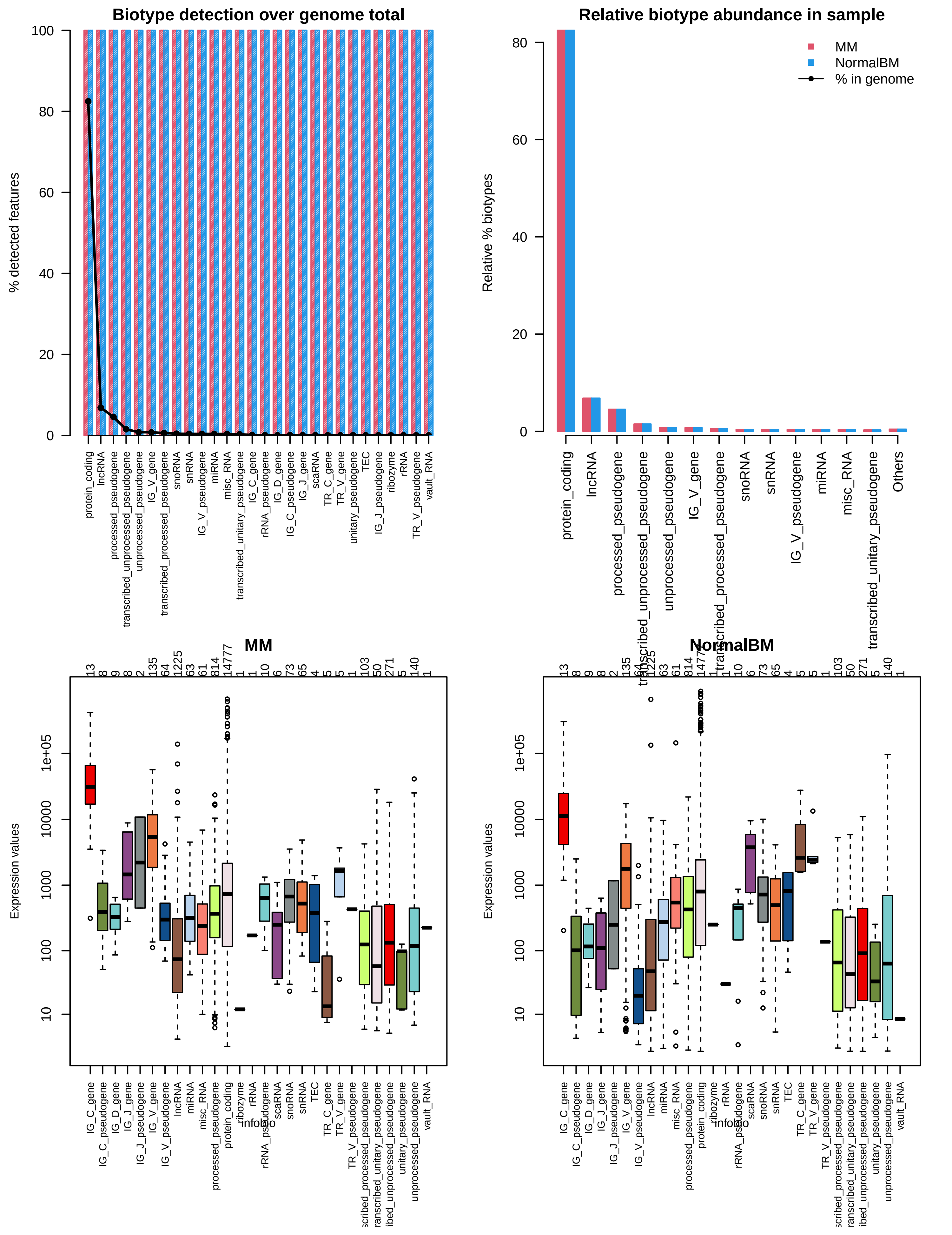

### Sequencing depth & Expression quantification

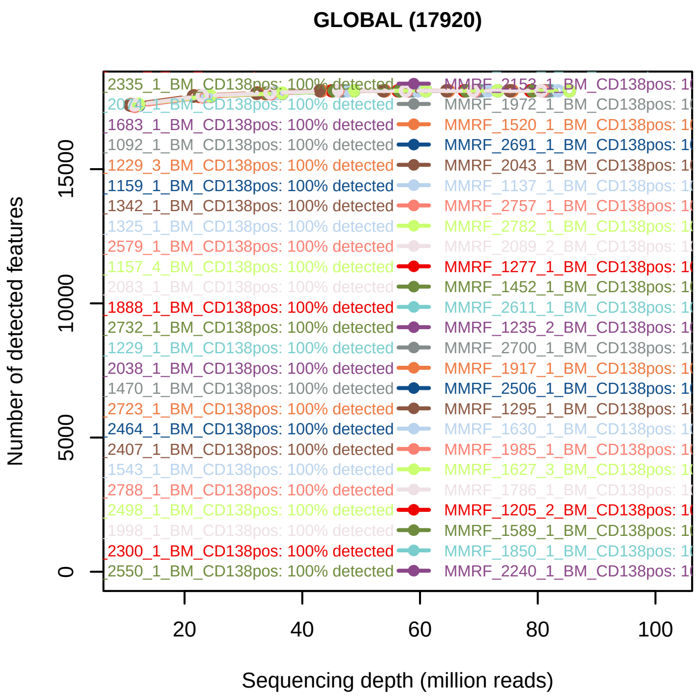


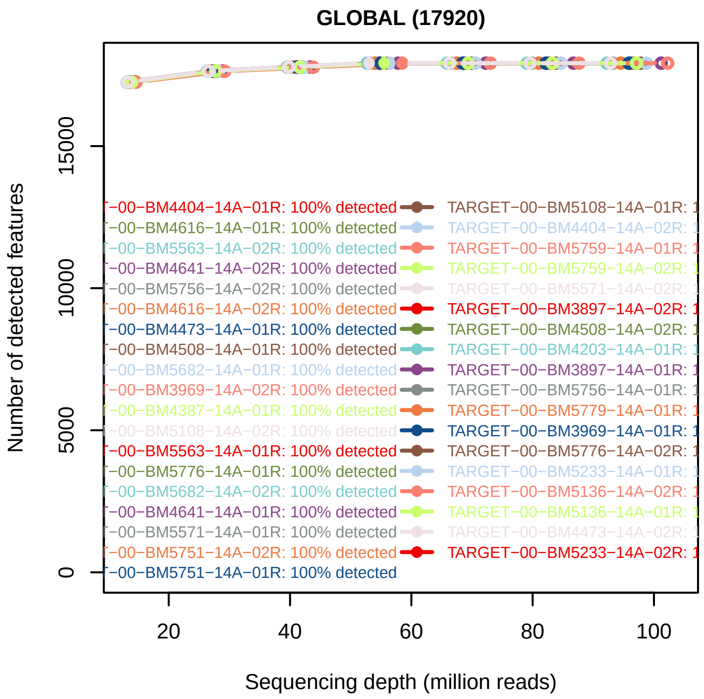


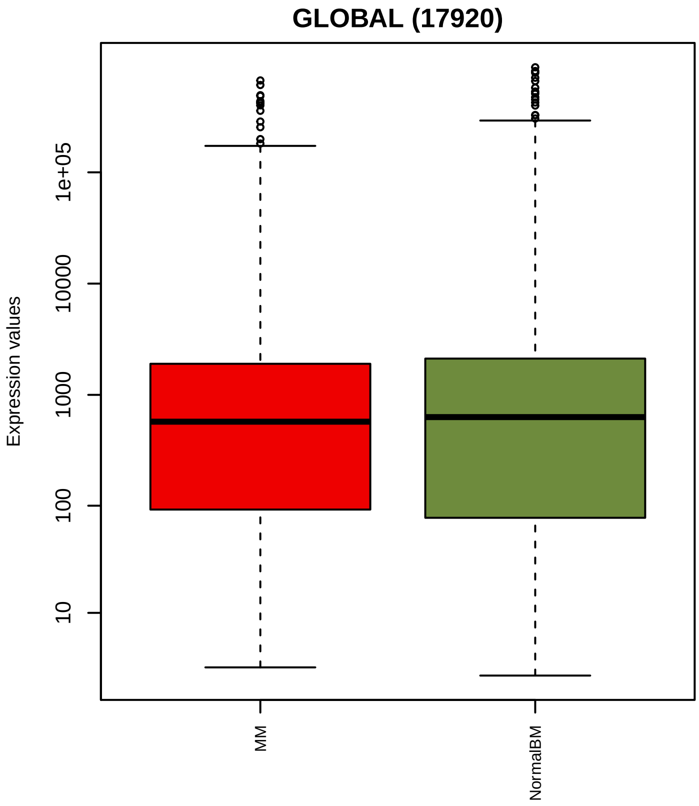


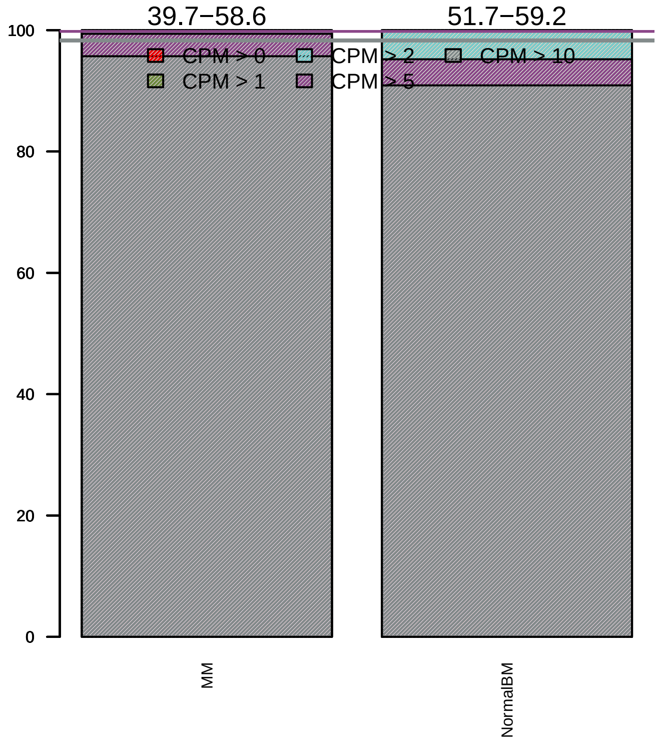

### Sequencing bias detection

#### *Diagnostic plot for feature length bias*

WARNING. At least one of the model p−values was lower than 0.05, but R2 < 70% for at least one condition.

Normalization for correcting length bias could be advisable.
Plese check in the plots below the strength of the relationship between length and expression.

**MM**

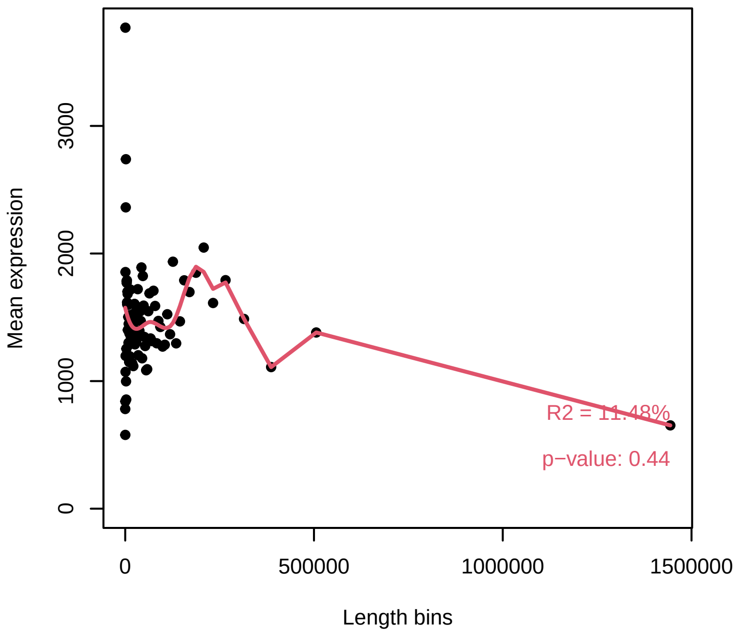


**NormalBM**

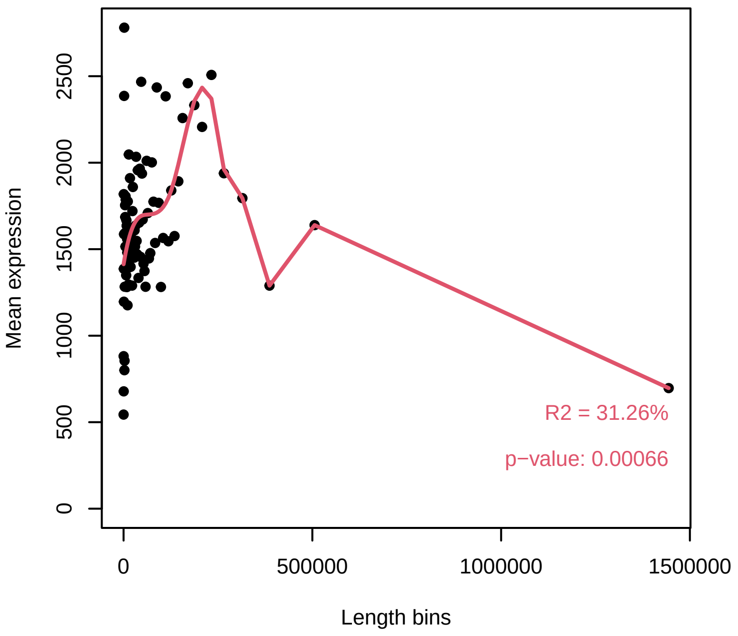


#### *Diagnostic plot for GC content bias*

PASSED. No normalization for correcting GC content bias is required.

**MM**

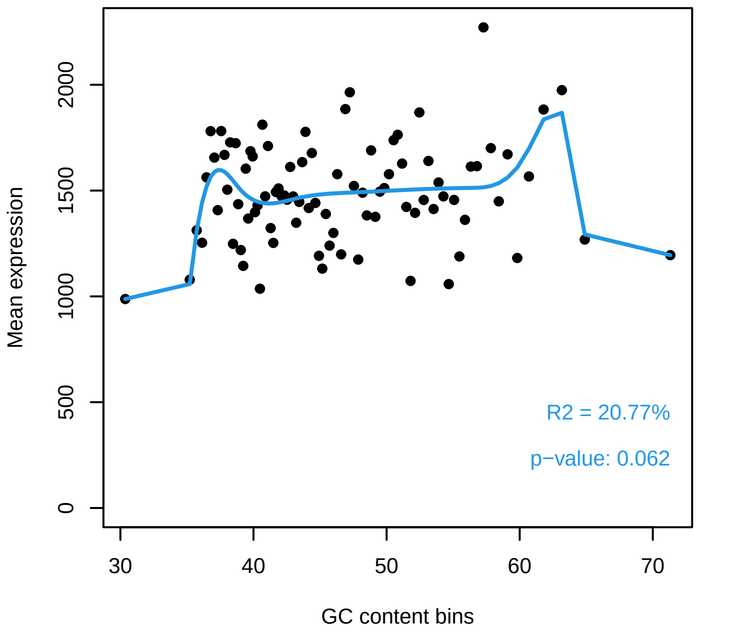


**NormalBM**

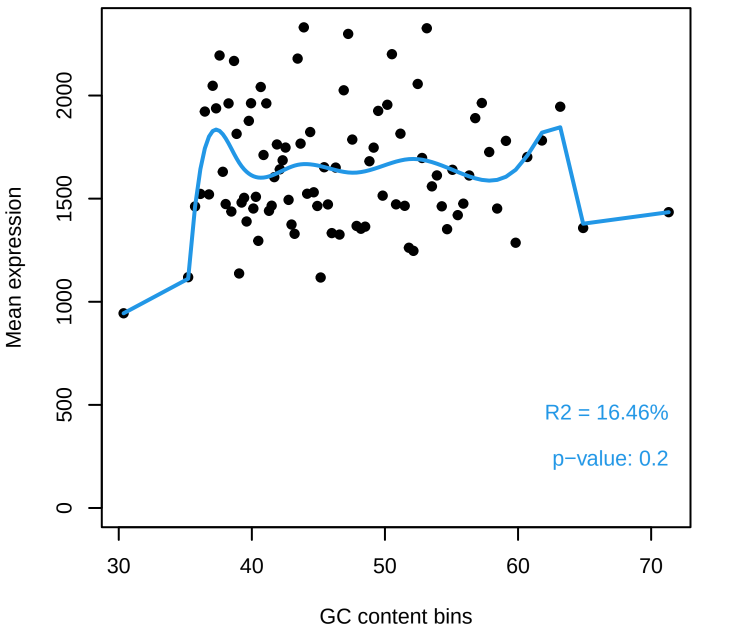

#### *Diagnostic plot for differences in RNA composition*

FAILED. There is a pair of samples with significantly different RNA composition

Normalization for correcting this bias is required.

**Reference sample: MMRF_2173_1_BM_CD138pos**

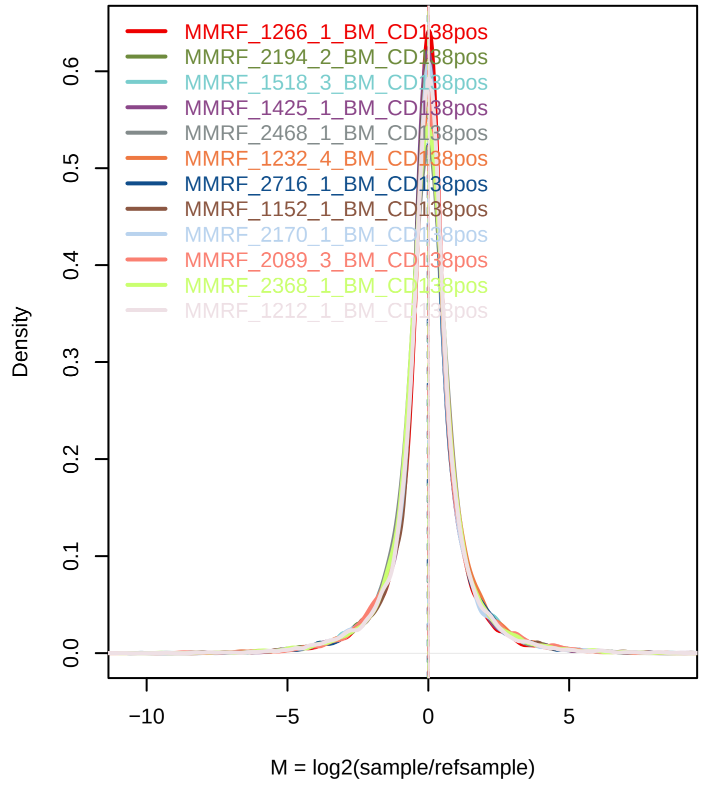


**Confidence intervals for median of M values**

| Sample | 0% | 100% | Diagnostic Test |
|---|---|---|---|
| MMRF_1266_1_BM_CD138pos | 0.0015 | 0.0425 | FAILED |
| MMRF_2194_2_BM_CD138pos | −0.0117 | 0.0251 | PASSED |
| MMRF_1518_3_BM_CD138pos | −0.031 | 0.0159 | PASSED |
| MMRF_1425_1_BM_CD138pos | −0.0207 | 0.0143 | PASSED |
| MMRF_2468_1_BM_CD138pos | −0.0218 | 0.0249 | PASSED |
| MMRF_1232_4_BM_CD138pos | −0.0032 | 0.0347 | PASSED |
| MMRF_2716_1_BM_CD138pos | −0.039 | −0.0029 | FAILED |
| MMRF_1152_1_BM_CD138pos | −0.0032 | 0.0304 | PASSED |
| MMRF_2170_1_BM_CD138pos | −0.0198 | 0.0164 | PASSED |
| MMRF_2089_3_BM_CD138pos | −0.024 | 0.0179 | PASSED |
| MMRF_2368_1_BM_CD138pos | −0.0222 | 0.0194 | PASSED |
| MMRF_1212_1_BM_CD138pos | 0.0016 | 0.0397 | FAILED |
| MMRF_2419_2_BM_CD138pos | −0.034 | 0.0295 | PASSED |
| MMRF_2047_1_BM_CD138pos | 0.0062 | 0.0412 | FAILED |
| MMRF_1038_1_BM_CD138pos | −0.0065 | 0.0275 | PASSED |
| MMRF_1596_1_BM_CD138pos | 0.0088 | 0.0437 | FAILED |
| MMRF_2230_1_BM_CD138pos | −0.0081 | 0.0261 | PASSED |
| MMRF_2480_1_BM_CD138pos | −0.0172 | 0.0158 | PASSED |
| MMRF_2455_1_BM_CD138pos | −0.0244 | 0.0326 | PASSED |
| MMRF_1671_4_BM_CD138pos | −0.0176 | 0.0153 | PASSED |
| MMRF_1490_1_BM_CD138pos | −0.0277 | 0.005 | PASSED |
| MMRF_1783_2_BM_CD138pos | −0.0158 | 0.0221 | PASSED |
| MMRF_1179_1_BM_CD138pos | −0.0296 | 0.0055 | PASSED |
| MMRF_1252_1_BM_CD138pos | 4e−04 | 0.0376 | FAILED |
| MMRF_1364_1_BM_CD138pos | −0.0113 | 0.0321 | PASSED |
| MMRF_2699_1_BM_CD138pos | −0.0082 | 0.0274 | PASSED |
| MMRF_2751_1_BM_CD138pos | −0.0093 | 0.0234 | PASSED |
| MMRF_2543_1_BM_CD138pos | −0.0207 | 0.0236 | PASSED |

### Exploratory PCA

Use this plot to see if samples are clustered according to the experimental design.

Use ARSyNseq function to correct potential batch effects.

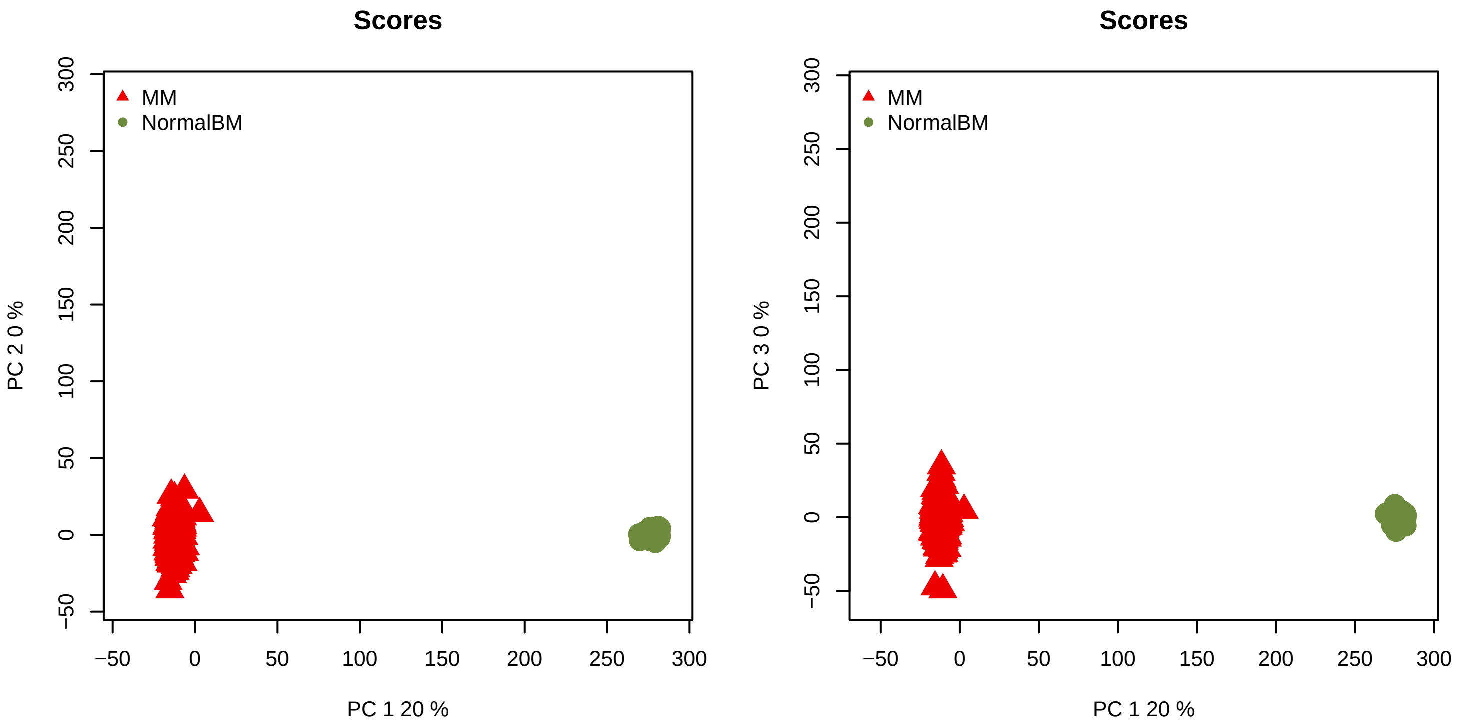
