## Supplementary material for "The network structure of hematopoietic cancers": Quality control for data pre-processing.: QCreport_MM_Before.pdf

### Quality Control of Expression Data

*Generated by NOISeq on 24 Feb 2023, 12:51:12*

#### Content

| *Plot* | *Description* |
|---|---|
| **Biotype detection** | Biotype abundance in the genome with %genes detected (counts > 0) in the sample/condition.<br>Biotype abundance within the sample/condition. |
| **Biotype expression** | Distribution of gene counts per million per biotype in sample/condition (only genes with counts > 0). |
| **Saturation** | Number of detected genes (counts > 0) per sample across different sequencing depths |
| **Expression boxplot** | Distribution of gene counts per million (all biotypes) in each sample/condition |
| **Expression barplot** | Percentage of genes with >0, >1, >2, >5 or >10 counts per million in each sample/condition. |
| **Length bias** | Mean gene expression per each length bin. Fitted curve and diagnostic test. |
| **GC content bias** | Mean gene expression per each GC content bin. Fitted curve and diagnostic test. |
| **RNA composition bias** | Density plots of log fold changes (M) between pairs of samples.<br>Confidence intervals for the median of M values. |
| **Exploratory PCA** | Principal Component Analysis score plots for PC1 vs PC2, and PC1 vs PC3. |

### Biotype detection

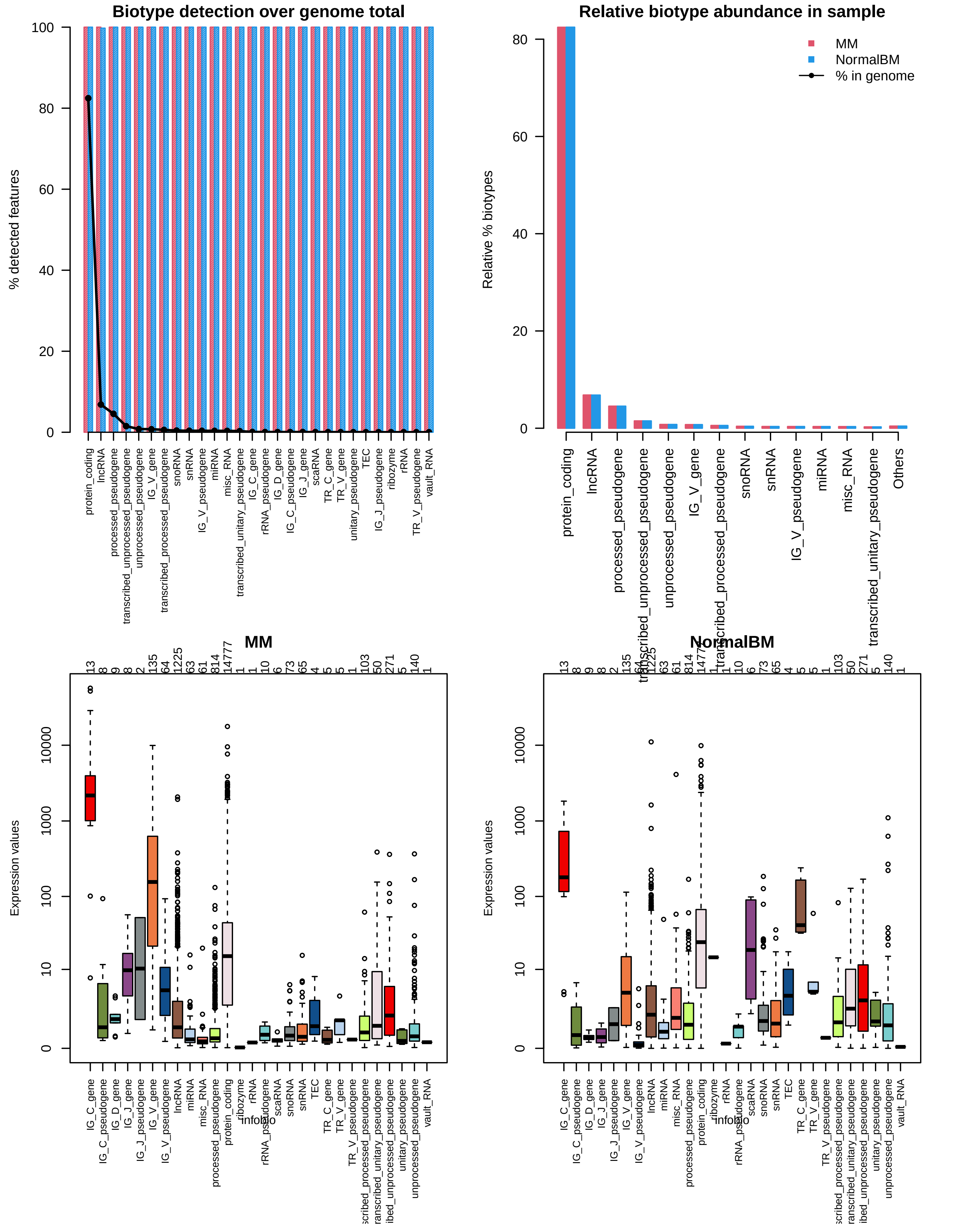

### Sequencing depth & Expression quantification

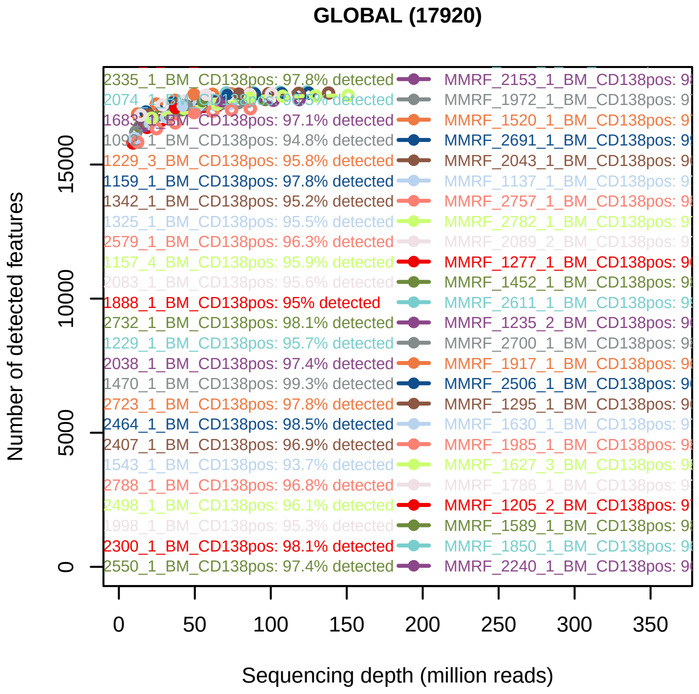


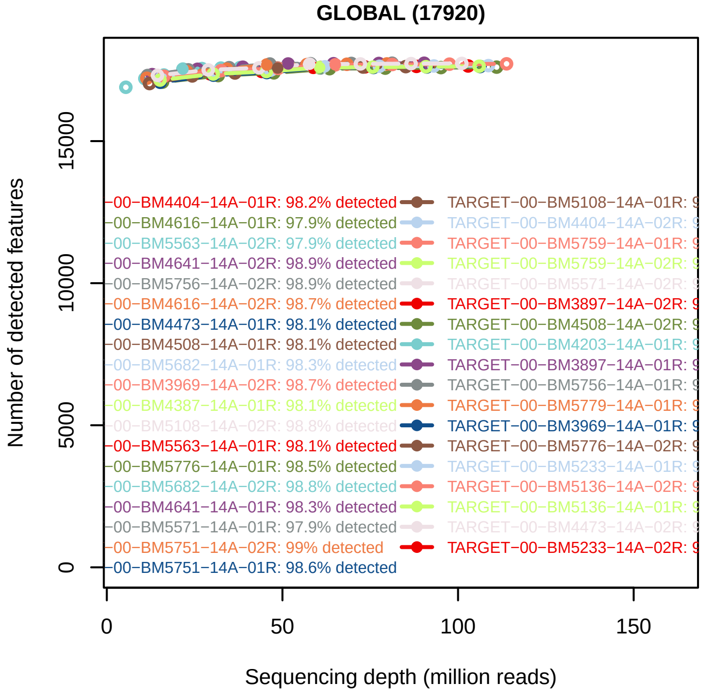


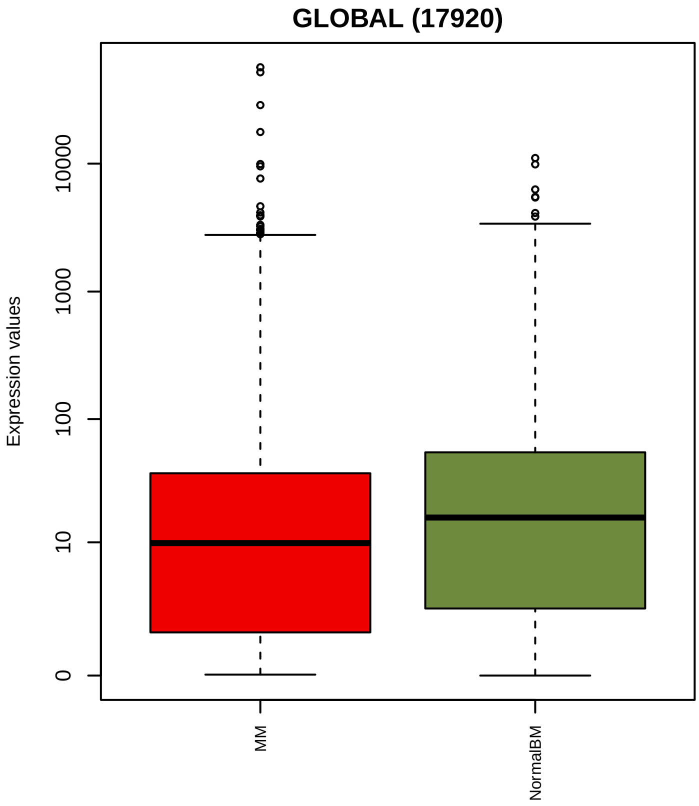


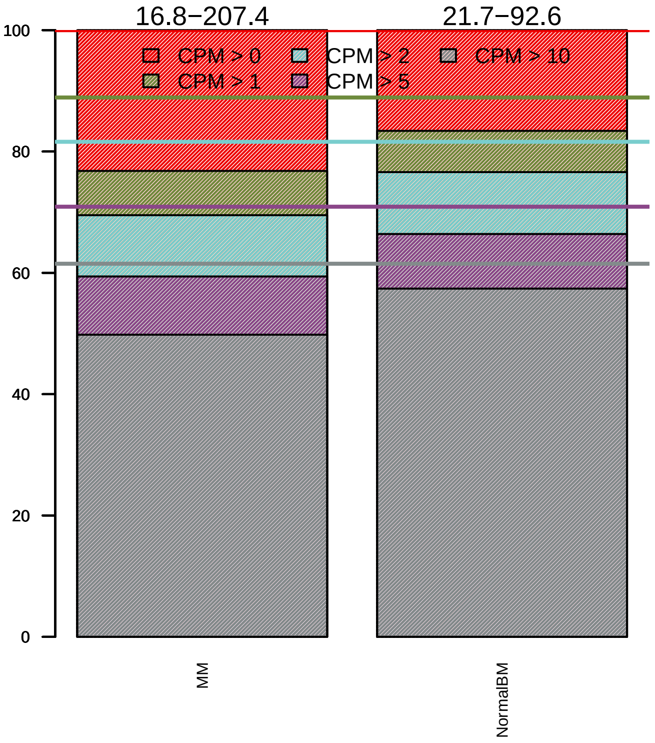

### Sequencing bias detection

#### *Diagnostic plot for feature length bias*

FAILED. At least one of the model p−values was lower than 0.05 and R2 > 70%.

Normalization for correcting length bias is recommended.

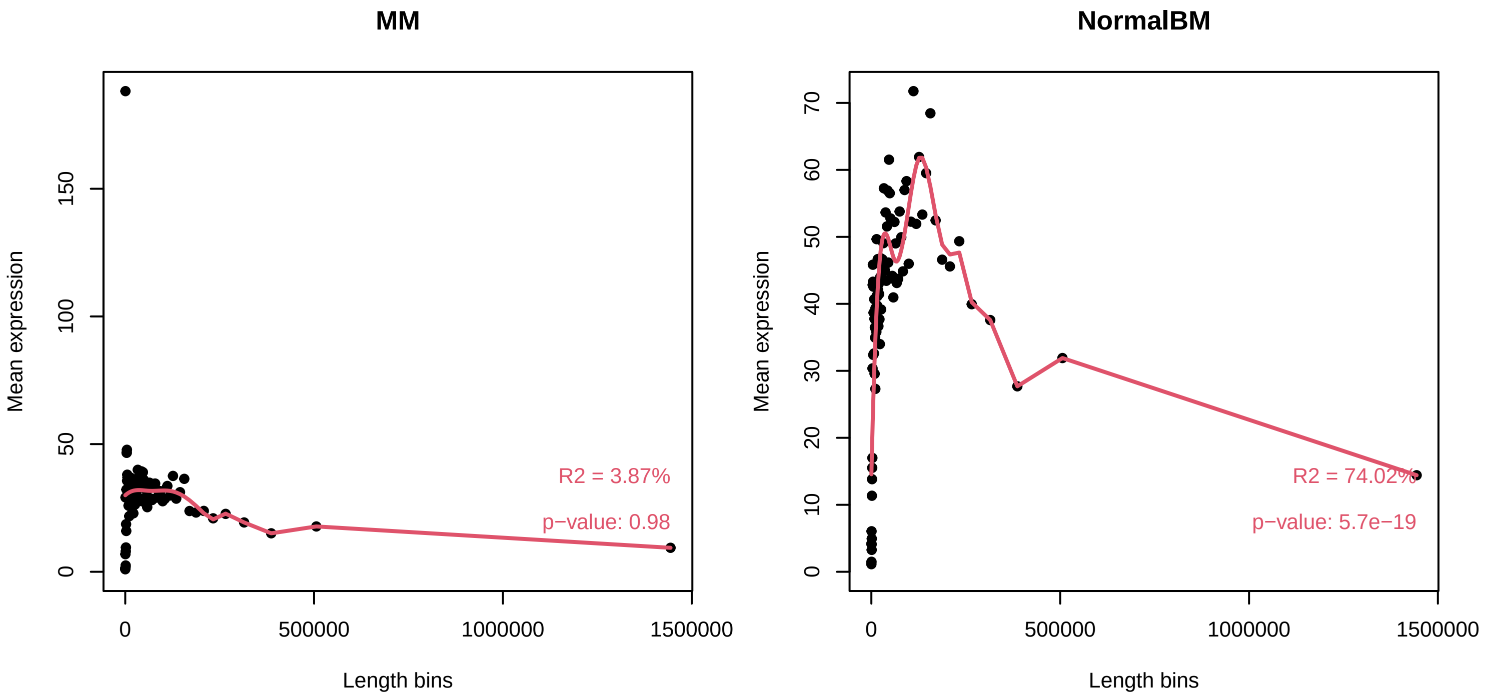


#### *Diagnostic plot for GC content bias*

WARNING. At least one of the model p−values was lower than 0.05, but R2 < 70% for at least one condition.

Normalization for correcting GC content bias could be advisable.
Plese check in the plots below the strength of the relationship between GC content and expression.

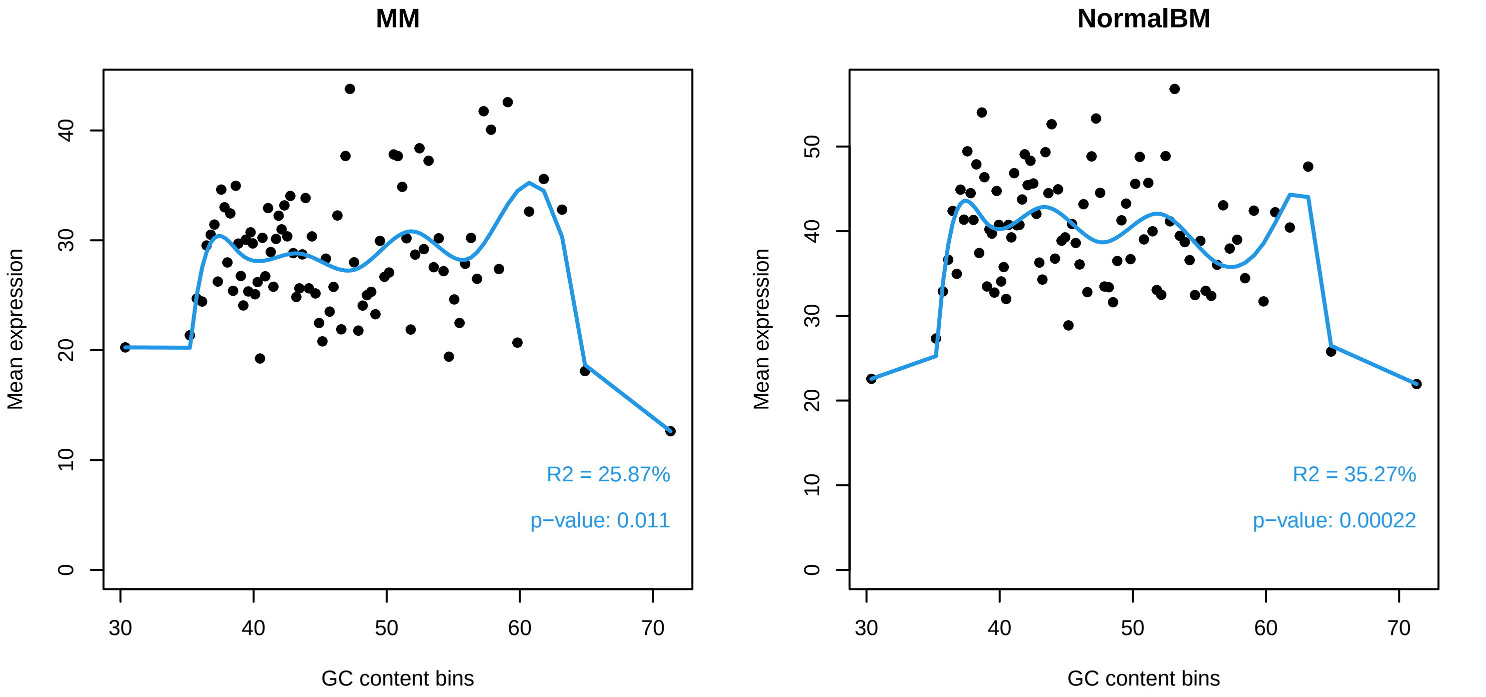

### *Diagnostic plot for differences in RNA composition*

FAILED. There is a pair of samples with significantly different RNA composition

Normalization for correcting this bias is required.

**Reference sample: MMRF_2173_1_BM_CD138pos**

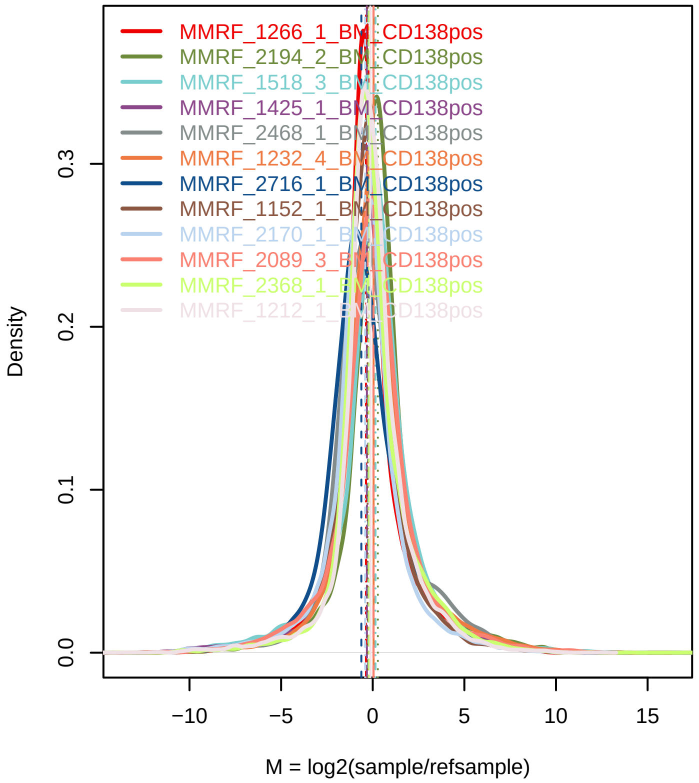


**Confidence intervals for median of M values**

| Sample | 0% | 100% | Diagnostic Test |
|---|---|---|---|
| MMRF_1266_1_BM_CD138pos | −0.406 | −0.3356 | FAILED |
| MMRF_2194_2_BM_CD138pos | 0.2472 | 0.3154 | FAILED |
| MMRF_1518_3_BM_CD138pos | 0.1315 | 0.1926 | FAILED |
| MMRF_1425_1_BM_CD138pos | −0.3653 | −0.275 | FAILED |
| MMRF_2468_1_BM_CD138pos | −0.3006 | −0.2189 | FAILED |
| MMRF_1232_4_BM_CD138pos | −0.0022 | 0.0568 | PASSED |
| MMRF_2716_1_BM_CD138pos | −0.6711 | −0.5658 | FAILED |
| MMRF_1152_1_BM_CD138pos | −0.3207 | −0.244 | FAILED |
| MMRF_2170_1_BM_CD138pos | −0.4433 | −0.4143 | FAILED |
| MMRF_2089_3_BM_CD138pos | 0.0026 | 0.0853 | FAILED |
| MMRF_2368_1_BM_CD138pos | −0.229 | −0.166 | FAILED |
| MMRF_1212_1_BM_CD138pos | −0.1606 | −0.0914 | FAILED |
| MMRF_2419_2_BM_CD138pos | −0.4434 | −0.3747 | FAILED |
| MMRF_2047_1_BM_CD138pos | −0.5089 | −0.4413 | FAILED |
| MMRF_1038_1_BM_CD138pos | −0.9969 | −0.9271 | FAILED |
| MMRF_1596_1_BM_CD138pos | 0.0117 | 0.0886 | FAILED |
| MMRF_2230_1_BM_CD138pos | −0.556 | −0.5087 | FAILED |
| MMRF_2480_1_BM_CD138pos | −0.1548 | −0.0846 | FAILED |
| MMRF_2455_1_BM_CD138pos | 0.0368 | 0.0945 | FAILED |
| MMRF_1671_4_BM_CD138pos | 0.1726 | 0.2205 | FAILED |
| MMRF_1490_1_BM_CD138pos | −0.1909 | −0.1168 | FAILED |
| MMRF_1783_2_BM_CD138pos | −0.2828 | −0.2207 | FAILED |
| MMRF_1179_1_BM_CD138pos | −0.3197 | −0.2448 | FAILED |
| MMRF_1252_1_BM_CD138pos | −0.3971 | −0.3245 | FAILED |
| MMRF_1364_1_BM_CD138pos | 0.1336 | 0.2142 | FAILED |
| MMRF_2699_1_BM_CD138pos | −0.3253 | −0.22 | FAILED |
| MMRF_2751_1_BM_CD138pos | −0.2446 | −0.1959 | FAILED |
| MMRF_2543_1_BM_CD138pos | −0.5376 | −0.4904 | FAILED |

### Exploratory PCA

Use this plot to see if samples are clustered according to the experimental design.

Use ARSyNseq function to correct potential batch effects.

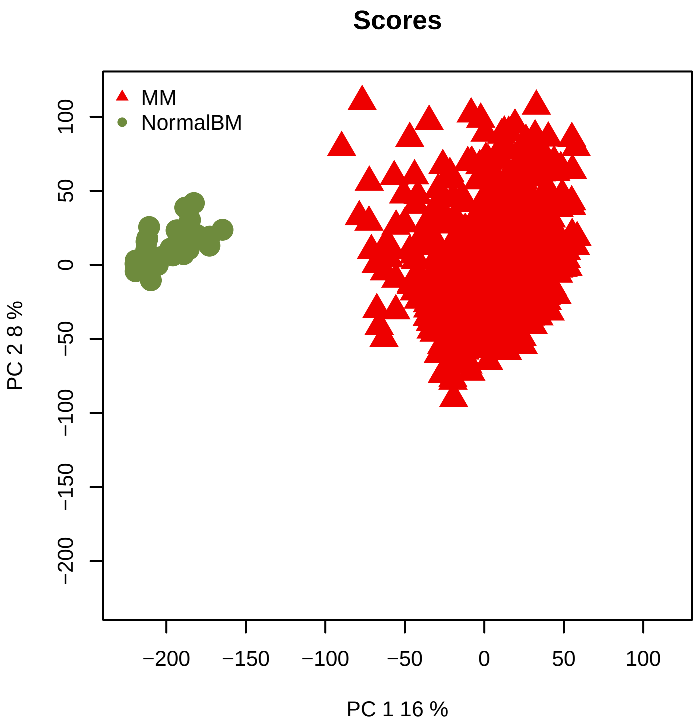


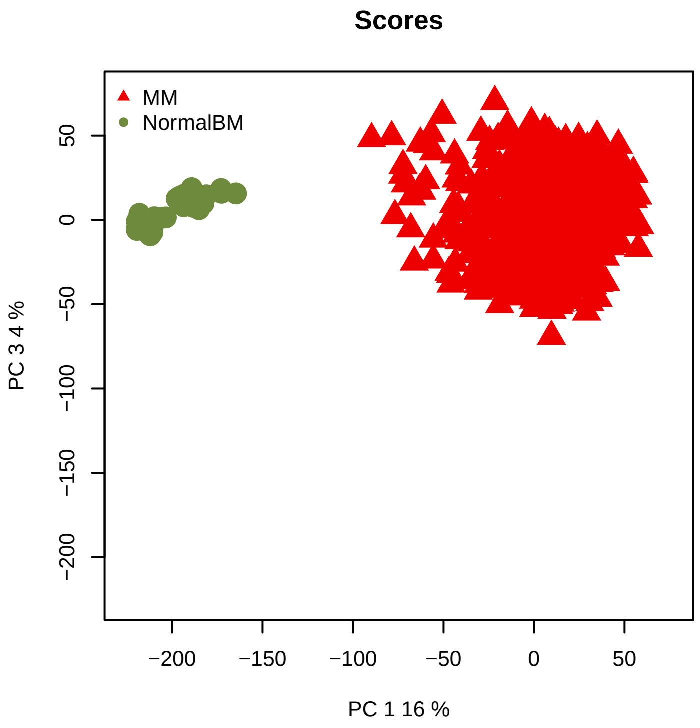
