## Supplementary material for "The network structure of hematopoietic cancers": Quality control for data pre-processing.: QCreport_TALL_After.pdf

### Quality Control of Expression Data

*Generated by NOISeq on 24 Feb 2023, 13:34:38*

#### Content

| *Plot* | *Description* |
|---|---|
| **Biotype detection** | Biotype abundance in the genome with %genes detected (counts > 0) in the sample/condition.<br>Biotype abundance within the sample/condition. |
| **Biotype expression** | Distribution of gene counts per million per biotype in sample/condition (only genes with counts > 0). |
| **Saturation** | Number of detected genes (counts > 0) per sample across different sequencing depths |
| **Expression boxplot** | Distribution of gene counts per million (all biotypes) in each sample/condition |
| **Expression barplot** | Percentage of genes with >0, >1, >2, >5 or >10 counts per million in each sample/condition. |
| **Length bias** | Mean gene expression per each length bin. Fitted curve and diagnostic test. |
| **GC content bias** | Mean gene expression per each GC content bin. Fitted curve and diagnostic test. |
| **RNA composition bias** | Density plots of log fold changes (M) between pairs of samples.<br>Confidence intervals for the median of M values. |
| **Exploratory PCA** | Principal Component Analysis score plots for PC1 vs PC2, and PC1 vs PC3. |

### Biotype detection

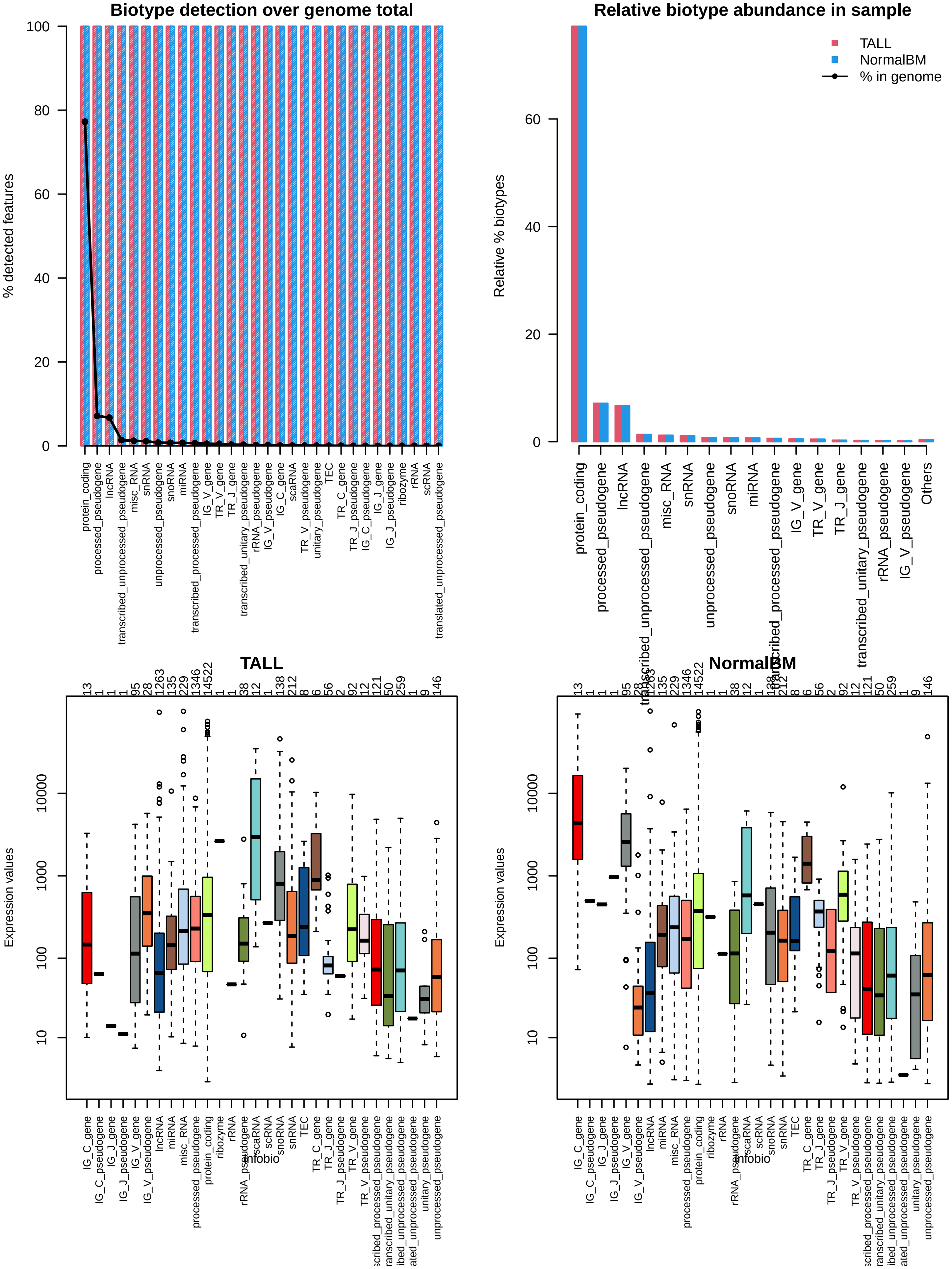

### Sequencing depth & Expression quantification

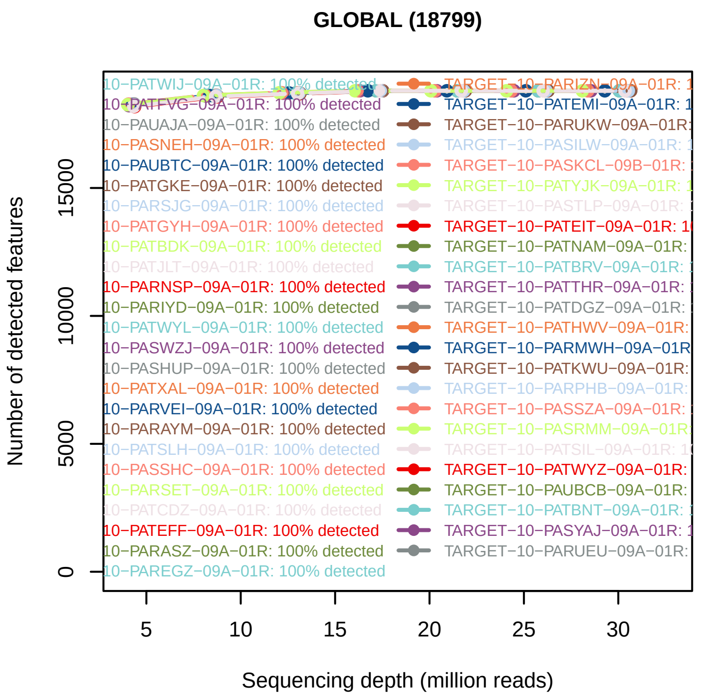


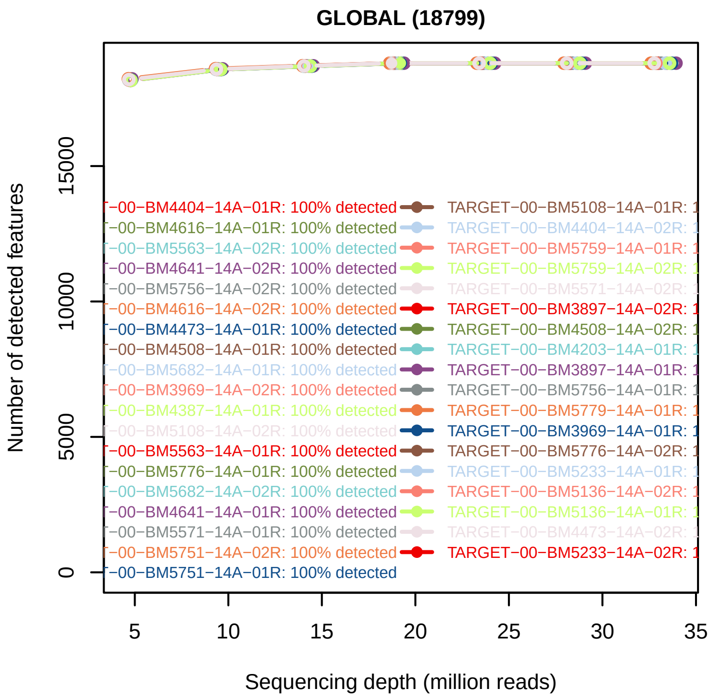


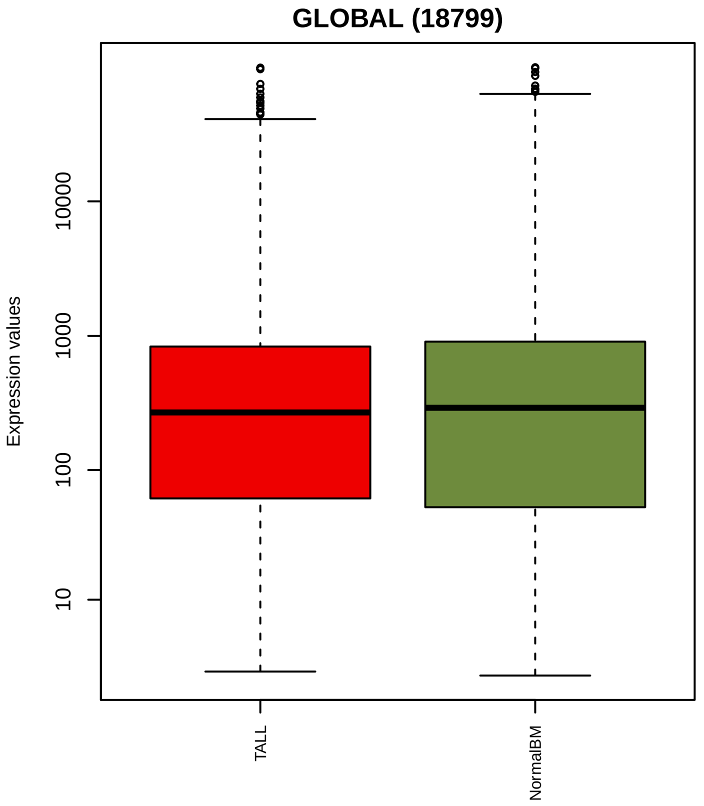


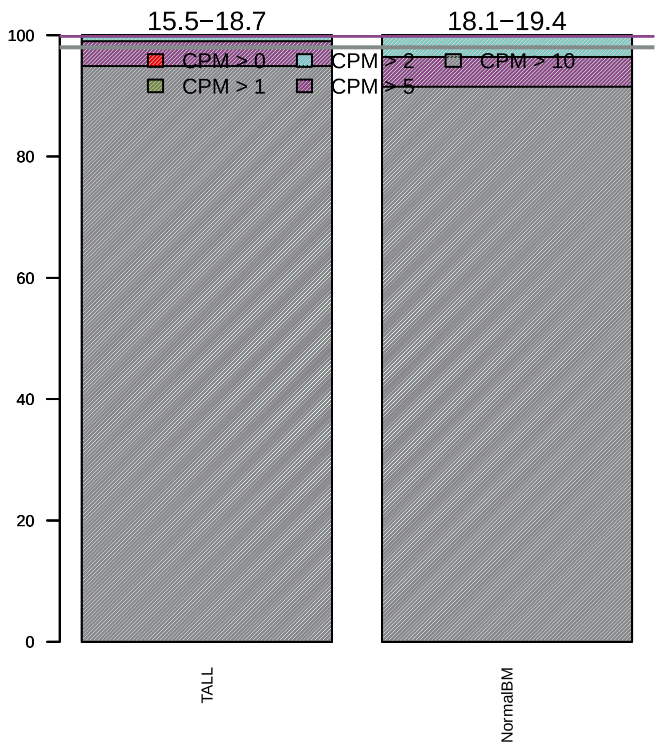

### Sequencing bias detection

#### *Diagnostic plot for feature length bias*

WARNING. At least one of the model p−values was lower than 0.05, but R2 < 70% for at least one condition.

Normalization for correcting length bias could be advisable.
Plese check in the plots below the strength of the relationship between length and expression.

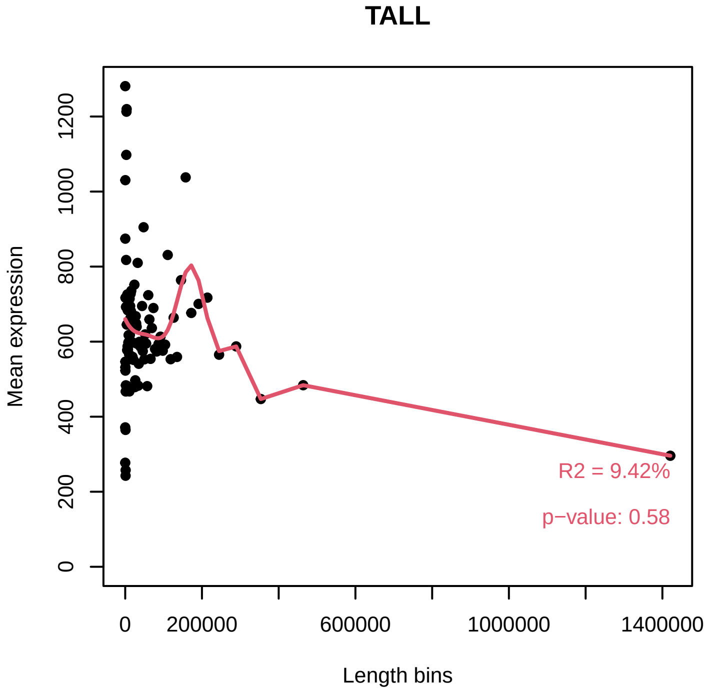


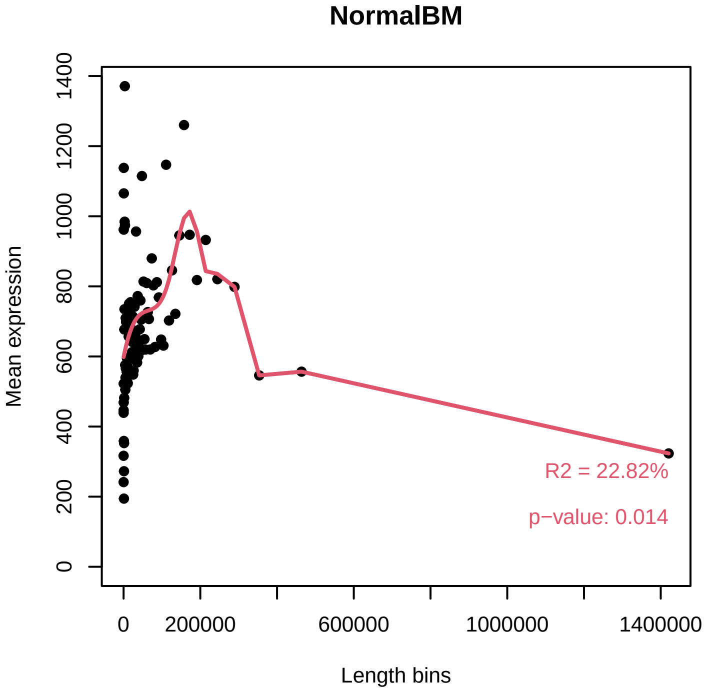


#### *Diagnostic plot for GC content bias*

PASSED. No normalization for correcting GC content bias is required.

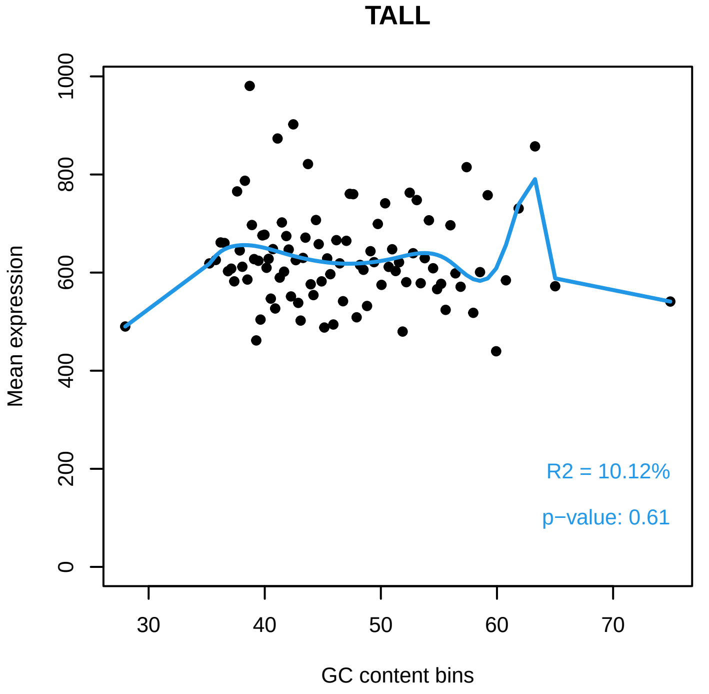


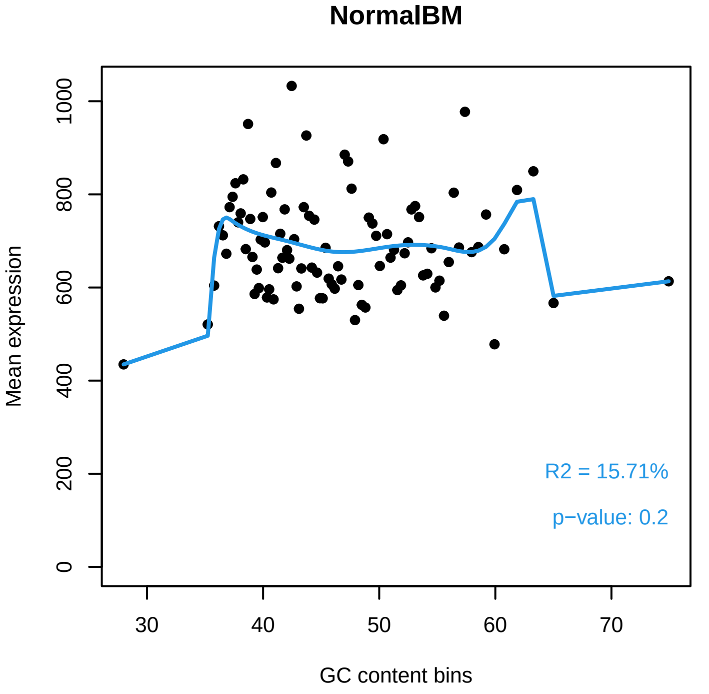

#### *Diagnostic plot for differences in RNA composition*

FAILED. There is a pair of samples with significantly different RNA composition

Normalization for correcting this bias is required.

##### Reference sample: TARGET−10−PATHGY−09A−01R

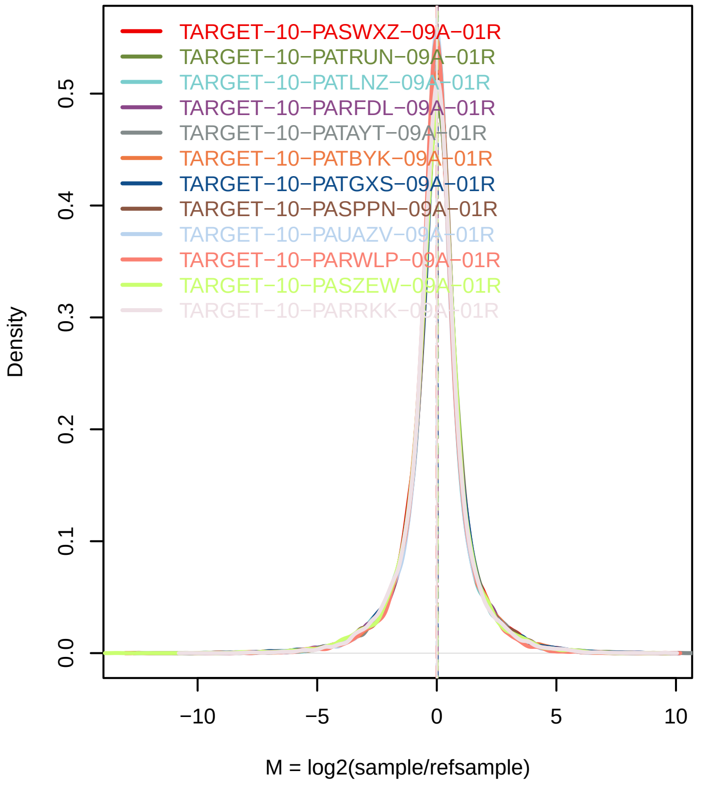


##### Confidence intervals for median of M values

| Sample | 0.01% | 99.99% | Diagnostic Test |
|---|---|---|---|
| TARGET−10−PASWXZ−09A−01R | [illegible] | 0.0444 | FAILED |
| TARGET−10−PATRUN−09A−01R | 0.0107 | 0.0557 | FAILED |
| TARGET−10−PATLNZ−09A−01R | −0.0182 | 0.0236 | PASSED |
| TARGET−10−PARFDL−09A−01R | 0.0084 | 0.0577 | FAILED |
| TARGET−10−PATAYT−09A−01R | 0.0154 | 0.0518 | FAILED |
| TARGET−10−PATBYK−09A−01R | −0.0093 | 0.0329 | PASSED |
| TARGET−10−PATGXS−09A−01R | 0.0129 | 0.0552 | FAILED |
| TARGET−10−PASPPN−09A−01R | 0.0003 | 0.0521 | FAILED |
| TARGET−10−PAUAZV−09A−01R | −0.0294 | 0.0179 | PASSED |
| TARGET−10−PARWLP−09A−01R | −0.0293 | 0.0126 | PASSED |
| TARGET−10−PASZEW−09A−01R | 7e−04 | 0.0489 | FAILED |
| TARGET−10−PARRKK−09A−01R | −0.0291 | 0.0231 | PASSED |
| TARGET−10−PAUAYB−09A−01R | −0.0206 | 0.0171 | PASSED |
| TARGET−10−PASVPZ−09A−01R | 0.0104 | 0.0515 | FAILED |
| TARGET−10−PATNIA−09A−01R | −0.0068 | 0.0405 | PASSED |
| TARGET−10−PASXMF−09A−01R | 0.0228 | 0.0571 | FAILED |
| TARGET−10−PASKXN−09A−01R | −0.0187 | 0.0261 | PASSED |
| TARGET−10−PASPBU−09A−01R | −0.0009 | 0.0372 | PASSED |
| TARGET−10−PARGFL−09A−01R | −0.0291 | 0.0084 | PASSED |
| TARGET−10−PATIBE−09A−01R | 0.0129 | 0.0581 | FAILED |
| TARGET−10−PAUACG−09A−01R | 0.0144 | 0.0634 | FAILED |
| TARGET−10−PATHJF−09A−01R | −0.0154 | 0.0325 | PASSED |
| TARGET−10−PASLBB−09A−01R | −0.0151 | 0.0295 | PASSED |
| TARGET−10−PATKYH−09A−01R | −0.0067 | 0.0457 | PASSED |
| TARGET−10−PASSRP−09A−01R | 0.0072 | 0.0505 | FAILED |
| TARGET−10−PASUIN−09A−01R | −0.0112 | 0.0296 | PASSED |
| TARGET−10−PASHXL−09A−01R | −0.0119 | 0.0248 | PASSED |
| TARGET−10−PARNMV−09A−01R | 0.0012 | 0.0464 | FAILED |

### Exploratory PCA

Use this plot to see if samples are clustered according to the experimental design.

Use ARSyNseq function to correct potential batch effects.

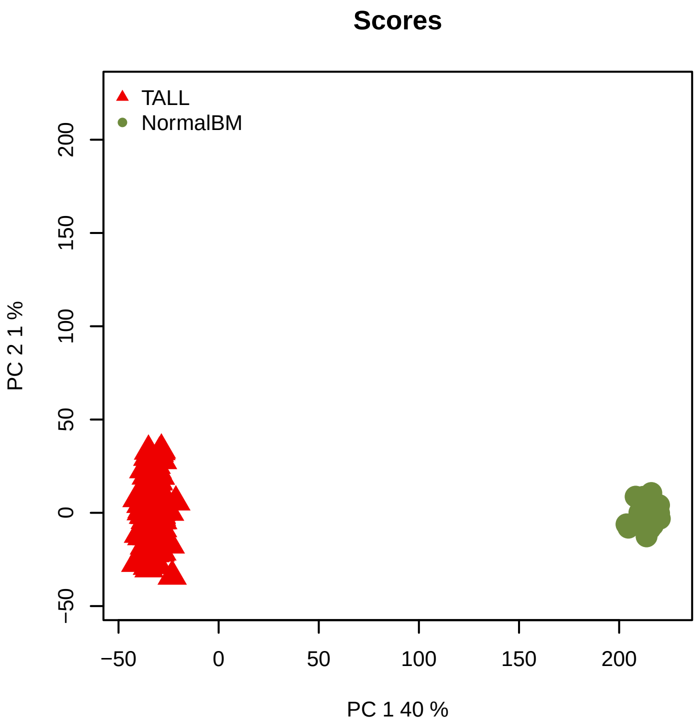


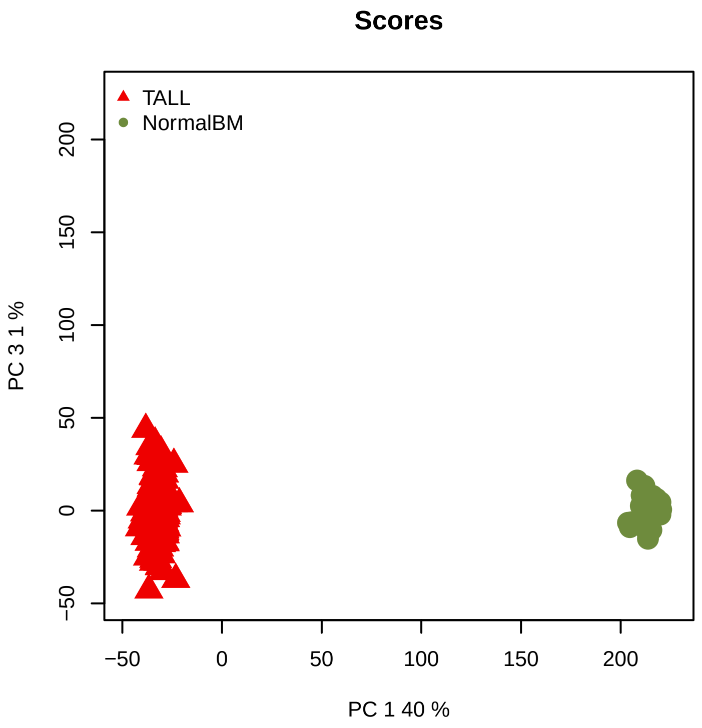
