## Supplementary material for "The network structure of hematopoietic cancers": Quality control for data pre-processing.: QCreport_TALL_Before.pdf

### Quality Control of Expression Data

*Generated by NOISeq on 24 Feb 2023, 13:05:02*

#### Content

| *Plot* | *Description* |
|---|---|
| **Biotype detection** | Biotype abundance in the genome with %genes detected (counts > 0) in the sample/condition.<br>Biotype abundance within the sample/condition. |
| **Biotype expression** | Distribution of gene counts per million per biotype in sample/condition (only genes with counts > 0). |
| **Saturation** | Number of detected genes (counts > 0) per sample across different sequencing depths |
| **Expression boxplot** | Distribution of gene counts per million (all biotypes) in each sample/condition |
| **Expression barplot** | Percentage of genes with >0, >1, >2, >5 or >10 counts per million in each sample/condition. |
| **Length bias** | Mean gene expression per each length bin. Fitted curve and diagnostic test. |
| **GC content bias** | Mean gene expression per each GC content bin. Fitted curve and diagnostic test. |
| **RNA composition bias** | Density plots of log fold changes (M) between pairs of samples.<br>Confidence intervals for the median of M values. |
| **Exploratory PCA** | Principal Component Analysis score plots for PC1 vs PC2, and PC1 vs PC3. |

### Biotype detection

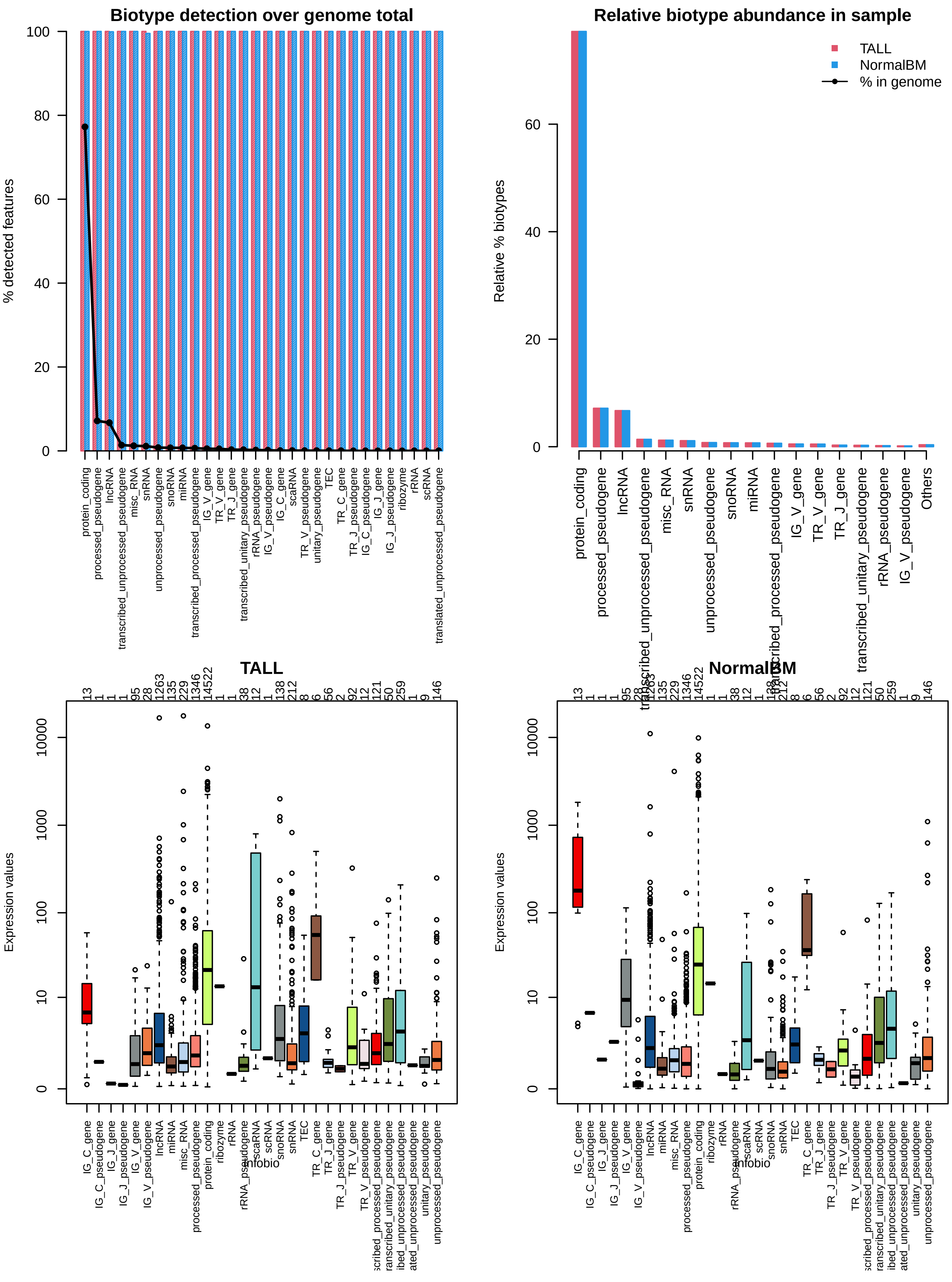

### Sequencing depth & Expression quantification

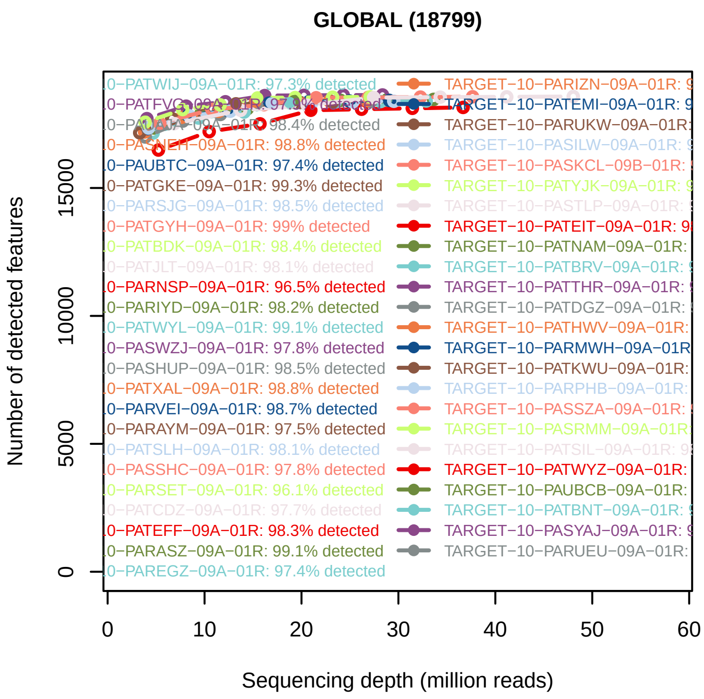


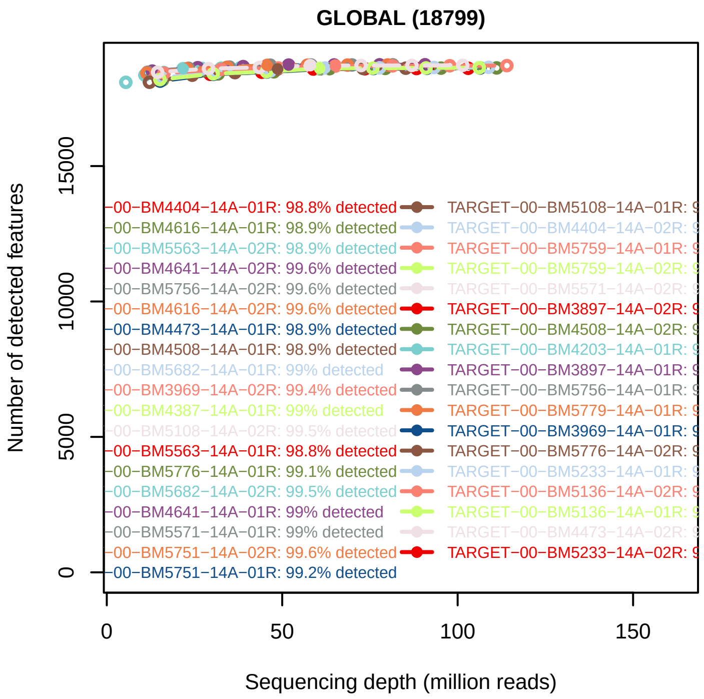


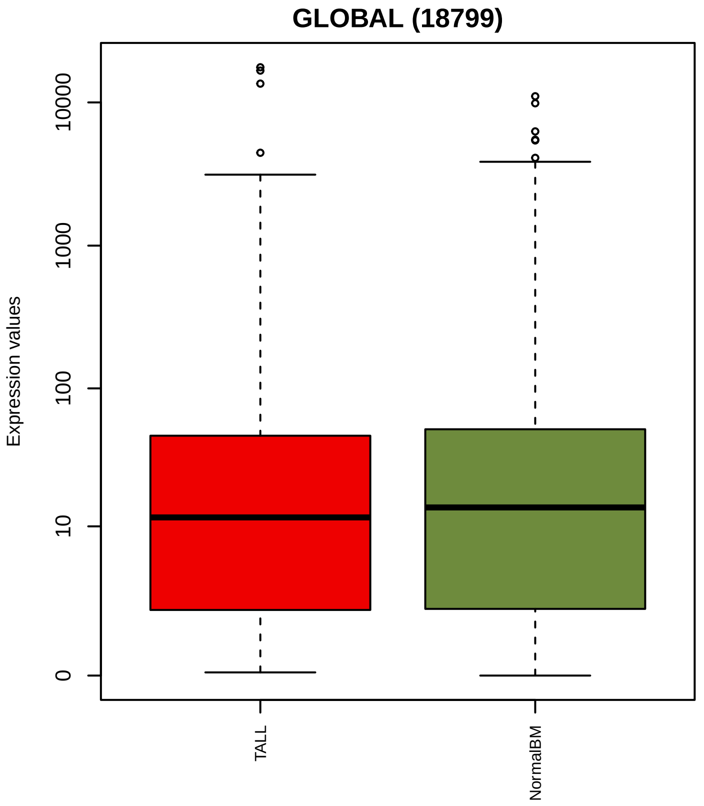


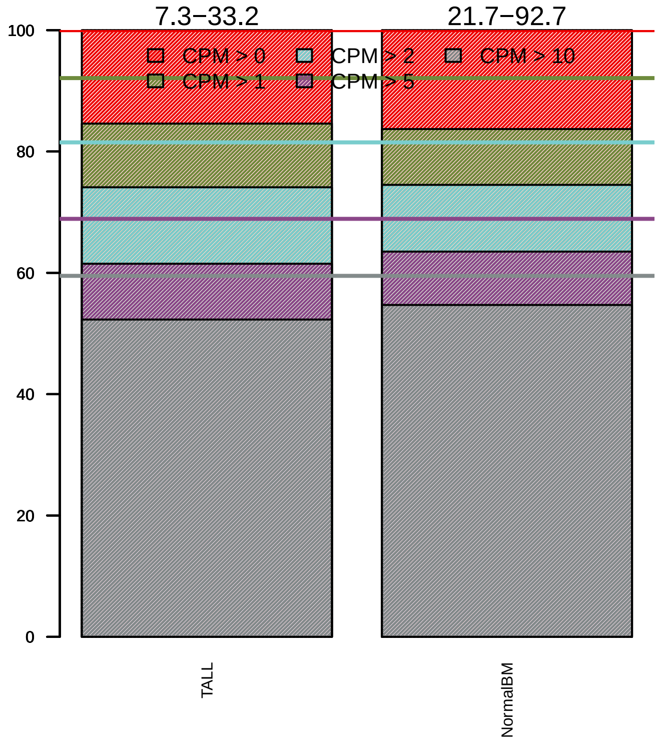

### Sequencing bias detection

#### *Diagnostic plot for feature length bias*

FAILED. At least one of the model p−values was lower than 0.05 and R2 > 70%.

Normalization for correcting length bias is recommended.

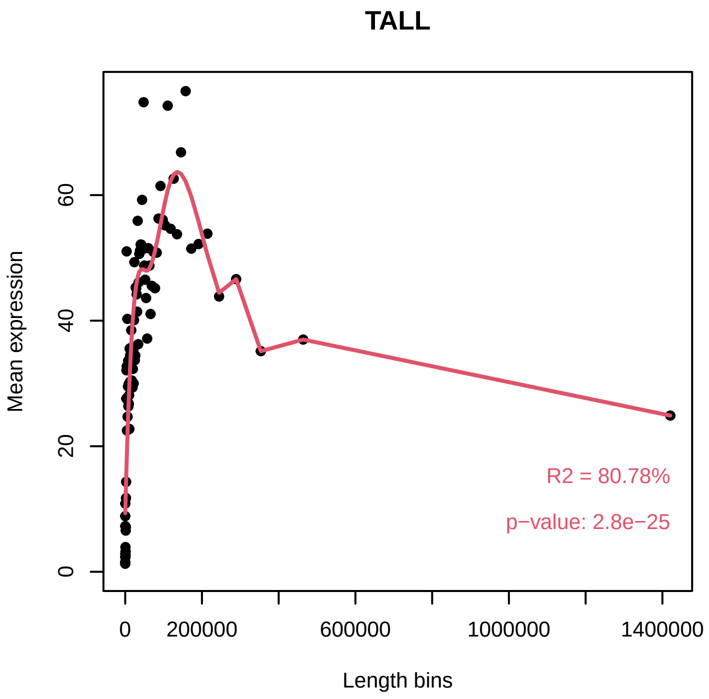


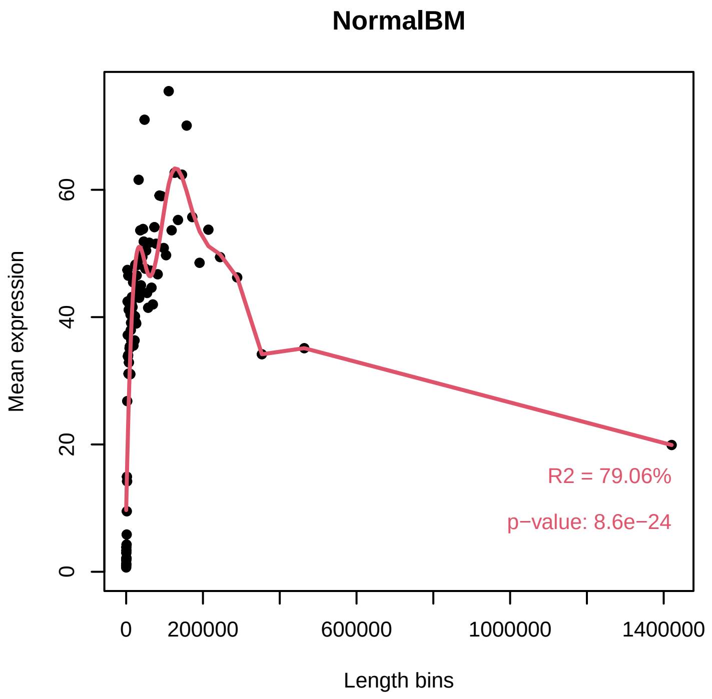


#### *Diagnostic plot for GC content bias*

FAILED. At least one of the model p−values was lower than 0.05 and R2 > 70%.

Normalization for correcting GC content bias is recommended.

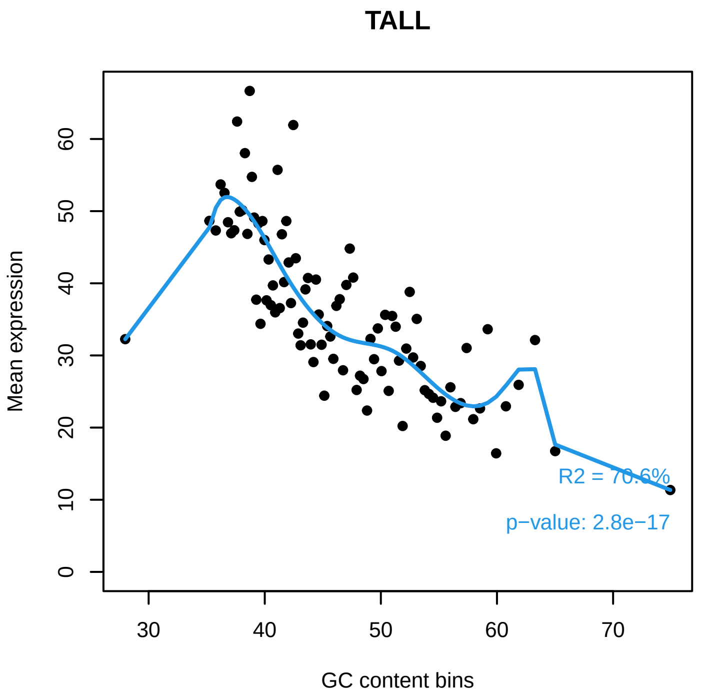


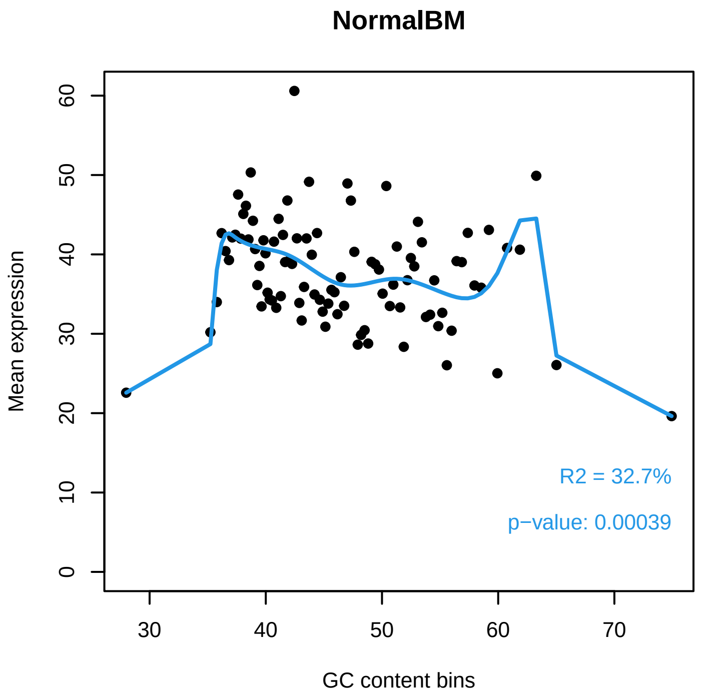

### *Diagnostic plot for differences in RNA composition*

FAILED. There is a pair of samples with significantly different RNA composition

Normalization for correcting this bias is required.

**Reference sample: TARGET−10−PATHGY−09A−01R**

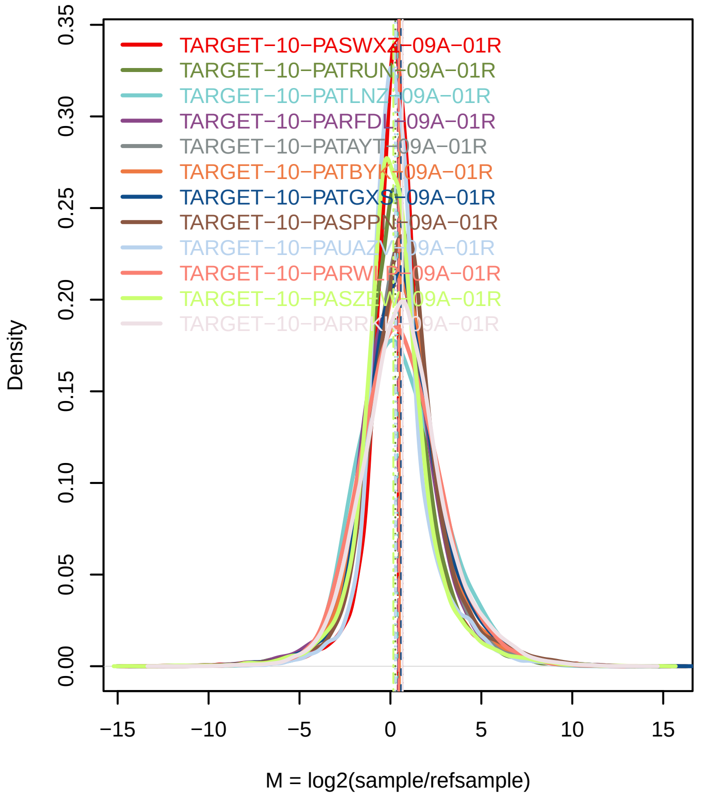


**Confidence intervals for median of M values**

| Sample | 0.01% | 99.99% | Diagnostic Test |
|---|---|---|---|
| TARGET−10−PASWX0.2379A−01R | | 0.4469 | FAILED |
| TARGET−10−PATRUN0.2914−01R | | 0.3187 | FAILED |
| TARGET−10−PATLNZ0.3712−01R | | 0.4875 | FAILED |
| TARGET−10−PARFDL0.3495−01R | | 0.4342 | FAILED |
| TARGET−10−PATAYT0.4577−01R | | 0.4952 | FAILED |
| TARGET−10−PATBYK−0.4957−01R | | 0.5667 | FAILED |
| TARGET−10−PATGXS−0.5923−01R | | 0.6546 | FAILED |
| TARGET−10−PASPPN0.6252−01R | | 0.7224 | FAILED |
| TARGET−10−PAUAZV0.2194−01R | | 0.2843 | FAILED |
| TARGET−10−PARWLP0.4396−01R | | 0.5314 | FAILED |
| TARGET−10−PASZEW0.1094−01R | | 0.208 | FAILED |
| TARGET−10−PARRKK0.6094−01R | | 0.7236 | FAILED |
| TARGET−10−PAUAYB−0.0976−01R | | −0.0232 | FAILED |
| TARGET−10−PASVPZ0.2552−01R | | 0.3595 | FAILED |
| TARGET−10−PATNIA−0.1397−01R | | −0.0654 | FAILED |
| TARGET−10−PASXMF0.1669−01R | | 0.2918 | FAILED |
| TARGET−10−PASKXN−0.017−01R | | 0.1312 | PASSED |
| TARGET−10−PASPBU0.2168−01R | | 0.2937 | FAILED |
| TARGET−10−PARGFL−0.2396−01R | | −0.1504 | FAILED |
| TARGET−10−PATIBE0.4994−01R | | 0.5799 | FAILED |
| TARGET−10−PAUACG0.6512−01R | | 0.7659 | FAILED |
| TARGET−10−PATHJF−0.937−01R | | 0.496 | FAILED |
| TARGET−10−PASLBB0.5301−01R | | 0.6507 | FAILED |
| TARGET−10−PATKYI0.2905−01R | | 0.3987 | FAILED |
| TARGET−10−PASSRP0.1291−01R | | 0.2257 | FAILED |
| TARGET−10−PASUIN0.2799−01R | | 0.3532 | FAILED |
| TARGET−10−PASHXL0.0747−01R | | 0.1342 | FAILED |
| TARGET−10−PARNMV.5093−01R | | 0.5792 | FAILED |

### Exploratory PCA

Use this plot to see if samples are clustered according to the experimental design.

Use ARSyNseq function to correct potential batch effects.

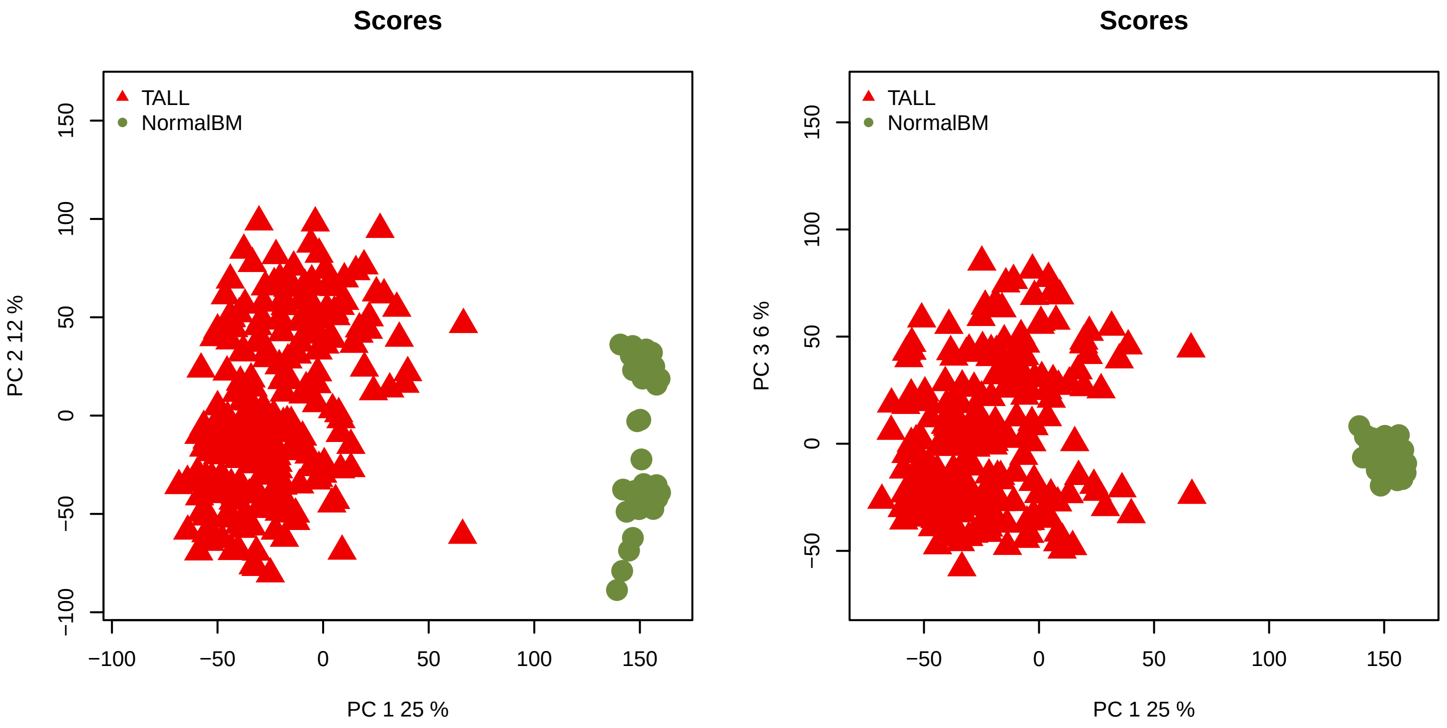
